# The landscape of tolerated genetic variation in humans and primates

**DOI:** 10.1101/2023.05.01.538953

**Authors:** Hong Gao, Tobias Hamp, Jeffrey Ede, Joshua G. Schraiber, Jeremy McRae, Moriel Singer-Berk, Yanshen Yang, Anastasia Dietrich, Petko Fiziev, Lukas Kuderna, Laksshman Sundaram, Yibing Wu, Aashish Adhikari, Yair Field, Chen Chen, Serafim Batzoglou, Francois Aguet, Gabrielle Lemire, Rebecca Reimers, Daniel Balick, Mareike C. Janiak, Martin Kuhlwilm, Joseph D. Orkin, Shivakumara Manu, Alejandro Valenzuela, Juraj Bergman, Marjolaine Rouselle, Felipe Ennes Silva, Lidia Agueda, Julie Blanc, Marta Gut, Dorien de Vries, Ian Goodhead, R. Alan Harris, Muthuswamy Raveendran, Axel Jensen, Idriss S. Chuma, Julie Horvath, Christina Hvilsom, David Juan, Peter Frandsen, Fabiano R. de Melo, Fabricio Bertuol, Hazel Byrne, Iracilda Sampaio, Izeni Farias, João Valsecchi do Amaral, Mariluce Messias, Maria N. F. da Silva, Mihir Trivedi, Rogerio Rossi, Tomas Hrbek, Nicole Andriaholinirina, Clément J. Rabarivola, Alphonse Zaramody, Clifford J. Jolly, Jane Phillips-Conroy, Gregory Wilkerson, Christian Abee, Joe H. Simmons, Eduardo Fernandez-Duque, ee Kanthaswamy, Fekadu Shiferaw, Dongdong Wu, Long Zhou, Yong Shao, Guojie Zhang, Julius D. Keyyu, Sascha Knauf, Minh D. Le, Esther Lizano, Stefan Merker, Arcadi Navarro, Thomas Batallion, Tilo Nadler, Chiea Chuen Khor, Jessica Lee, Patrick Tan, Weng Khong Lim, Andrew C. Kitchener, Dietmar Zinner, Ivo Gut, Amanda Melin, Katerina Guschanski, Mikkel Heide Schierup, Robin M. D. Beck, Govindhaswamy Umapathy, Christian Roos, Jean P. Boubli, Monkol Lek, Shamil Sunyaev, Anne O’Donnell, Heidi Rehm, Jinbo Xu, Jeffrey Rogers, Tomas Marques-Bonet, Kyle Kai-How Farh

## Abstract

Personalized genome sequencing has revealed millions of genetic differences between individuals, but our understanding of their clinical relevance remains largely incomplete. To systematically decipher the effects of human genetic variants, we obtained whole genome sequencing data for 809 individuals from 233 primate species, and identified 4.3 million common protein-altering variants with orthologs in human. We show that these variants can be inferred to have non-deleterious effects in human based on their presence at high allele frequencies in other primate populations. We use this resource to classify 6% of all possible human protein-altering variants as likely benign and impute the pathogenicity of the remaining 94% of variants with deep learning, achieving state-of-the-art accuracy for diagnosing pathogenic variants in patients with genetic diseases.

**One Sentence Summary:** Deep learning classifier trained on 4.3 million common primate missense variants predicts variant pathogenicity in humans.

## Main Text

A scalable approach for interpreting the effects of human genetic variants and their impact on disease risk is urgently needed to realize the promise of personalized genomic medicine (*1–3*). Out of more than 70 million possible protein-altering variants in the human genome, only ∼0.1% are annotated in clinical variant databases such as ClinVar (*4*), with the remainder being variants of uncertain clinical significance (*5, 6*). Despite collaborative efforts by the scientific community, the rarity of most human genetic variants has meant that progress towards deciphering personal genomes has been incremental (*7, 8*). Consequently, clinical sequencing tests frequently return without definitive diagnoses, a frustrating outcome for both patients and clinicians (*9, 10*). In certain cases, patients have needed to be recontacted and diagnoses reversed when the presumed pathogenic variant was later found to be a common variant in previously understudied human populations (*11–13*). Common variants can often be ruled out as the cause of penetrant genetic disease, since their high frequency in the population indicates that they are tolerated by natural selection, aside from rare exceptions due to founder effects and balancing selection (*14–16*).

An emerging strategy for solving clinical variant interpretation on a genome-wide scale is the use of information from closely related primate species to infer the pathogenicity of orthologous human variants (*17*). Because chimpanzees and humans share 99.4% protein sequence identity (*18*), a protein-altering variant present in one species can be expected to produce similar effects on the protein in the other species. By conducting population sequencing studies in closely related non-human primate species, it is feasible to systematically catalog common variants and rule these out as pathogenic in human, analogous to how sequencing more diverse human populations has helped to advance clinical variant interpretation (*8, 17*). Nonetheless, earlier work (*17*) was limited by the very small primate population sequencing datasets available, which bounded the number of common variants discovered, and the scale of machine learning classifiers that could be trained.

## RESULTS

### A database of 4.3 million benign missense variants across the primate lineage

To expand upon this strategy, we sequenced 703 individuals from 211 primate species (*19*), and aggregated these with data from previous studies (*19–26*), yielding a total of 809 individuals from 233 species. We identified 4.3 million unique missense (protein-altering) variants and 6.7 million unique synonymous (non-protein altering) variants (Fig. 1A), after excluding variants at positions that lacked unambiguous 1:1 mapping with human, or which resulted in non-concordant amino acid translation outcomes because of changes at neighboring nucleotides (fig. S1). The species selected for sequencing represent close to half of the 521 extant primate species on Earth (*27*) and cover all major primate families, from Old World monkeys and New World monkeys to lemurs and tarsiers. We targeted a small number of individuals per species (3.5 on average) to ensure that we primarily sampled common variants that have been filtered by natural selection rather than rare mutations (fig. S2).

**Fig. 1.**
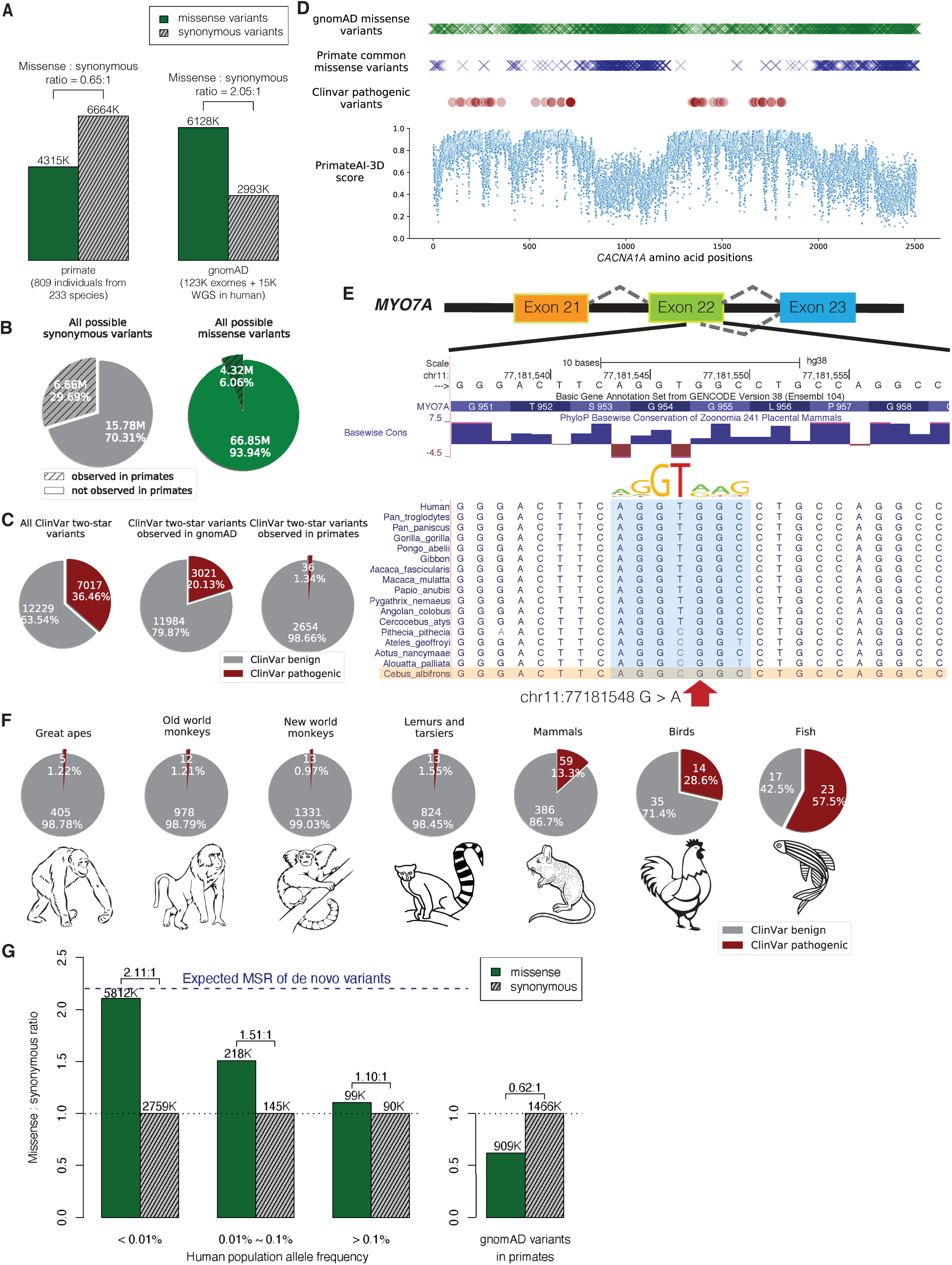
Common primate variants are largely benign in human. (**A**) Counts of missense (solid green) and synonymous (shaded grey) variants from primates compared to the gnomAD database. Missense : synonymous counts and ratios are displayed above each bar. (**B**) Fractions of all possible human synonymous (grey) and missense variants (green) observed in primates. (**C**) Counts of benign (grey) and pathogenic (red) missense variants with two-star review status or above in the overall ClinVar database (left pie chart), compared to ClinVar variants observed in gnomAD (middle), and compared to ClinVar variants observed in primates (right). Conflicting benign and pathogenic annotations and variants interpreted only with uncertain significance were excluded. (**D**) Observed gnomAD (green) or primate (blue) missense variants in each amino acid position in the *CACNA1A* gene. Red circles represent the positions of annotated ClinVar pathogenic missense variants. Bottom scatterplot shows PrimateAI-3D predicted pathogenicity scores for all possible missense substitutions along the gene. (**E**) Multiple sequence alignment showing the ClinVar pathogenic variant chr11:77181548 G>A (red arrow) creating a cryptic splice site in human sequence (extended splice motif, blue). This variant is tolerated in Cebus Albifrons and other species with a G>C synonymous change in the adjacent nucleotide that stops the splice motif from forming. (**F**) Pie charts showing the fraction of benign (grey) and pathogenic (red) missense variants with ClinVar two-star review status or above in great apes, old world monkeys, new world monkeys, lemurs/tarsiers, mammals, chicken, and zebrafish. (**G**) Missense : synonymous ratios across the human allele frequency spectrum, with MSR of human variants seen in primates shown for comparison. The blue dashed line represents the expected missense : synonymous ratio of de novo variants. Colors and legend are the same as (**A**).

Compared to the genome Aggregation Database (gnomAD) cohort of 141,456 human individuals from diverse populations (*28, 29*), the primate sequencing cohort contained ∼20% more exome variants despite sequencing 1/175th the number of individuals (Fig. 1A and fig. S3), attesting to the remarkable genetic diversity present in non-human primate species (*19, 30*), many of which are critically endangered (*31*). The overlap of primate variants with gnomAD was low, consistent with independent mutational origins in each species (fig. S3). Out of the 22 million possible synonymous variants in the human genome, 30% were observed in the primate cohort, compared to just 6% of possible missense mutations (Fig. 1B). Because de novo mutations would have laid down unbiased proportions of missense and synonymous variants, the observed depletion of missense mutations in the primate cohort is consistent with the majority of newly-arising human missense mutations being removed by natural selection due to their deleteriousness (*8, 32–34*).

The surviving missense variants are seen at high frequencies in primate populations, and represent a subset of missense variants that have tolerated filtering by natural selection and are unlikely to be pathogenic (*35*).

Missense variants from the primate cohort are strongly enriched for benign consequence in the ClinVar clinical variant database (Fig. 1C). Amongst ClinVar variants with higher review levels (2-star and above, indicating consensus by multiple submitters) (*4*), missense variants found in at least one non-human primate species were Benign or Likely Benign ∼99% of the time, compared to 63% for ClinVar missense variants in general, and 80% for missense variants seen in gnomAD (Fig. 1C). The high fraction of pathogenic variants in gnomAD is consistent with the majority of these variants having arisen recently. Indeed, recent exponential human population growth introduced large numbers of rare variants through random de novo mutations (95% of variants in the gnomAD cohort are at < 0.01% population allele frequency), without sufficient time for selection to purge deleterious variants from the population (*36–40*). Consequently, the gnomAD cohort provides a comparatively unfiltered look at variation caused by random mutations, whereas primate common variants represent the subset of random mutations that have survived.

The regions of human disease genes that were most densely populated by ClinVar pathogenic variants were also strongly depleted for primate common variants, with examples shown for *CACNA1A* (Fig. 1D) and *CREBBP* (fig. S4), genes responsible for familial epilepsy (*41, 42*) and Rubinstein-Taybi syndrome (*43, 44*). Missense variants in the gnomAD cohort were partially depleted within these same critical regions (Fig. 1D and fig. S4), indicating that humans and primates experience similar selective pressures. However, deleterious variants were incompletely removed in humans, consistent with the shorter amount of time they were exposed to natural selection.

Prior to using primate data as an indicator of benign consequence in a diagnostic setting, it is vital to understand why a handful of human pathogenic ClinVar variants appear as tolerated common variants in primates. Our clinical laboratory independently reviewed evidence for each of the 36 ClinVar pathogenic variants that appeared in the primate cohort, according to ACMG guidelines (*14*). Among these 36 variants, 8 were reclassified as variants of uncertain significance based on insufficient evidence of pathogenicity in the literature and an additional 9 were hypomorphic or mild clinical variants (table S1). The remaining 19 variants appear to be truly pathogenic in human, and are presumably tolerated in primate because of primate-human differences, such as interactions with changes in the neighboring sequence context (*45, 46*). In one such example, a compensatory synonymous sequence change at an adjacent nucleotide explains why the variant is benign in primate, but creates a pathogenic splice defect in human (Fig. 1E). We also expect that some of the variants identified among primates are rare pathogenic variants by chance, despite the small number of individuals sequenced within each species. By expanding our cohort to sequence a large number of individuals per species, we would definitively exclude rare variation from our catalogue of primate variation, as well as grow the database of benign variants to improve clinical variant interpretation.

As evolutionary distance from human increases, cases where the surrounding sequence context has changed sufficiently to alter the effect of the variant should also increase, until common variants in more distant species could no longer be reliably counted on as benign in human. We examined variation in each major branch of the primate tree, as well as variation from mammals (mouse, rat, cow, dog), chicken, and zebrafish, and evaluated their pathogenicity in ClinVar (Fig. 1F). Common variants from species throughout the primate lineage, including more distant branches such as lemurs and tarsiers, varied from 98.6% to 99% benign in the human ClinVar database, but this dropped to 87% for placental mammals, and 71% for chicken. The high fraction of variants that are pathogenic in human, yet tolerated as common variants in more distant vertebrates, indicates that selection on orthologous variants diverges substantially in distantly-related species, as a consequence of changes in the surrounding sequence context and other differences in the species’ biology (fig. S5).

We have made the primate population variant database, which contains over 4.3 million likely benign missense variants, publicly available at https://primad.basespace.illumina.com as a reference for the genomics community. Overall, this resource is over 50 times larger than ClinVar in terms of number of annotated missense variants, and consists almost entirely of variants of previously unknown significance. Most primate variants are rare or absent in the human population, with 98% of these variants at allele frequency < 0.01% (fig. S6). This makes it challenging to establish their pathogenicity through other means, since even the largest sequencing laboratories would be unlikely to observe any given variant in more than one unrelated patient. Despite their rarity, the subset of human variants that appear in primates have a low missense : synonymous ratio consistent with being depleted of deleterious missense variants (Fig. 1G). This contrasts with the high missense : synonymous ratio for rare human variants in the overall gnomAD cohort, which approaches the 2.2:1 ratio expected for random de novo mutations in the absence of selective constraint (*47*). At higher allele frequencies, natural selection has had more time to purge deleterious missense variants, allowing the human missense : synonymous ratio to start to converge toward the ratio observed for the subset of human variants that are present in other primates.

### Gene-level selective constraint in humans versus non-human primates

The primate variant resource makes it possible to compare natural selection acting on individual genes across the primate lineage and identify human-specific evolutionary differences. Since the current primate cohort only contains an average of 3-4 individuals per species, we focused on comparing selective constraint in human genes versus primates as a whole. We found that the missense : synonymous ratios of individual genes were well-correlated between human and primates (Spearman r = 0.637) (Fig. 2A), indicating that genes which were depleted for deleterious missense mutations in human were also consistently depleted throughout the primate lineage. Moreover, the missense : synonymous ratios of both human and primate genes correlated similarly well with the probability of genes being loss of function intolerant (pLI) (Spearman correlation -0.534 and -0.489, respectively) (*28*). Had there been substantial divergence between human and primate, pLI, an independent metric derived from human protein-truncating variation, would have been expected to show much clearer agreement with human missense : synonymous ratios than primate.

**Fig. 2.**
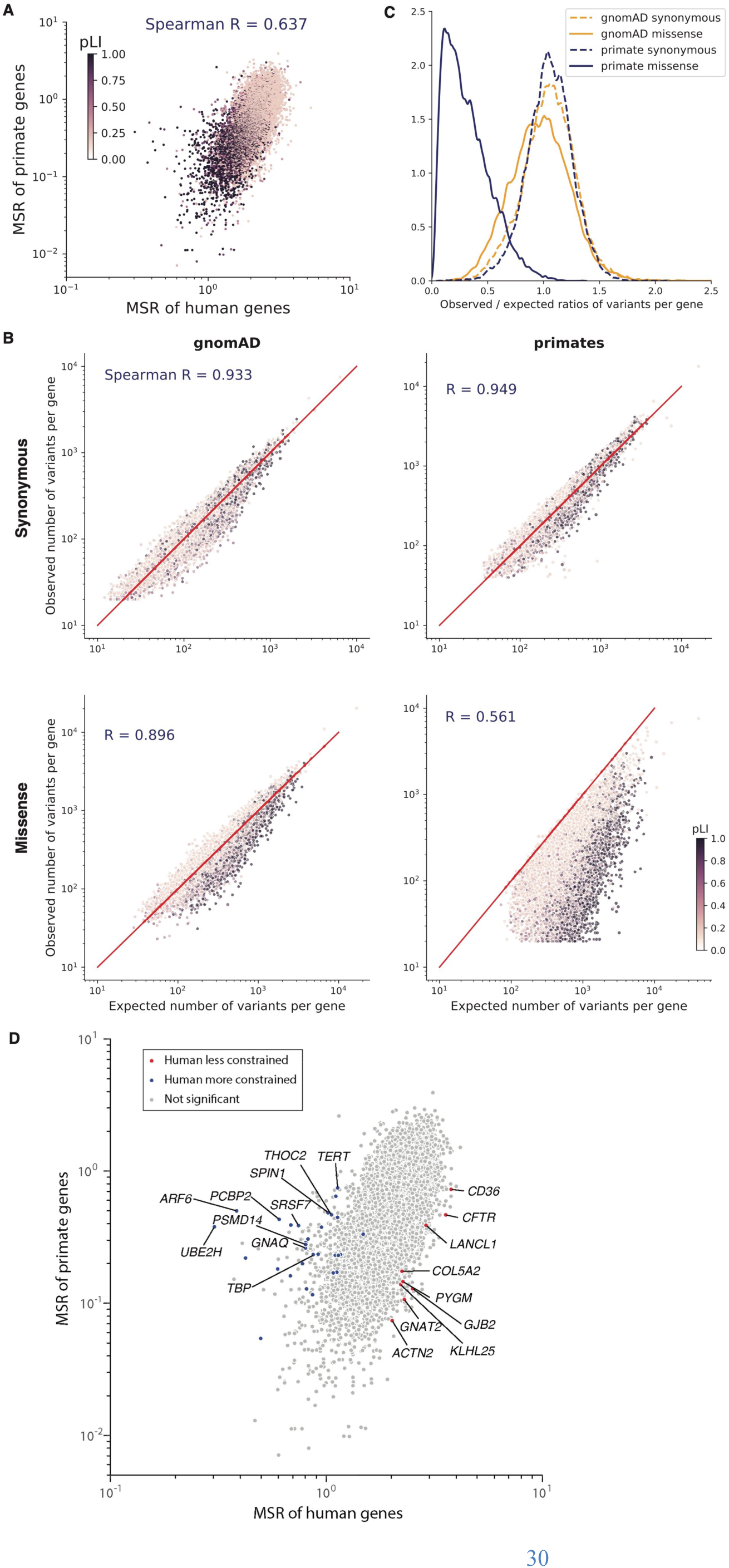
Selective constraint of primate genes compared to human. (**A**) Scatter plot of missense : synonymous ratios between primate and human genes. Each gene is colored by its pLI score, with darker points showing haploinsufficient genes. (**B**) Observed and expected counts of synonymous (top) and missense (bottom) variants per gene in gnomAD (left) and primates (right). Genes are colored by their pLI scores. (**C**) Distributions of observed/expected ratios of synonymous (dashed lines) and missense (solid lines) variants for all genes. Results for primate genes (orange) and gnomAD genes (blue) are shown. (**D**) Scatter plot of missense : synonymous ratios between primate and human genes. Highlighted points are genes that are under significantly stronger (blue) or weaker (red) constraint in humans compared to non-human primates under both methods (Benjamini-Hochberg FDR < 0.05), while grey points show non-significant genes. The top 10 genes with the largest effect sizes in either direction are labeled.

To measure the selective constraint on each gene, we calculated the observed versus expected number of variants per gene, using trinucleotide mutation rates to model the expected probability of observing each variant (fig. S7) (*28, 29*). We modeled each primate species separately to account for differences in genetic diversity and the number of individuals sampled per species.

The expected and observed counts of synonymous variants were highly correlated in both the gnomAD and primate cohorts, indicating that our model accurately captured the background distribution of neutral mutations (Fig. 2B; Spearman correlation 0.933 and 0.949, respectively). In contrast, for missense variants the expected and observed counts per gene diverged substantially (Spearman correlation 0.896 and 0.561 for human and primate, respectively), due to depletion of deleterious missense variants by natural selection in highly constrained genes (for example, high pLI genes). The most highly constrained genes were almost completely scrubbed of common missense variants in the primate cohort, whereas rare missense variants in the gnomAD cohort were depleted to a more modest extent due to the large sample size of gnomAD (Fig. 2C).

We next aimed to identify genes whose selective constraint was different in human compared to the rest of the primate lineage, a task made difficult by differences in diversity, allele frequency, and sample size between the human and primate cohorts (*34, 48, 49*). To this end, we developed two orthogonal strategies, and took the intersection of genes identified under both approaches.

First, we used population genetic modeling (*34, 50, 51*) to estimate the average selection coefficient, *s*, ranging from 0 (benign) to 1 (severely pathogenic), of missense mutations in each gene, using a model of recent human population growth (figs. S7 and S8). We fit a single value of *s* per gene across non-human primate species, and identified genes that differed between *s_primate_* and *s_human_* using a likelihood ratio test, which we validated using population simulations (fig. S9). In a second approach, we fit a curve approximating the relationship between human and primate missense : synonymous ratios using a Poisson generalized linear mixed model (*52*), and identified genes where the observed human missense : synonymous ratio deviated from what would have been expected given the gene’s missense : synonymous ratio in primates (fig. S10). We also adjusted for gene length to account for shorter genes having more variability in their missense : synonymous ratio measurements than longer genes. The two methods were broadly concordant, with a Spearman correlation of 0.80 between the genes’ effect sizes in the two tests. Estimates of selection coefficients and observed and expected counts for each gene in human and primate are provided in table S2.

In total, we found 39 genes where selective constraint differed significantly between human and other primates under both methods (Benjamini-Hochberg FDR < 0.05 (*53*); Fig. 2D). The top three genes where *s_human_* decreased the most relative to *s_primate_* were *CFTR*, *GJB2*, and *CD36*, autosomal recessive disease genes for cystic fibrosis (*54*), hereditary deafness (*55*), and platelet glycoprotein deficiency (*56*), respectively. All three genes are known for deleterious mutations that are unusually common in local geographic human populations (*57–60*), suggesting that they may be experiencing reduced selection due to heterozygote advantage that protects against specific environmental pathogens (*60–64*). On the other end of the spectrum, *TERT*, known for its role in maintaining telomere length (*65, 66*), was among the top genes where *s_human_* increased the most relative to *s_primate_*. Humans have adapted to a much longer lifespan compared with other primate species, which have a median lifespan of 20-30 years, suggesting that increased selection on *TERT* may have occurred as part of human adaption towards extended longevity. We note that with the current size of the primate cohort, it is not possible to distinguish whether the increased selection on *TERT* occurred only in humans, or if it is part of a gradual trend towards extended longevity that began earlier in the great ape lineage, which also have longer lifespans relative to other primates (∼40 years). Expanding the primate cohort by sequencing more individuals per species would improve detection of additional species-specific and lineage-specific evolutionary adaptation, and shed light on the evolutionary path that led to the present human condition.

### PrimateAI-3D, a deep learning network for classifying protein-altering variants

We constructed PrimateAI-3D, a semi-supervised 3D-convolutional neural network for variant pathogenicity prediction, which we trained using 4.5 million common missense variants with likely benign consequence (Fig. 3A). In a departure from prior deep learning architectures that operated on linear sequence (*17, 67*), we voxelized the 3D structure of the protein at 2 Angstrom resolution (figs. S11 and S12) and used 3D-convolutions to enable the network to recognize key structural regions that may not be apparent from sequence alone (Fig. 3A). As an example, we show PrimateAI-3D predictions for *STK11* (Fig. 3B), the tumor suppressor gene responsible for Peutz-Jeghers hereditary polyposis syndrome (*68–71*), with each amino acid position colored by the average PrimateAI-3D score at that position. Common primate variants used for training and annotated ClinVar pathogenic variants from separate parts of the linear sequence form distinct clusters in 3D space. Although ClinVar variants are shown for illustration, it is important to note that the network was not trained on either human-engineered features or annotated variants from clinical variant databases, thereby avoiding potential human biases in variant annotation. Rather, it learns to infer pathogenicity based on the local enrichment or depletion of common primate variants, taking only the protein’s multiple sequence alignment and 3D structure as inputs.

**Fig. 3.**
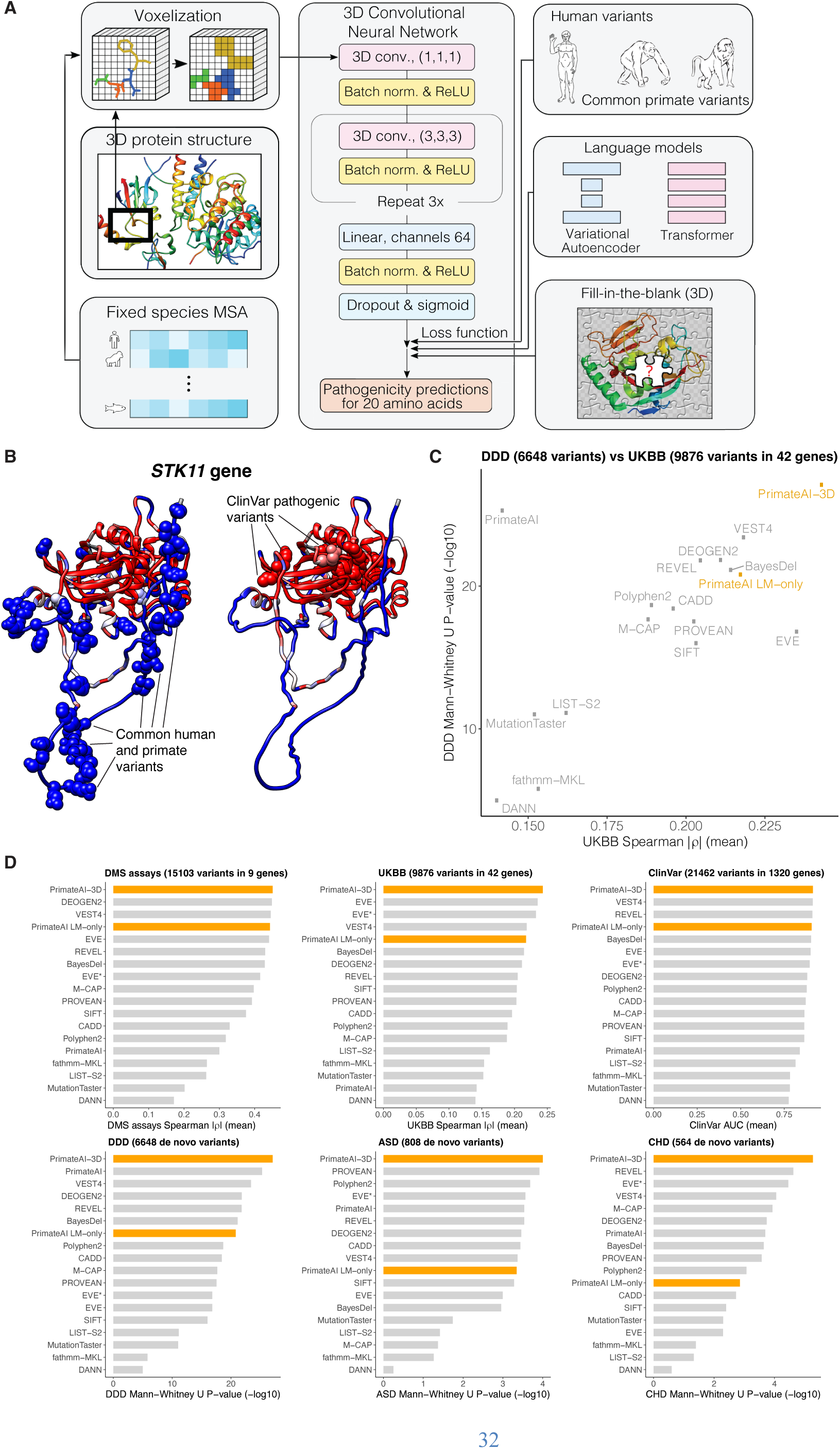
PrimateAI-3D architecture and variant classification performance. (**A**) PrimateAI-3D workflow. Human protein structures and multiple sequence alignments are voxelized (left) as input to a 3D convolutional neural network that predicts pathogenicity of all possible point mutations of a target residue (middle). The network is trained using a loss function with three components (right): common human and primate variants; fill-in-the-blank of a protein structure; score ranks from language models. (**B**) Protein structure of the *STK11* gene, colored by PrimateAI-3D pathogenicity prediction scores (blue: benign; red: pathogenic). Spheres indicate residues with common human and primate variants (left) or residues with pathogenic mutations from ClinVar (right). For spheres, the color corresponds to the pathogenicity score of only the variant. For other residues, pathogenicity scores are averaged over all variants at that site. (**C**) Scatterplot shows performance of methods that predict missense variant pathogenicity in two clinical benchmarks (DDD and UKBB). Datasets are a subset of variants for which all methods have predictions. (D) Six barplots show method performance for six testing datasets (DMS assays, UKBB, ClinVar, DDD, ASD, and CHD).

PrimateAI-3D can utilize protein structures from either experimental sources or computational prediction (*72–76*); we used AlphaFold DB (*72, 73*) and HHpred (*74*) predicted structures for the broadest coverage across human genes. For training data, we incorporated all common missense variants from the 233 non-human primate species (*17*), and common human missense variants (allele frequency > 0.1% across populations) in gnomAD (*28, 29*), TOPMed (*77, 78*), and UK Biobank (*79, 80*), resulting in a total of 4.5 million unique missense variants of likely benign consequence. This dataset covers 6.34% of all possible human missense variants, and is over 50-fold larger than the current ClinVar database (79,381 missense variants after excluding variants of uncertain significance and those with conflicting annotations), greatly enlarging the training dataset available for machine learning approaches. Because the training dataset consists only of variants labeled as benign, we created a control set of randomly selected variants that were matched to the common variants by trinucleotide mutation rate, and trained PrimateAI-3D to separate common variants from matched controls as a semi-supervised learning task.

In parallel with the variant classification task, we generated amino acid substitution probabilities for each position in the protein by masking the residue and using the sequence context to predict the missing amino acid, borrowing from language model architectures that are trained to predict missing words in sentences (*81, 82*). We trained both a 3D convolutional “fill-in-the-blank” model, which tasked the network with predicting the missing amino acid in a gap in the voxelized 3D protein structure, and separately, a language model utilizing the transformer architecture to predict the missing amino acid using the surrounding multiple sequence alignment as context (*83*). We implemented these models as additional loss functions to further refine the PrimateAI-3D predictions (fig. S13). We also trained a variational autoencoder (*67*) on multiple sequence alignments and found that it performed comparably to our transformer architecture (fig. S14). Hence, we incorporated the average of their predictions in the loss function, which performed better than either alone.

We evaluated PrimateAI-3D and 15 other published machine learning methods (*67, 84*) on their ability to distinguish between benign and pathogenic variants along six different axes (Fig. 3C, 3D, and fig. S15): predicting the effects of rare missense variants on quantitative clinical phenotypes in a cohort of 200,643 individuals from the UK Biobank (UKBB); distinguishing missense de novo mutations (DNM) seen in 31,058 patients with neurodevelopmental disorders (*85–87*) (DDD) from de novo missense mutations in 2,555 healthy controls (*88–93*); distinguishing de novo missense mutations seen in 4,295 patients with autism spectrum disorders (*88–94*) (ASD) from de novo missense mutations in the shared set of 2,555 healthy controls; distinguishing de novo missense mutations seen in 2,871 patients with congenital heart disease (*95*) (CHD) from de novo missense mutations in the shared set of 2,555 healthy controls; separating annotated ClinVar benign and pathogenic variants (ClinVar) (*4*); and average correlation with in vitro deep mutational scan experimental assays across 9 genes (*96–105*) (DMS assays). Our set of clinical benchmarks is the most comprehensive to date, and has a particular focus on rigorously testing the performance of classifiers on large patient cohorts across a diverse range of real world clinical settings (table S3).

For the UK Biobank benchmark, we analyzed 200,643 individuals with both exome sequencing data and broad clinical phenotyping, and identified 42 genes where the presence of rare missense variants was associated with changes in a quantitative clinical phenotype controlling for confounders such as population stratification, age, sex, and medications (table S4). These gene-phenotype associations included diverse clinical lab measurements such as low-density lipoprotein (LDL) cholesterol (increased by rare missense variants in *LDLR*, decreased by variants in *PCSK9*), blood glucose (increased by variants in *GCK*), and platelet count (increased by variants in *JAK2*, decreased by variants in *GP1BB*), as well as other quantitative phenotypes such as standing height (increased by variants in *ZFAT*) (table S4). To test each classifier’s ability to distinguish between pathogenic and benign missense variants, we measured the correlation between pathogenicity prediction score and quantitative phenotype for patients carrying rare missense variants in each of these genes. We report the average correlation across all gene-phenotype pairs for each classifier, taking the absolute value of the correlation, since these genes may be associated with either increase or decrease in the quantitative clinical phenotype.

The neurodevelopmental disorders cohort (DDD), autism spectrum disorders cohort (ASD), and congenital heart disease cohort (CHD) are among the largest published trio-sequencing studies to date, and consist of thousands of families with a child with rare genetic disease and their unaffected parents. In each cohort, we cataloged de novo missense mutations that appeared in affected probands but were absent in their parents, as well as de novo missense mutations that appeared in a set of shared healthy controls. We evaluated the ability of each classifier to separate the de novo missense mutations that appear in cases versus controls on the basis of their prediction scores, using the Mann-Whitney U test to measure performance.

PrimateAI-3D outperformed all other classifiers at distinguishing pathogenic from benign variants in the four patient cohorts we tested (UKBB, DDD, ASD, CHD); it was also the top performer at separating pathogenic from benign variants in the ClinVar annotation database, and had the highest average correlation with the deep mutational scan assays (Fig. 3D and fig. S15). After PrimateAI-3D, there was no clear runner-up, with second place occupied by six different classifiers in the six different benchmarks. We observed a moderate correlation between the performance of different classifiers in UKBB and DDD (Spearman r = 0.556; Fig. 3C), which are the two largest clinical cohorts and therefore likely the most robust for benchmarking (with 200,643 and 33,613 patients, respectively), but outside of PrimateAI-3D, strong performance of a classifier on one task had limited generalizability to other tasks. Our results underscore the importance of validating machine learning classifiers along multiple dimensions, particularly in large real-world cohorts, to avoid overgeneralizing a classifier’s performance based upon an impressive showing along a single axis.

PrimateAI-3D’s top-ranked performance at separating benign and pathogenic missense variants in ClinVar was unexpected, since the other machine learning classifiers (with the exception of EVE) were trained either directly on ClinVar, or on other variant annotation databases with a high degree of content overlap. Because they are primarily based on variants described in the literature, clinical variant databases are subject to ascertainment bias (*12, 106, 107*), which may have contributed to supervised classifiers picking up on tendencies of human variant annotation that are unrelated to the task of separating benign from pathogenic variants (figs. S16, S17, and S18). Given the challenges with human annotation, we also investigated whether PrimateAI-3D could assist in revising incorrectly labeled ClinVar variants, by comparing annotations in the current ClinVar database and those from a September 2017 snapshot. Disagreement between PrimateAI-3D and the 2017 version of ClinVar was highly predictive of future revision and the odds of revision increased with PrimateAI-3D confidence (fig. S19). Among variants with the 10% most confident PrimateAI-3D predictions, the odds of revision were 10-fold elevated if PrimateAI-3D was in disagreement with the ClinVar label (*P* < 10^-14^).

The performance of PrimateAI-3D on clinical variant benchmarks scaled directly with training dataset size, indicating that additional primate sequencing data will be the key to unlocking further gains (Fig. 4 and fig. S20). The current primate cohort already covers 30% of all possible synonymous variants in the human genome, despite containing only 809 individuals from 233 species (Fig. 4B). By increasing the number of species and the number of individuals sequenced per species, we expect to saturate the majority of the remaining tolerated substitutions in the human genome (fig. S21), including both coding and noncoding variation, leaving the remaining deleterious variants to be deduced by process of elimination.

**Fig. 4.**
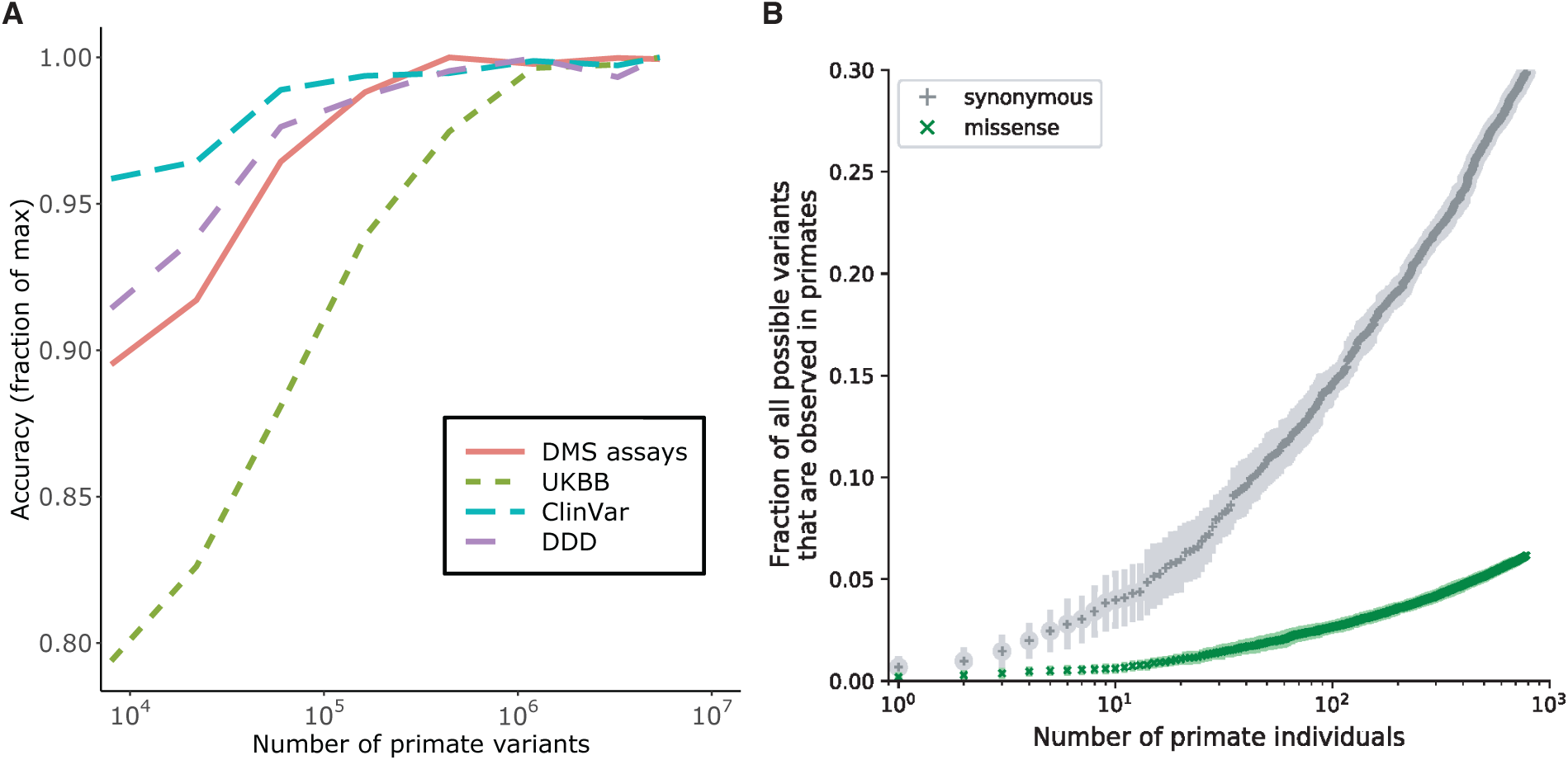
Impact of training dataset size on classification accuracy. (**A**) Improved performance of PrimateAI-3D with increasing number of common human and primate variants in the training dataset (x-axis). Performance of each dataset (y-axis) was divided by the maximum performance observed across all training dataset sizes. (**B**) Cumulative fractions of all possible human synonymous (grey) and missense (green) variants observed as common variants in 234 primate species, including human (allele frequency > 0.1%). Each point shows the average of ten permutations, calculated with a different random ordering of the list of primate species each time.

### Discovery of candidate disease genes for neurodevelopmental disorders

We applied PrimateAI-3D to improve statistical power for discovering candidate disease genes that are enriched for pathogenic de novo mutations in the neurodevelopmental disorders cohort (fig. S22). De novo missense mutations from affected individuals in the DDD cohort (*87*) were enriched 1.36-fold above expectation, based on estimates of background mutation rate using trinucleotide context (*47*). We selected a PrimateAI-3D classification threshold of 0.821, which called an equal number of pathogenic missense mutations (n=7,238) as the excess of de novo missense mutations in the cohort (Fig. 5A). Stratifying missense mutations by this threshold increased enrichment of pathogenic de novo missense mutations to 2.0-fold, substantially increasing statistical power for disease gene discovery in the cohort (Fig. 5B).

**Fig. 5.**
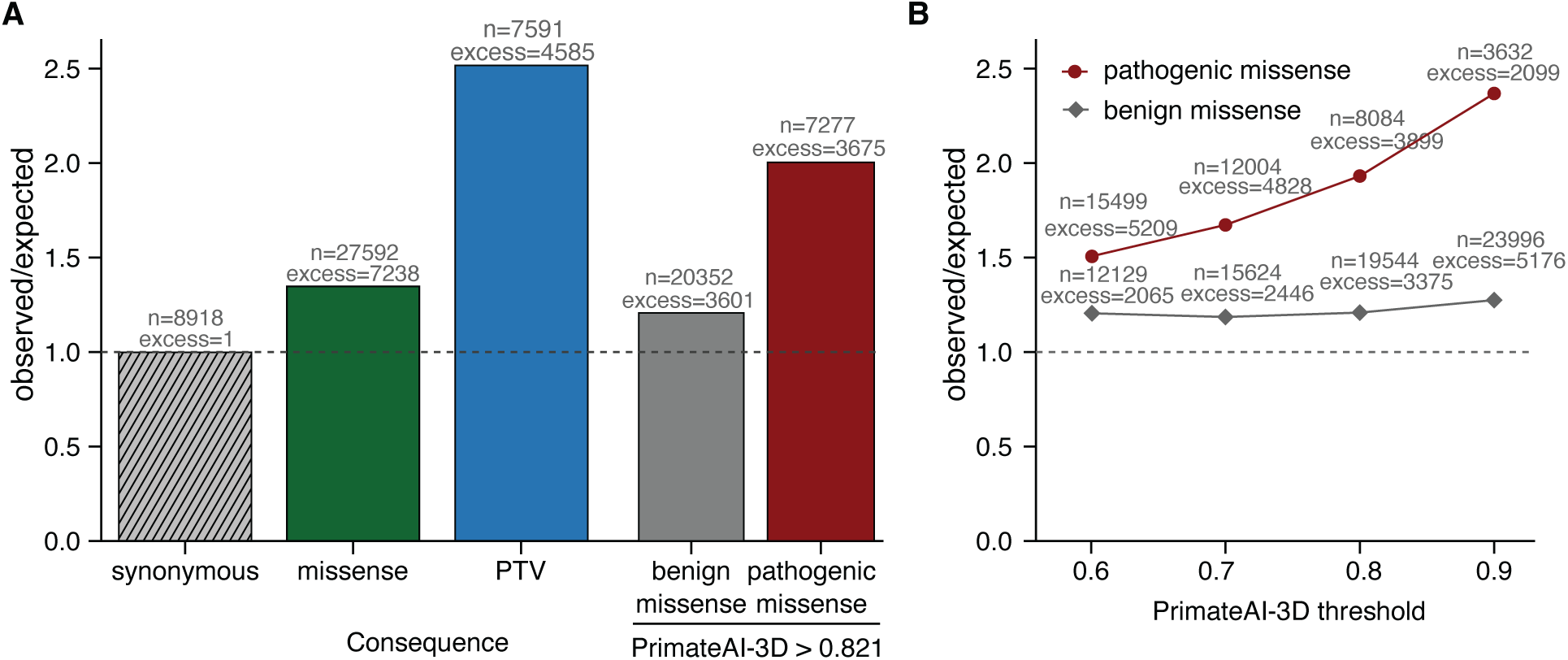
Enrichment of de novo mutations in the neurodevelopmental disorder cohort over expectation. (**A**) Enrichment of DNMs from Kaplanis *et al.* (*87*) across all genes. Enrichment ratios are given for synonymous, all missense, and protein-truncating variants (PTV), along with missense split by PrimateAI-3D score into benign (<0.821) and pathogenic (>0.821). (**B**) Enrichment of benign and pathogenic missense above expectation at varying PrimateAI-3D thresholds for classifying pathogenic missense.

By applying PrimateAI-3D to prioritize pathogenic missense variants, we identified 290 genes associated with intellectual disability at genome-wide significance (*P* < 6.4×10^-7^) (Table 1), of which 272 were previously discovered genes that either appeared in the Genomics England intellectual disability gene panel (*108*), or were already identified in the prior study (*109*) without stratifying missense variants (table S5). We excluded two genes, *BMPR2* and *RYR1* as borderline significant genes that already had well-annotated non-neurological phenotypes. Further clinical studies are needed to independently validate this list of candidate genes and understand their range of phenotypic effects.

**Table 1.** Additional genes discovered in intellectual disability. Genes achieving the genome-wide significance (p < 6.4×10^-7^) are shown when considering only missense de novo mutations with PrimateAI-3D scores ≥0.821. Counts of protein truncating and missense DNMs are provided. *P* values for gene enrichment are shown when the statistical test was run only with missense mutations with PrimateAI-3D score ≥ 0.821, and when it was repeated for all missense mutations.

| HGNC symbol | Protein-truncating variants | Missense |  | *P* value |  |
| --- | --- | --- | --- | --- | --- |
| | | PrimateAI-3D score $\geq 0.821$ | All missense | PrimateAI-3D score $\geq 0.821$ | All missense |
| *AP1G1* | 2 | 4 | 5 | $4.1 \times 10^{-7}$ | $5.9 \times 10^{-5}$ |
| *ATP2B2* | 1 | 9 | 11 | $2.1 \times 10^{-7}$ | $1.4 \times 10^{-3}$ |
| *CELF2* | 2 | 4 | 4 | $1.2 \times 10^{-7}$ | $6.7 \times 10^{-5}$ |
| *MAP4K4* | 2 | 6 | 7 | $3.9 \times 10^{-7}$ | $5.0 \times 10^{-4}$ |
| *MED13* | 3 | 6 | 9 | $6.6 \times 10^{-8}$ | $3.5 \times 10^{-5}$ |
| *MFN2* | 0 | 6 | 8 | $3.4 \times 10^{-7}$ | $1.0 \times 10^{-5}$ |
| *NR4A2* | 2 | 4 | 5 | $3.7 \times 10^{-7}$ | $3.3 \times 10^{-5}$ |
| *PIP5K1C* | 0 | 8 | 9 | $2.8 \times 10^{-8}$ | $4.9 \times 10^{-4}$ |
| *RAB5C* | 2 | 4 | 5 | $8.6 \times 10^{-8}$ | $1.5 \times 10^{-5}$ |
| *SPOP* | 1 | 4 | 6 | $4.1 \times 10^{-7}$ | $1.7 \times 10^{-6}$ |
| *SPTBN2* | 1 | 10 | 16 | $3.9 \times 10^{-7}$ | $4.5 \times 10^{-3}$ |
| *XPO1* | 1 | 7 | 7 | $5.0 \times 10^{-7}$ | $7.2 \times 10^{-4}$ |
| *EIF4A2* | 2 | 4 | 4 | $1.7 \times 10^{-7}$ | $2.1 \times 10^{-4}$ |
| *LMBRD2* | 0 | 3 | 4 | $6.0 \times 10^{-7}$ | $1.3 \times 10^{-4}$ |
| *MARK2* | 4 | 3 | 5 | $2.3 \times 10^{-7}$ | $3.8 \times 10^{-5}$ |
| *NOTCH1* | 4 | 6 | 17 | $4.1 \times 10^{-7}$ | $1.3 \times 10^{-6}$ |

## Discussion

Our results demonstrate the successful pairing of primate population sequencing with state-of-the-art deep learning models to make meaningful progress towards solving variants of uncertain significance. Primate population sequencing and large-scale human sequencing are likely to fill complementary roles in advancing clinical understanding of human genetic variants. From the perspective of acquiring additional benign variants to train PrimateAI-3D, humans are not suitable, as the discovery of common human variants (>0.1% allele frequency) plateaus at roughly ∼100,000 missense variants after only a few hundred individuals (*17*), and further population sequencing into the millions mainly contributes rare variants which cannot be ruled out for deleterious consequence. On the other hand, these rare human variants, because they have not been thoroughly filtered by natural selection, preserve the potential to exert highly penetrant phenotypic effects, making them indispensable for discovering new gene-phenotype relationships in large population sequencing and biobank studies. Fittingly, classifiers trained on common primate variants may accelerate these target discovery efforts, by helping to differentiate between benign and pathogenic rare variation.

The genetic diversity found in the 520 known non-human primate species is the result of ongoing natural experiments on genetic variation that have been running uninterrupted for millions of years. Today, over 60% of primate species on Earth are threatened with extinction in the next decade due to man-made factors (*31*). We must decide whether to act now to preserve these irreplaceable species, which act as a mirror for understanding our genomes and ourselves, and are each valuable in their own right, or bear witness to the conclusion of many of these experiments.

## Materials and Methods

### Primate polymorphism data

We aggregated high-coverage whole genomes of 809 primate individuals across 233 primate species, including 703 newly sequenced samples and 106 previously sequenced samples from the Great Ape Genome project (*19*). Samples that passed quality evaluation were then aligned to 32 high-quality primate reference genomes (*110*) and mapped to the GRCh38 human genome build.

We developed a random forest (RF) classifier to identify false positive variant calls and errors resulting from ambiguity in the species mapping. In addition, we removed variants that fell in primate codons that did not match the human codon at that position, as well as those residing in primate transcripts with likely annotation errors. We also devised quality metrics based on the distribution of RF scores and Hardy-Weinberg equilibrium, and developed a unique mapping filter to exclude variants in regions of non-unique mapping between primate species.

### Identifying differential selection between humans and primates via population modeling

We first established a neutral background distribution of mutation rates per gene for each primate species by fitting the Poisson Random Field (PRF) model to the segregating synonymous variants in each species. The observed number of segregating synonymous sites is a Poisson random variable, with the mean determined by mutation rate, demography, and sample size (*34*). For simplicity, we assumed an equilibrium (i.e. constant) demography for all species besides human; for human, we used Moments (*51*) to find a best fitting demographic history based on the folded site frequency spectrum of synonymous sites. We adopted a Gamma distributed prior on mutation rates, which also accounts for the impact of GC content on mutation rate. We optimized the prior parameters via maximum likelihood and computed the posterior distribution of the mutation rate per gene.

The number of segregating nonsynonymous sites is modeled as a Poisson random variable similar to synonymous sites with additional selection parameters. We assumed that every nonsynonymous mutation in a gene shares the same population scaled selection coefficient γ_!“_. To explicitly estimate selection coefficient of each gene per species, we devised a two-step procedure analogous to an EM algorithm to control for differences in population size across species.

To identify genes where human constraint is different from non-human primate selection, we developed a likelihood ratio test to test whether population scaled selection coefficients are significantly different between human and other primates. We then assessed whether our population genetic modeling improved the correlation of selection estimates of our primate data with previous gene-constraint metrics in humans, including pLI (*28*) and s_het (*111*). To validate the performance of our model, we performed population genetic simulations.

### Poisson generalized linear mixed modeling of selection between humans and primates

In addition to population genetics model described above, we also applied an orthogonal approach to detect differences in selection between humans and primates based on missense-to-synonymous ratio (MSR). We fit a Poisson generalized linear mixed model (GLMM) to the pooled polymorphic synonymous and missense mutations across all primates to estimate the depletion of missense variants in each gene. Then, we fit a second Poisson GLMM to the human data, controlling for the primate depletion estimates, and compared the pooled primate MSR to the human MSR for each gene.

### PrimateAI-3D Model

PrimateAI-3D is a 3D convolutional neural network that uses protein structures and multiple sequence alignments (MSA) to predict the pathogenicity of human missense variants. To generate the input for a 3D convolutional neural network, we voxelized the protein structure and evolutionary conservation in the region surrounding the missense variant. The network was trained to optimize three objectives: distinction between benign and unknown human variants; prediction of a masked amino acid at the variant site; per-gene variant ranks based on protein language models.

### Protein structures and multiple sequence alignments

For 341 species, we used vertebrate and mammal MSAs from UCSC Multiz100 (*112, 113*) and Zoonomia (*114*). Another 251 species appeared in Uniprot for least 75% of all human proteins (*115*). For each protein, alignments from all 341+251=592 species were merged. Human protein structures were taken from AlphaFold DB (June 2021) (*73*). Proteins that did not sequence-match exactly to our hg38 proteins (2590; 13.5%) were homology modeled using HHpred (*74*) and Modeller (*116*).

### Protein voxelization and voxel features

A regular sized 3D grid of 7×7×7 voxels, each spanning 2Åx2Åx2Å, was centered at the Cα atom of the residue containing the target variant (Fig. S11). For each voxel, we provided a vector of distances between its center and the nearest Cα and Cβ atoms of each amino acid type (Fig. S11; details in Supplementary Text section 1). We also provided additional voxel features including the pLDDT confidence metric from AlphaFold DB (Fig. S12), and the evolutionary profile, consisting of each amino acid’s frequency at the corresponding position in the 592 species alignment.

### Model architecture

The first layers of the PrimateAI-3D model reduce the voxel tensor to a 64-vector through repeated valid-padded 3D convolutions with a kernel size of 3×3×3. A final hidden dense layer transforms this 64-length vector into a 20-length vector, corresponding to one output unit per amino acid at that position. The model was trained simultaneously using multiple loss functions to optimize the following complementary aspects of pathogenicity:

#### Benign primate variants

Using 4.5 million benign missense variants from primates, we sampled the same number of unknown variants from the set of all possible human missense variants, with the distribution of mutational probabilities matching the benign set, based on a trinucleotide mutation rate model. Variants for the same protein position were combined in a 20-length vector (benign: 0, unknown: 1) which was the target label for the network. We used mean squared error (MSE) as the loss function for non-missing labels and ignored missing labels.

#### 3D fill-in-the-blank

We removed all atoms of a target residue before voxelization, discarding any information about the residue from the input tensor to the network. The network was then trained to predict a 20-length vector, labeled 0 (benign) for amino acids that occur at the target site in any of the 592 species and 1 (pathogenic) otherwise. All human protein positions with at least one possible missense variant were included in this dataset.

#### Variant ranks from language models

For each gene, we took the average pathogenicity ranking from two protein language models, PrimateAI language model (PrimateAI LM, described below) and our reimplementation of the EVE variational autoencoder algorithm which we extended to all human proteins (EVE*) (*67*). We calculated the pairwise logistic rank loss as described in Pasumarthi *et al.*(*117*).

### PrimateAI Language Model

The PrimateAI language model (PrimateAI LM) is a MSA transformer (*83*) for fill-in-the-blank residue classification, which was trained end-to-end on MSAs of UniRef-50 proteins (*118, 119*) to minimize an unsupervised masked language modelling (MLM) objective (*81*). Our model requires ∼50x less computation for training than previous MSA transformers due to several improvements in architecture and training (Fig. S9).

### Model training procedure

Each batch had the same number of samples from each of the three variant datasets (∼33 with a batch size of 100). For the language model ranks dataset, all 33 samples had to come from the same protein. The number of times a protein was chosen for a batch was proportional to the length of the protein. In order to make our model robust against protein orientations, we randomly rotated the protein atomic coordinates in 3D before voxelizing a variant.

### Model Evaluation

We compared performance of our model and other models (*84*) on variants for which all models had scores. Deep mutational scanning assays were available for 9 human genes: Amyloid-beta (*102*), *YAP1* (*96*), *MSH2* (*120*), *SYUA* (*101*), *VKOR1* (*121*), *PTEN* (*99, 100*), *BRCA1* (*122*), *TP53* (*123*), and *ADRB2* (*124*). For each assay and prediction model, we calculated the absolute Spearman rank correlation between prediction and assay scores. The UK Biobank dataset (*79, 80*) contains 42 gene-phenotype pairs which were significantly associated by rare variant burden testing using all rare missense variants, without applying missense pathogenicity prioritization. The evaluation was the same as with DMS assays, except that correlations were calculated from the quantitative phenotypes of individuals carrying the variant, instead of the assay score for the variant. For ClinVar (*4*), we filtered to high-quality 2-star variants and evaluated model performance by calculating per-gene area under the receiver operating characteristic curve (AUC). For the rare disease cohorts, we collected de novo missense mutations from patients with developmental disorders (*85–87*), autism spectrum disorders (*88–94*) or congenital heart disorders (*95*). For all three datasets, we compared against DNMs from healthy controls (*88–93*). We applied the Mann-Whitney U test to measure how well each model’s prediction scores could distinguish patient variants from control variants.

## Acknowledgments

We would like to thank Daniel MacArthur, Yun Song, and Mark Daly for helpful discussions, and the gnomAD team at the Broad Institute for their assistance with the website.

## Author contributions

HG, TH, JE, JGS, JM, MSB, YY, AD, PF, LK, LS, YW, AA, YF, SC, SB, GL, RR, DB, FA, KF performed the analysis and wrote the manuscript. MCJ, MK, JDO, SM, AV, JB, MR, FES, LA, JB, MG, DdV, IG, RAH, MR, AJ, ISC, JH, CH, DJ, PF, FRdM, FB, HB, IS, IF, JVdA, MM, MNFdS, MT, RR, TH, NA, CJR, AZ, CJJ, JPC, GW, CA, JHS, EFD, SK, FS, DW, LZ, YS, GZ, JDK, SK, MDL, EL, SM, AN, TB, TN, CCK, JL, PT, WKL, ACK, DZ, IG, AM, KG, MHS, RMDB, GU, CR, JPB contributed the primate samples and sequencing data. ML, SS, AOD, HR, JX, JR, TMB, and KF supervised the work.

## Data and materials availability

All sequencing data have been deposited at the European Nucleotide Archive under the accession number PRJEB49549. Primate variants and PrimateAI-3D prediction scores are available with a non-commercial license upon request and are displayed on https://primad.basespace.illumina.com. The source code of PrimateAI-3D is accessible via https://github.com/Illumina/PrimateAI-3D and is also archived at https://doi.org/10.5281/zenodo.7738731. To reduce problems with circularity that have become a concern for the field, the authors explicitly request that the prediction scores from the method not be incorporated as a component of other classifiers, and instead ask that interested parties employ the provided source code and data to directly train and improve upon their own deep learning models.

## Supplementary Materials

Materials and Methods Supplementary Text Figs. S1 to S28 Tables S1 to S6 References (*125–169*)

