## Supplementary materials for "The landscape of tolerated genetic variation in humans and primates"

Science
AAAS

Supplementary Materials for

### The landscape of tolerated genetic variation in humans and primates

Hong Gao[1]†, Tobias Hamp[1]†, Jeffrey Ede[1], Joshua G. Schraiber[1], Jeremy McRae[1], Moriel Singer-Berk[2], Yanshen Yang[1], Anastasia Dietrich[1], Petko Fiziev[1], Lukas Kuderna[1,3], Laksshman Sundaram[1], Yibing Wu[1], Aashish Adhikari[1], Yair Field[1], Chen Chen[1], Serafim Batzoglou[1]‡, Francois Aguet[1], Gabrielle Lemire[2,4], Rebecca Reimers[4], Daniel Balick[5], Mareike C. Janiak[6], Martin Kuhlwilm[3,7,8], Joseph D. Orkin[3,9], Shivakumara Manu[10,11], Alejandro Valenzuela[3], Juraj Bergman[12,13], Marjolaine Rouselle[12], Felipe Ennes Silva[14,15], Lidia Agueda[16], Julie Blanc[16], Marta Gut[16], Dorien de Vries[6], Ian Goodhead[6], R. Alan Harris[17], Muthuswamy Raveendran[17], Axel Jensen[18], Idriss S. Chuma[19], Julie Horvath[20,21,22,23,24], Christina Hvilsom[25], David Juan[3], Peter Frandsen[25], Fabiano R. de Melo[26], Fabricio Bertuol[27], Hazel Byrne[28], Iracilda Sampaio[29], Izeni Farias[27], João Valsecchi do Amaral[30,31,32], Mariluce Messias[33,34], Maria N. F. da Silva[35], Mihir Trivedi[11], Rogerio Rossi[36], Tomas Hrbek[27,37], Nicole Andriaholinirina[38], Clément J. Rabarivola[38], Alphonse Zaramody[38], Clifford J. Jolly[39], Jane Phillips-Conroy[40], Gregory Wilkerson[41]§, Christian Abee[42], Joe H. Simmons[41], Eduardo Fernandez-Duque[42,43], Sree Kanthaswamy[44], Fekadu Shiferaw[45], Dongdong Wu[46], Long Zhou[47], Yong Shao[46], Guojie Zhang[47,48,49,50,51], Julius D. Keyyu[52], Sascha Knauf[53], Minh D. Le[54], Esther Lizano[3,55], Stefan Merker[56], Arcadi Navarro[3,57,58,59], Thomas Batallion[12], Tilo Nadler[60], Chiea Chuen Khor[61], Jessica Lee[62], Patrick Tan[61,63,64], Weng Khong Lim[63,64,65], Andrew C. Kitchener[66,67], Dietmar Zinner[68,69,70], Ivo Gut[16,71], Amanda Melin[72,73], Katerina Guschanski[18,74], Mikkel Heide Schierup[12], Robin M. D. Beck[6], Govindhaswamy Umapathy[10,11], Christian Roos[75], Jean P. Boubli[6], Monkol Lek[76], Shamil Sunyaev[77,5], Anne O'Donnell[2,4,78], Heidi Rehm[2,79], Jinbo Xu[1,80], Jeffrey Rogers[17]*¶, Tomas Marques-Bonet[3,16,55,57]*, Kyle Kai-How Farh[1]*

**This PDF file includes:**

Materials and Methods
Supplementary Text
Figs. S1 to S28
Captions for Tables S1 to S6

References *(125-169)*

### Materials and Methods

#### Data generation

##### *Generation of canonical gene set*

All coordinates used in the paper refer to human genome build UCSC hg38 / GRCh38, including the coordinates for variants in other species. Protein-coding DNA sequences and multiple sequence alignments of 99 vertebrate genomes with human were downloaded from the UCSC genome browser for the hg38 build. (http://hgdownload.soe.ucsc.edu/goldenPath/hg38/multiz100way/alignments/knownCanonical.exonNuc.fa.gz) (*112, 113*). For genes with multiple canonical gene annotations, the coding transcript with the highest conservation score across the MSA was selected. In total, 19,158 transcripts were selected to represent the canonical gene sets used for all the analyses in this paper.

##### *Human polymorphism data*

We downloaded human polymorphism data from genome Aggregation Database v2.1.1, which collected the whole-exome sequencing data of 125,748 individuals (gnomad.exomes.r2.1.1.sites.vcf.bgz file) and the whole-genome sequencing of 71,702 individuals from gnomAD v3.0 (http://gnomad.broadinstitute.org/) (*28, 29, 125*). For the subsequent analyses using gnomAD dataset, we used a merged variant set between gnomAD WES and WGS. We also collected variants from 65K TOPMed WGS (*77, 78*) and 200K UK Biobank WGS (*79, 80*) and merged those with gnomAD (after removing 2,404 TOPMed samples from gnomAD). We excluded variants that failed the default quality control filters as annotated in VCF files or fell outside canonical coding regions. To avoid effects due to balancing selection, we also excluded variants from the extended MHC region (chr6: 28,510,120 – 33,480,577) for the subsequent analyses. In total, we obtained 126,873 unique common missense variants (allele frequency > 0.1%) and 7,306,297 unique rare missenses (allele frequency < 0.1%), which were included in the benign set for training of primateAI-3D. We also performed mutation rate correction on gnomAD variants following our previous paper (*17*) for missense : synonymous ratio analyses.

##### *Primate sequencing data*

We sequenced and aggregated the whole genomes of 809 primate samples across 233 primate species, which included samples from Great Ape Genome project (*19*) and other previous studies (*20-26, 126*). Among these, 783 samples passed quality evaluation and 26 samples failed QC. The major challenge of variant calling of these samples is that fewer than 60 primate species have genome builds available and the majority of samples are lack of reference genomes. Therefore, variant calling is performed using reference genomes from the closest species of primate samples. We aligned the sequencing data to 32 high-quality genome references (*110*), most of which are derived from long-read sequencing technologies. We adopted multiple hard filtering steps to remove low quality variants (*110*). As variants that are identity-by-state with human are of primary interest, we derived lift-over chain files between hg38 and each primate reference species from the multiple species alignment of 50 primate species and 8 mammal species (*110*) and lifted over the variants of primate samples to hg38. We examined the quality of coding variants by evaluating several QC metrics, including the number of stop-gained variants called per sample, the missense : synonymous ratios and the number of indels per sample.

##### *Machine learning classifiers of variants*

Due to the lack of reference genomes for the majority of primate species, we conducted a synthetic experiment to evaluate the impact of mapping to reference genomes of closely-related species on variant calling. We took a gorilla sample and mapped it to both hg38 and the gorilla genome (*110*). Mapping to gorilla and lifting-over to hg38 produced 6.6M variants after hard-filtering; while mapping to hg38 resulted in 45.4M variants. After hard filtering of low-quality variants and removing 38M fixed substitutions between hg38 and the gorilla genome, we observed 4.9M variants were shared between the two sets and 801.3K variants were called against hg38 but not called against gorilla. These variants are mostly false positives due to sequencing reads that were mapped to an incorrect region of the human genome or to regions of the human genome that are duplicated compared to the gorilla genome. It is essential to reduce these false positive variants substantially.

We developed machine learning classifiers to distinguish these false positive variants from the high quality variants for each primate sample. To train the classifiers, we used multiple sequence features per variant: GC content, GC skewness and local composition complexity within +/- 100bp of variants (*127*). In addition, we extracted variant features directly from VCF files, such as allelic count, mapping quality, the p-value of Fisher's exact test to detect strand bias, variant quality by depth, the symmetric odds ratio to detect strand bias, and genotype quality. We included the read depth (DP) of the variant normalized by the mean coverage of the primate sample and the fraction of alternative allele read depth out of the variant coverage. We observed that existence of indels nearby substantially influences the quality of variants called, thus we indicated indels within +/- 5bp and 10bp of variants. We also considered variant context features, including the mean coverage of the flanking regions around the variant normalized by the mean coverage of the primate sample, e.g., within +/- 100bp or +/- 500bp of the variant. Additionally, we observed that false positive variants often reside in poorly-mapped regions, which tend to accumulate overcalled variants. We counted the number of heterozygote SNPs within the flanking regions of variants (within +/- 100bp or +/- 500bp), which are normalized by the median counts of variants within the same length regions of the sample. Likewise, we included the normalized counts of alternative homozygote variants within the flanking regions of variants.

We labelled the 801.3K false positives as poor-quality variants and the remaining 44.6M variants as good-quality, including the 38M fixed substitutions. We randomly sampled 80% of the variant set for training and the rest 20% for testing.  Various classification methods were evaluated, including random forest, logistic regression, and multi-layer perceptron network. Random forest classifiers outperformed other methods with higher area under receiver operating characteristic (ROC) curve. We then independently trained six random forest classifiers using six gorilla samples by repeating the steps above and generated predicted scores for each variant using the six classifiers. The averaged predicted scores are denoted as the RF score of the variant. We chose a stringent cut-off of $<0.05$ for RF scores to minimize the effect of poor-quality variant calling on the ClinVar and other analyses.

We next evaluated the impact of applying the trained RF classifiers to other gorilla samples, which also showed comparable area under ROC curve. We then assessed whether the trained RF classifiers can be applied to other species pairs. Because reference genomes are unavailable for the majority of our species, we cannot directly assess the number of false positives when mapping to the close species. Instead, we designed several experiments to evaluate the accuracy of the gorilla-trained classifier that make use of the reference genomes we had in hand. First, we mapped a gorilla sample to the chimpanzee genome and applied the classifiers previously trained between gorilla and human. Second, we chose another pair of primates, rhesus macaque and baboon, to test the performance of the same classifiers. Last, we took human samples from an independent data set, Platinum Genomes project (*128*) and mapped those to chimpanzee and gorilla genomes. We also trained a second random forest classifier using the human and chimpanzee pairs and tested the performance of trained classifiers on human and gorilla pairs. All the results show the classifiers trained on one pair of closely-related species are generally applicable to another pair of closely-related species with comparable area under ROC.

***Cascaded variant filtering***

In addition to random forest filtering, we performed extra variant filtering steps. First, because we are interested in learning about pathogenicity of coding variants in humans, we removed variants that fell in codons where neither the reference nor the alternative allele resulted in a codon that matched the human codon at that position. Interestingly, this codon-match indicator also naturally reflects the clustering of these primate reference species into four major groups, great apes, Old World monkeys, New World monkeys and lemurs / tarsiers, as shown in the heatmap of Fig. S1A. Requiring codon match between primate and human genomes eliminated more than 50% of stop-gained variants in our primate dataset.

Next, we applied a series of gene-specific filtering steps to reduce the poor-quality primate variants in samples of each primate reference species. We excluded variants falling in primate transcripts carrying annotation errors compared with human transcripts, such as those with incorrect start codons or splicing donor and acceptor sites, which implies that transferring the human annotation directly to primates may have been problematic for these transcripts. We also removed variants in primate transcripts carrying in the middle of the sequence stop-gained variants that were not observed in the list of gnomAD protein-truncating variants.

We compared the distribution of variant random forest scores of a gene with the exome-wide distribution of variant RF scores and removed all variants in genes with a skewed distribution (Wilcoxon rank sum test p-value < 1e-20). We also merged the variants from all the samples mapped to a specific reference species and performed the Hardy-Weinberg equilibrium test for variants in primate reference species with at least seven samples. We then removed variants in the genes that carry any variants with excessive heterozygosity which also deviate from the Hardy-Weinberg proportions with p-value < 0.05.

In order to identify and exclude duplicated regions in primate genomes, we developed a unique mapping filter. First, we removed low quality sequencing reads of primate samples by filtering out reads with mapping quality < 20 from sample BAM files; then, we mapped the remaining reads to both hg38 and the relevant primate reference genome. We divided all reference genomes into 1kb bins and identified the best-mapped region in primate genomes as where the largest fraction of reads from one hg38 1kb bin are mapped (if reads from one hg38 1kb bin are mapped to two consecutive primate 1kb bins, the two bins are merged into one region). The fraction of reads from the hg38 bin that fall into the best-mapped region of the primate genome is the unique mapping score for that bin for each sample. By averaging the unique mapping scores across all the samples mapped to a specific reference species, we generated the unique-mapper (UM) score which applies to all the variants of the reference species that fall in the specific 1kb hg38 bin. For

ClinVar analyses, we chose a stringent cut-off of >90% for this score to ensure the one-to-one mapping between human and primate genomes.

These variant filters effectively reduced the number of stop-gained variants per primate sample to be close to the average number of stop-gained variants of human samples from Platinum Genomes project (*128*) (shown in Fig. S1B). The missense : synonymous ratios (MSR) gradually decreased after applying each of these filtering steps (Fig. S1C). In contrast, the missense : synonymous ratios of those excluded variants tend to be well above 1.0, implying they are either potentially deleterious or unreliably called. In addition, low-quality indels have also been substantially reduced (Fig. S1D).

***Mammal polymorphisms***
For mammal polymorphisms, we inherited the dbSNP variants of orangutan, rhesus, marmoset, cow, pig, mouse, goat, chicken and zebrafish from our previous paper (*17*) and lifted those over to hg38. We then excluded variants that failed codon-match requirement between hg38 and other species genomes. 109,732 unique missenses among the good quality variants of orangutan, rhesus, and marmoset were included in the training data set of PrimateAI-3D.

For each of primate, mammal and other vertebrate species, we computed a depletion metric following our previous paper (*17*), which measures the decrease of MSR of gnomAD common variants which are identical-by-state with other species, compared to the MSR of orthologous rare variants (Fig. S5).

***Evaluation of fraction of common variants in primate polymorphisms***
Due to that the averaged sample size per primate species is 2.5, we investigated the impact of small sample size on the fraction of common variants (allele frequency > 0.1% in each primate species) in the primate polymorphisms.

We then used the gnomAD allele frequencies of human common variants to simulate allele frequency spectra of primates at various sample sizes. For each primate species, we sampled genotypes according to gnomAD allele frequencies assuming the sample size is identical to that of the primate species. The fraction of gnomAD common variants discovered was averaged across 100 simulations for that specific sample size. We pooled the variants across the 233 simulated primate species and estimated the fractions of common variants for missense and synonymous variants separately, which are shown as allele frequency spectra in Fig. S2. According to this simulation, it is estimated that 95.1% of observed variants (>95.1%) are common variants (>0.1%), ~ 3% of synonymous variants in primates are rare (allele frequency < 0.1%) while ~ 94% of primate missense variants are common. Since the human allele frequency spectrum would be expected to have a larger fraction of rare variants than most other primate species due to the recent exponential expansion of human population size, the actual proportion of primate variants that are common (>0.1%) may be substantially higher than the 95.1% we estimated from simulations with gnomAD data.

From this simulation, we also obtained the average numbers of common synonymous variants at various sample size levels, which are used in the saturation analysis.

***Generation of training variant set for PrimateAI-3D***

For the benign variant set, we first included 126,873 unique common missense variants from human population data. To generate the primate polymorphism benign set, we relaxed our filtering criteria as deep learning algorithms naturally tolerate noise and benefit more from larger amounts of training data. We still removed variants falling in poor-quality genes or genes with poor annotations. However, we relaxed the unique mapper score cut-off to > 60% and the random forest score to < 0.17 and obtained 4,315,321 unique missense variants from primate sequencing. After merging human and primate missenses with missenses from dbSNP primates and variants from study on chimpanzee and bonobos (*129*), we obtained 4,514,581 unique missenses for the benign set in total.

All possible missense variants were generated from each base position of canonical coding regions by substituting the nucleotide at the position to the other three nucleotides. We excluded variants falling in start or stop codons, resulting in 71,166,190 all possible missense variants. After removing 4,514,581 benign variants and 6,207,640 human rare variants, 60,443,969 variants with unknown significance were left.

***ClinVar analysis of polymorphism data for human, primates, mammals, and other vertebrates***

To examine the clinical impact of variants that are identical-by-state with primate species, we downloaded the release variant summary for the ClinVar database (ftp://ftp.ncbi.nlm.nih.gov/pub/clinvar/vcf_GRCh38/variant_20210328.vcf.gz released on 28-March-2021) (*4*). The database contained 872,942 variants on the hg38 genome build, of which 753,511 were single nucleotide variants. Synonymous and stop-gained variants in the ClinVar database were excluded. Next, we required variants to have two-star review status or above, which includes "criteria provided, multiple submitters, no conflicts" and "reviewed by expert panel". We then removed those with unknown significance or conflicting interpretations of pathogenicity. We merged variants with Benign or Likely Benign annotations into a single category, as well merging variants with Pathogenic or Likely Pathogenic annotations. After these filtering steps, there were a total of 7,017 variants in the pathogenic category and 12,229 variants in the benign category (Fig. 1C).

We analyzed ClinVar variants that were identical-by-state with variation in primates, mammals and other vertebrates and compared to those present in human population, such as gnomAD database. A summary of the numbers of benign and pathogenic ClinVar variants that were present in great apes, Old World monkeys, New World monkeys, lemurs and tarsiers, more distant mammals, birds and fish is shown in Fig. 1F.

***Saturation of all possible human synonymous mutations with increasing number of primate populations sequenced***

We performed simulations to investigate the expected saturation of all ~22M possible human synonymous mutations by sampling common variants present in the 521 extant primate species. We considered various sample sizes for primates, including 10, 20, 50, 100, 200, 500 and 1000. In the previous section, we estimated the numbers of common synonymous variants observed in different sample sizes of humans via simulation.

For each primate species with each sample size, we simulated two times the number of common synonymous variants observed in human (allele frequency > 0.1%), because humans appear to have roughly half the number of variants per individual as other primate species (*130*). We assigned simulated variants based on the observed distribution of human common synonymous variants in the 192 trinucleotide contexts. For example, if 2% of human common synonymous variants were from the TAG>TCG trinucleotide context, we would require that 2% of the simulated variants were randomly sampled TAG>TCG mutations. This has the effect of controlling for the effects of mutational rate, genetic drift, and gene conversion bias, using trinucleotide context. The curves in Fig. S21 show the effects of varying the number of species, and the number of individuals per species on the saturation of the ~22 all possible human synonymous variants in the genome. With a small sample size of 10, more than 50% of human synonymous variants will be present in the 521 primate species. As the sample size per primate species increases to 1000, about 80% of human synonymous mutations will be covered if we can sequence all the extant primate species. With 521 primate species, all CpG transitions (100.0%) and non-CpG transitions (96.6%) would be observed, but only 62.3% of transversions would be covered, due to their much lower mutation rates. We note that this analysis assumes each species is a homogeneous population, which would underestimate the amount of variation due to subpopulation and subspecies structure; hence, saturating all transversions may still be likely within the 521 extant primate species.

**Derivation of primate mutation rates and observed/expected ratios per gene**

We estimated primate mutation rates using intronic sequences 50-200bp away from exons, where the impact of selection and other factors is minimal. We slid a window of 3 nucleotides along the intronic sequence in steps of one nucleotide and for each primate reference genome counted the number of each of the 64 possible trinucleotides. Next, we counted the number of variants with each of the 192 trinucleotide mutation contexts that were observed in all samples mapped to that reference genome. The ratio of the observed occurrences of each trinucleotide mutation context to the number of possible trinucleotides across the reference genome served as our estimate of the mutation rate.

Using these mutation rates, we computed the expected number of synonymous variants per gene for each primate reference species. We normalized our mutation rates to ensure that we expect the same total number of synonymous variants as are observed across the genome, although they may be distributed in different genes than are observed. To do this, we summed up the intronic mutation rates across the 192 trinucleotide contexts along the sequence of each gene and normalized this value by the total mutation rate of the whole exome. Multiplying the normalized mutation rate per gene with the total number of observed synonymous variants in that same gene, we generated the expected number of synonymous variants per gene. We then assessed the quality of primate mutation rates by computing the Spearman correlation between the observed and expected numbers of synonymous variants. The Spearman correlations vary from 0.295 to 0.925 among primate reference species due to the numbers of samples mapped to reference species range from 1 to 169.

Next, we evaluated multiple approaches to aggregating those mutation rates across primate reference species. First, we took the median of mutation rates across all the 31 non-human primate reference species. Second, as the mutation rate of one species is in practice interpreted as the probability of observing one mutation at a base position with one specific trinucleotide context, we computed the probability of observing at least one mutation across 31 reference species via a Binomial model for a specific base position with a trinucleotide context. Third, we calculated the same probability as the second approach except that at the base position that failed codon-match requirement, the mutation rate was assigned to zero. Last, we selected nine primate reference species with more than 20 samples and the Spearman correlation between the observed and expected numbers of synonymous variants > 0.75, including Aotus nancymaae, Ateles fusciceps, Cebus albifrons, Cercopithecus mitis, Lemur catta, Macaca mulatta, Pithecia pithecia, Rhinopithecus roxellana, and Papio anubis.  We took the median of mutation rates generated from these nine species to represent the primate mutation rates.

For each set of aggregated mutation rates, we generated the expected numbers of synonymous variants per gene across all the primate samples and took the ratio between the observed and expected numbers of synonymous variants to produce the observed/expected (O/E) ratio of synonymous for each gene. We multiplied each type of aggregated primate mutation rates with their specific O/E ratios of synonymous variants of genes to produce a new set of mutational probabilities of variants. Next, we evaluated the four sets of aggregated mutation rates and these four sets of adjusted mutational probabilities using the Spearman correlation between the mutational probability of a variant and the indicator of its presence in primate variants or not across all possible synonymous variants in the exome. The median mutation rates of nine primate species adjusted by the O/E ratios of genes achieved the highest Spearman correlation of 0.414. These adjusted primate mutation rates also outperform directly applying gnomAD mutation rates

(*29, 47*) with the Spearman correlation of 0.367, implying that primates have different mutational preference from humans.

Likewise, we computed the expected number of missense variants per gene across all primate species using this optimal set of primate mutation rates, which was normalized using the identical correction factor of synonymous variants (Fig. 2B). We computed the O/E ratios for missenses per gene (Fig. 2C). We then multiplied this best set of primate mutation rates by the O/E ratios of synonymous to generate mutational probabilities for each of all possible missense variants, which are used to select the matched set of variants with unknown significance for the PrimateAI-3D training.

In comparison with human data, we adopted the gnomAD mutation rates (*29, 47*) and variants, and computed the observed and expected numbers of synonymous and missense variants using the similar approach, as well as the O/E ratios of genes for synonymous and missense variants, respectively, shown in Fig. 2B and 2C.

**Identifying differential selection between humans and primates**

We sought to develop a model to 1) quantify the broad-scale similarity of natural selection between humans and primates and 2) identify genes evolving subject to remarkably different selective pressure in humans compared to primates. Because our strategy of mapping to divergent reference genomes means that some observed variants within a primate species are actually fixed differences between that species and the reference to which it was mapped, we based our estimates of selection on the number of segregating missense variants in each gene per primate species (i.e., we excluded variants that were carried on all chromosomes sampled from a given species). This ensures that we do not underestimate selection in the primate samples, because fixed variants are more neutral than segregating variants. Nonetheless, the number of segregating missense variants is shaped in complex ways by both sampling and demographic forces, we took a two-pronged approach to tackling this question. First, we built an explicit population genetic model to model selection across primates. Second, we developed a Poisson Generalized Linear Model to robustly detect genes that are differentially selected between humans and primates.

***Explicit population genetic model of selection***

Our explicit population genetic model proceeds through two phases: first, we model the counts of synonymous segregating sites to learn a neutral background distribution of mutation rates per gene per species. Then, we apply that neutral background distribution to estimate the average selection per gene across species.

*General modelling framework*

We first established a neutral baseline for each species by fitting a model to the segregating synonymous variants in each species. We employed the Poisson Random Field model, under which the observed number of segregating sites is a Poisson random variable, with the mean determined by mutation, demography, selection, and sample size (*34*). For simplicity, we assumed an equilibrium (i.e. constant) demography for all species besides human; for human, we used Moments (*51*) to find a best fitting demographic history based on the folded site frequency spectrum of synonymous sites.

With a best fitting demographic model in hand, we let $X_{igk}$ be the number of mutations of type $k$ ($k = 0$ is synonymous, $k = 1$ is missense) in gene $g$ of species $i$, $\theta_{ig} = 4N_i\mu_g$ be the per site population scaled mutation rate, and $L_{gk}$ be the number of sites of type $k$ in gene $g$. We then use dadi (*50*) to compute $p_i(\gamma_{\mathrm{ig}})$ which can be interpreted by noting that $\theta_{ig}p_i(\gamma)$ is approximately the probability that a site in gene $g$ with population scaled selection coefficient $\gamma_{\mathrm{ig}} = 2\mathrm{N_i s_g}$ is segregating in a sample from species $i$.

Then, the distribution of $X_{ijk}$ is Poisson with mean $\theta_{ig}L_{gk}p_i(\gamma_{ig})$, i.e.

$$\mathrm{P}(\mathrm{X_{igk}} = \mathrm{x}|\theta_{ig}) = \frac{\left(\theta_{\mathrm{ig}}\mathrm{L_{gk}}\mathrm{p_i}(\gamma_{\mathrm{ig}})\right)^{\mathrm{x}}}{\mathrm{x}!}\mathrm{e}^{-\theta_{\mathrm{ig}}\mathrm{L_{gk}}\mathrm{p_i}(\gamma_{ig})}.$$

*Background neutral model to estimate per gene mutation rates*

We anticipated that due to a combination of true variation in mutation rate and data quality across the genome, different genes would have a different effective per base-pair mutation rate. Although we could have used the estimated mutation rates from earlier work, we wanted to create a robust estimate with very few parameters per species. To accommodate this, we adopted a Gamma distributed prior on $\theta_{ig}$, and applied it to synonymous sites (i.e., $k = 0, \gamma_{ig} = 0$) and integrated over it to result in a scaled negative-binomial distribution.

$$P(X_{ig0} = x_0) = \int_0^\infty \left( \frac{\left(\theta_{ig} L_{g0} p_i(0)\right)^{x_0}}{x_0!} e^{-\theta_{ig} L_{g0} p_i(0)} \right) \left( \frac{\beta^\alpha}{\Gamma(\alpha)} {\theta_{ig}}^{\alpha-1} e^{-\beta\theta_{ig}} \right) d\theta_{ig}$$

$$= \frac{L_{g0} p_i(0)}{x_0!} \frac{\beta^\alpha}{\Gamma(\alpha)} \frac{\Gamma(\alpha + x)}{\left(\beta + L_{g0} p_i(0)\right)^{\alpha + x_0}}$$

To account for the impact of GC content on mutation rate, we parameterized the gamma distribution mean as a log-linear function of GC content,

$$log(\mu_g) = m_0 + m_{GC} GC_g,$$

$$log(\sigma) = sd_0.$$

from which we can compute gene-specific α and β parameters,

$$\alpha_g = \frac{\mu_g^2}{\sigma^2}$$

$$\beta_g = \frac{\mu_g}{\sigma^2}.$$

We then optimized the parameters $m_0$, $m_{GC}$, and $sd_0$ by maximum likelihood in each species to learn the background distribution of mutation rates.

Given our optimized parameters, we can compute a posterior distribution on the mutation rate per gene,

$$P(\theta_{ig} | X_{ig0} = x_0) = \frac{\left(\beta_g + L_{g0} p_i(0)\right)^{\alpha_g + x_0}}{\Gamma(\alpha + x_0)} \theta_{ig}^{\alpha_g + x_0 - 1} e^{-\left(\beta_g + L_{g0} p_i(0)\right)\theta_{ig}}$$

which is simply a gamma distribution with parameters ${\alpha_g}' = \alpha_g + x_0$ and ${\beta_g}' = \beta_g + L_{g0} p_{i(0)}$.

Fig. S7A shows a histogram of the average population scaled mutation rate across species.

*Modelling selection across species*

With the parameters $m_0$, $m_{GC}$, and $sd_0$ in hand, we can parameterize the distribution of the number of segregating nonsynonymous sites given a selection coefficient. To generate the expected number of segregating sites given a selection coefficient, we used dadi (*50*) to generate $p_i(\gamma)$ across a grid of population scaled selection coefficients, from 2Ns = 0 to 2Ns = 10000. We assumed further that every nonsynonymous mutation in a gene shares the same population scaled selection coefficient, $\gamma_{ig}$.

We then took the posterior distribution of $\theta_{ig}$ estimated from the synonymous sites as the distribution of mutation rates for nonsynonymous sites to obtain

$$\mathrm{P}\big(\mathrm{X_{ig1}} = \mathrm{x_1}|\gamma_{ig}\big)$$
$$= \int_0^\infty \left(\frac{\big(\theta_{ig}L_{g1}p_i(\gamma_{ig})\big)^{x_1}}{x_1!}e^{-\theta_{ig}L_{g1}p_i(\gamma_{ig})}\right)\left(\frac{\big(\beta_g + L_{g0}p_i(0)\big)^{\alpha_g+x_0}}{\Gamma(\alpha+x_0)}\theta_{ig}^{\alpha_g+x_0-1}e^{-(\beta_g+L_{g0}p_i(0))\theta_{ig}}\right)d\theta_{ig}$$
$$= \frac{L_{g1}p_i(\gamma_{ig})}{x_1!}\frac{\big(\beta_g + L_{g1}p_i(0)\big)^{\alpha+x_0}}{\Gamma(\alpha+x_0)}\frac{\Gamma(\alpha_g + x_0 + x_1)}{\big(\beta_g + L_{g0}p_i(0) + L_{g1}p_i(\gamma_{ig})\big)^{\alpha_g+x_0+x_1}}.$$

*Finding average selection coefficients via an EM-like procedure*

Noting that from population genomic data, the only thing that can be explicitly determined is the population-scaled selection coefficient, $\gamma_{ig} = 2N_i s_{ig}$, we devised a two-step procedure to control for demographic differences across species (i.e., differences in $N_i$). In principle $\gamma_{ig}$ can be different across species due to differences in either $N_i$ or $s_{ig}$, but we assumed that for the majority of genes, $s_{ig} \equiv s_g$ is identical across species, and the differences in $\gamma_{ig}$ are driven primarily by differences in $N_i$ . Our procedure can then be thought of as analogous to an EM algorithm, in which first we estimate $\gamma_{ig}$ separately for each species and gene, followed by estimating $N_i$ by assuming that $s_{ig}$ is identical across species, and finally using our estimated $N_i$ to produce estimates of $s_g$.

First, we inferred $\gamma_{ig}$ for each gene and species separately. This resulted in some information loss, because in species with small sample sizes, there may be many genes with 0 segregating missense variants, and hence result in an inference of an MLE $\gamma_{ig} = -\infty$. Nonetheless, for each species we obtained thousands of genes with an estimate of $\gamma_{ig}$. To control for some variation in estimated $\gamma$ across species, we additionally restricted to genes with $\gamma_{ig}$ between 10 and 1000.

We then assumed that for almost all remaining genes (1000-10000 depending on the species), the difference in estimated $\gamma_{ig} = 2N_i s_{ig}$ between species is due to the $N\_i$ being different, and that $s_{ig} \equiv s_g$ is identical across species for gene $g$. For each gene, we averaged estimated $\gamma_{ig}$ across species to obtain $\overline{\gamma_g} \approx 2\bar{N}s_g$, the average population scaled selection coefficient for gene $g$. Then, we computed $R_{ig} = \gamma_{ig}/\overline{\gamma_g}$, which is an estimator of $N_i/\bar{N}$ because $\gamma_{ig}/\overline{\gamma_g} \approx 2N_i s_g/2\bar{N}s_g = N_i/\bar{N}$. To account for noise among the outliers and the fact that $s_g$ is not truly identical across species, we estimated a single $R_i$ value per species by taking the median of $R_{ig}$ across all genes in species $i$.

Finally, we grouped species together to re-estimate $\overline{\gamma}_g$ by substituting $\gamma_{ig} \to R_i\overline{\gamma}_g$, which provides an approximation of $\gamma_{ig}$ from $\overline{\gamma_g}$ via $R_i\overline{\gamma}_g \approx N_i/\bar{N}2\bar{N}s_g = 2N_i s_g$. Then, we maximized the likelihood

$$L(\overline{\gamma_g}) = \prod_{i\in I}\mathrm{P}\big(\mathrm{X_{ig1}}|\mathrm{R_i}\overline{\gamma}_g\big)$$

to find the maximum likelihood estimate of $\overline{\gamma_g}$ across a group of species $I$. In the following, we take $I$ to be either the set $P$ of all non-human primates, or $I$ to be the singleton set $H$ containing just humans.

Fig. S7B shows our estimated $\overline{\gamma_g}$ as a function of the pooled missense : synonymous ratio among primates. The strong negative correlation is expected, as genes with higher missense : synonymous ratios have smaller estimated selection. To convert our estimates of $\overline{\gamma_g}$ to estimates of the selection coefficient, s, we divided by 2*10,000, which we take to be a typical effective population size among primates.

*Comparing human and primate selection*
Next, we compared human and primate constraint using this modelling approach. First, we computed a distribution of fitness effects (DFE) across genes in humans and primates by plotting a histogram of selection coefficients per gene in both our grouping of primates and our human data (Fig. S7C). We note that the much larger sample size of the human data compared to the primate data results in fewer outliers of strong selection, because almost every gene has missense variants in the human data, whereas in the primate data there are many genes observed with few to no primate missense variants.

To identify genes where human constraint is different from non-human primate selection, we developed a likelihood ratio test to test whether $s$ is significantly different between human and other primates by testing if $\overline{\gamma_g}$ is different between human and primate. Under the null model, $\gamma_{gh} = \overline{\gamma_g}$, so the likelihood is

$$\mathrm{L}_1(\overline{\gamma}_\mathrm{g}) = \left(\prod_{i\in P} P(X_{ig1}; R_i\overline{\gamma}_\mathrm{g})\right) \times P(X_h g1; R_h\overline{\gamma}_\mathrm{g})$$

where $P$ is the set of non-human primates and the subscript $h$ indicates human data. Under the alternative hypothesis, $\gamma_{hg} \neq \overline{\gamma_g}$ so the likelihood is

$$\mathrm{L}_2(\overline{\gamma}_\mathrm{g}, \gamma_{hg}) = \left(\prod_{i\in P} P(X_i; R_i\overline{\gamma}_\mathrm{g})\right) \times P(X_h; R_h\gamma_\mathrm{hg}).$$

This then forms the likelihood ratio test statistic,

$$\Lambda = 2(log\, L_2 - log\, L_1).$$

Note that under the null hypothesis $\gamma_{hg} = \overline{\gamma_g}$ we have that $L_1 = L_2$ and hence this represents a nested hypothesis test. Thus, under the null hypothesis that $\gamma_h = \overline{\gamma}$, the test statistic follows a $\chi^2$ distribution with one degree of freedom, $\Lambda \sim \chi^2(df = 1)$, by standard likelihood ratio theory.

Intuitively, this test determines whether $s_{hg}$ is significantly (P<0.05) different from $s_g$, because we already corrected for the effective population size of humans using $R_h$. Specifically, $R_h\gamma_{hg} \approx N_h/\overline{N}2\overline{N}s_{hg} = 2N_h s_{hg}$ while $R_h\overline{\gamma_g} \approx N_h/\overline{N}2\overline{N}s_g = 2N_h s_g$, so that if $\gamma_{hg} \neq \overline{\gamma_g}$, then $s_{hg} \neq s_g$.

Fig. S7D shows the relationship between $\overline{\gamma_g}$ and $\gamma_{hg}$, showing that there is a strong correlation. Colored points indicate genes that are significant according to the combined significance test described in the main text and subsequently.

*Comparison with alternative approaches*
We next assessed whether our population genetic modeling improved the correlation of selection estimates of our primate data with previous gene-constraint metrics in humans, including pLI (*28*) and s_het (*111*). We found that explicitly modeling the selection coefficient improves the correlation with these constraint metrics over the raw missense : synonymous ratio when using either primate data or human data (Fig. S8).

*Testing the model via simulation*

To verify the performance of our model, we performed population genetic simulations.

We generated data in line with our human and primate data by simulating data for 15000 genes. We used the number of synonymous and nonsynonymous mutations possible given the human reference genome for each gene, to create a realistic distribution of polymorphism in our simulations. For each gene, we drew the selection coefficient from a gamma distribution with mean 0.01 and standard deviation 0.01. We also drew the mutation rate per basepair for each gene from a gamma distribution with mean $5x10^{-8}$ and standard deviation $1x10^{-8}$. We then sampled from 213 species, with sample sizes matching those in our real primate data. For each species, we drew an effective population size from a gamma distribution with mean 10,000 and standard deviation 5,000.

To generate data for each species, we sampled from the Poisson random field model. Specifically, given an effective population size, mutation rate, sample size, and selection coefficient, the counts of sites from each gene in each species are sampled from a Poisson distribution as in the previous section. We then ran the inference pipeline inference on the simulated data, and Fig S9A shows that our inferred selection coefficients are very strongly correlated with the simulated selection coefficients, although they are somewhat upwardly biased for very weak selection of roughly $s \sim 1/10000$, which is expected based on the fact that the population scaled selection coefficient would be less than 1 on average, and thus have only minimal effects on segregating polymorphism. This shows that our model infers selection precisely from genome-scale data.

We then tested the power and calibration of the likelihood ratio test by simulating 2000 of the 15000 genes as having a different selection coefficient in humans compared to the rest of primates. To simulate the change in selection coefficient, we drew a random factor with a log-uniform distribution between 0.01 and 100 and multiplied the primate selection coefficient by that factor for each of the 2000 genes. We then performed the inference pipeline followed by the likelihood ratio test, with p-values corrected for multiple testing by the Benjamini-Hochberg procedure, as in the main text. Fig S9B shows that we obtain good control of the false discovery rate for genes with no change in selection between humans and primates, while obtaining greater power for larger shifts between human and primate.

***Poisson generalized linear mixed model***

Our population genetic model successfully modelled the similarity of selection between primates and humans, which we wanted to confirm with a less explicit model. To do so, we used a Poisson generalized linear mixed model, inspired by the model used in SnIPRE (*131*). We pooled polymorphic synonymous and missense mutations across all primates, and compared the pooled MSR to the MSR of human variants. Because these two quantities are related in a non-linear way, we used a two-step process. First, we fit a Poisson GLMM to the pooled primate data to estimate the depletion of missense variation at each gene. Then, we fit a second Poisson GLMM to the human data, controlling for the primate depletion estimates. This allowed us to estimate how much more or less depleted for missense variation each human gene is compared to what is expected based on primate genes. To control for noisily estimated missense : synonymous ratios, we only fit the Poisson GLMM to genes that had an average unique mapper score > 0.9 across all primate species and for which we had at least 100 synonymous variants pooled across primates.

*Primate GLMM*

To model primate variation, we set up a simple model in which there is a background mutation rate and the impact of GC content as fixed effects, and random effects accounting for gene-specific mutation rates and the depletion of nonsynonymous variation. Recalling from the population genetic model that $X_{igk}$ is the number of mutations of type $k$ ($k = 0$ is synonymous, $k = 1$ is missense) in gene $g$ of species $i$, we summed over all non-human primates to get the total number of variants of type $k$ in gene $g$ across non-human primates,

$$X_{gk}^{(P)} = \sum_{i \in P} X_{igk}$$

where P is the set of non-human primates, and the superscript $(P)$ indicates that the value is over non-human primates and is not an exponent. Then, we modeled $X_{gk}^{(P)}$ as a Poisson GLMM,

$$X_{gk}^{(P)} \sim Poisson\left(\mu_{gk}^{(P)}\right)$$

where

$$log\left(\mu_{gk}\right) = \beta_0^{(P)} + \beta_1^{(\mathrm{P})}\mathrm{GC_g} + \delta_g^{(P)} + \epsilon_g \mathrm{k} + log\left(L_{gk}\right)$$

with $\beta_0^{(P)}$ being a fixed effect corresponding to background mutation rate, $\beta_1^{(P)}$ being a fixed effect that corresponds to the impact of GC content, $\delta_g^{(P)}$ a random effect corresponding to the discrepancy between the mutation rate of gene $g$ and the genome-wide background, $\epsilon_g$ a random effect corresponding to the deficit of missense variation, $k$ is an indicator of mutation type, and $L_{gk}$ the number of sites of type $k$ in gene $g$. Note that k here serves as an indicator of whether a site is a synonymous or a missense mutation.

We fit this model using the R package glmer (*52*).

*Human GLMM*

With the estimates of $\epsilon_g$ in hand, we built a model for the human data $X_{gk}^{(H)} \equiv X_{hgk}$, with

$$X_{gk}^{(H)} \sim Poisson\left(\mu_{gk}^{(H)}\right)$$

Where

$$log\left(\mu_{gk}^{(H)}\right) = \beta_0^{(H)} + \beta_1^{(H)} GC_g + \beta_2 \epsilon_g k + \beta_3 \epsilon_g^2 k + \beta_4 \epsilon_g^3 k + \delta_g^{(H)} + \eta_g k + log\left(L_{gk}\right)$$

Here, $\beta_1^{(H)}$, $\beta_2^{(H)}$, $\delta_g^{(H)}$, and $L_{gk}$ have the same interpretation as the primate model, just applied to human data. However, we now have additional fixed effects $\beta_2$, $\beta_3$, and $\beta_4$ that model a nonlinear relationship between the missense depletion in primates and the missense depletion in humans. Thus, our remaining random effect, $\eta_g$ can be thought of as the deviation of the observed depletion of missense variants in humans compared to what would be expected based on the primate depletion. In particular, $\eta_g < 0$ indicates that a gene has even fewer missense variants than would be expected based on primates, and is thus suggestive of stronger constraint in humans than in primates, while $\eta_g > 0$ indicates an excess of missense variants compared to expected based on primates, implying relaxed constraint in humans compared to primates.

To see if the model is able to capture the nonlinear relationship of human and primate MSR, Fig. S10 shows the log-scaled MSRs of both human and primate, along with the function $\beta_2 x + \beta_3 x^2 + \beta_4 x^3$ which models the relationship of primate MSR to human MSR. It is able to capture the nonlinearity near the edges of the MSR distribution much better than a linear fit.

*GLMM p-values*

To determine which $\eta_g$ values were significantly different from 0, and hence indicative of human having less or more constraint than non-human primates, we developed an approach to controlling for gene length by binning genes by gene length and creating Z-scores based on the $\eta_g$ within each bin. Intuitively, shorter genes are likely to have a large magnitude of $\eta_g$ by chance. Moreover, because we don't actually expect that selection is identical between humans and primates, we anticipate identifying genes for which $\eta_g$ significantly deviates from a background distribution.

Our procedure was to first bin genes into quantiles by the number of amino acids in the human reference genome, each bin consisting of 100 genes. We then computed a Z-score by bin by computing the mean and standard deviation of $\eta_g$ within each bin and standardizing each $\eta_g$ by bin-wise mean and standard deviation. We then computed p-values assuming that the Z-score follows the standard normal distribution $Z \sim N(0,1)$.

#### PrimateAI-3D Model

We developed a comprehensive deep learning algorithm called "PrimateAI-3D". It takes as input protein structures and fixed-species multiple sequence alignments. A method called voxelization captures the protein structure surrounding a target variant and the conservation of its amino acids across evolution. A 3D convolutional neural network takes this voxelized structure as input and converts it into predictions of pathogenicity. PrimateAI-3D is trained to integrate three diverse objectives: distinction between human and primate common variants and unknown human variants; prediction of acceptable alternative amino acids at a protein site after removing all atoms of the original amino acid ("fill-in-the-blank in 3D"); ranking of scores that follows the ranking produced by the PrimateAI language model and the variational autoencoder from EVE.

##### *Training data preparation*

*Protein structures*

We downloaded predicted human protein structures from the AlphaFold DB (June 2021) (*73*). 16,568 protein sequences in that database matched exactly to one of our 19,158 hg38 proteins. For the rest, we performed homology modelling: we created a BLAST (*132*) database from the sequences in AlphaFold DB and searched against it with the remaining 2,590 proteins. For 2,167 proteins, we found a sequence match with >80% sequence identity, >80% target sequence coverage and <75 residues not covered (for 1,073 of those 2,590 proteins, both sequence identity and target coverage were >99%). We applied homology modelling software Modeller (*116*) using the AlphaFold structures as templates to calculate structures that exactly matched the sequences of our target proteins. For the remaining 423 proteins with more than 75 residues not covered by an AlphaFold DB structure, we used HHpred (*74*) to predict a structure for the entire protein. Then we used both the AlphaFold DB and HHpred structures as templates in Modeller. This procedure covered another 384 proteins with structure, leaving 39 proteins that failed in HHpred or Modeller. All these 39 proteins were several thousand amino acids long and were excluded from our training dataset.

*Multiple sequence alignments*

PrimateAI-3D took in the multiz100 alignment from the UCSC database (*112, 113*) to calculate the evolutionary conservation of each human protein residue in a set of 100 vertebrate species, similar to PrimateAI (*17*). In addition, we also included the protein sequences derived from the recently released Zoonomia study which consists of whole-genome alignments of 241 mammalian species (*114*). Finally, we obtained alignments of 251 species that covered at least 75% of all human proteins in the human Jackhmmer alignments (*133*) that we generated to replicate EVE (*67*); section "Model evaluation"). We aligned the amino acid sequences of the three sets of species to produce an alignment of human proteome across 592 species by filling any missing parts with gaps.

*Protein voxelization and voxel features generation*

Protein voxelization is the process of converting sets of protein atomic coordinates into tensors that have the same shape for all sets of coordinates and that can then be used in a traditional machine learning device. A regular sized 3D grid of cubes ("voxels") is centered at the Cα atom of the residue with the target variant (Fig. S11) and each voxel captures its atomic and evolutionary environment. We evaluated different combinations of grid sizes of voxels and voxel sizes and selected a grid size of 7x7x7 voxels with a voxel size 2Åx2Åx2Å, which achieves the best performance.

For each voxel, we computed a vector composed of diverse features, which is explained in detail below.

Atomic distance profile

For each voxel, we recorded the shortest distance between the Cα of one type of residuals (e.g., Alanine residue in Fig. S11) and the center of the voxel. We repeated this procedure for other amino acids and obtained the shortest distance of the voxel center to each of 21 amino acid types (20 standard amino acids plus one amino acid representing all non-standard amino acids). We then recorded the shortest distance for Cβ instead of Cα atoms, resulting in 2*21=42 distance values for each voxel (Fig. S11), which is referred to as the atomic distance profile of a variant. Detailed procedure is explained in Supplementary Text section 1.

Structure quality features

Several features measure the confidence of the structure around each voxel, including an indicator whether the protein structure is present in AlphaFold DB as-is (i.e., with a perfect sequence match to one of our protein sequences) and an indicator whether we used Modeller with AlphaFold DB structures as templates (in case the match was not perfect). We also included residue-specific quality features such as the pLDDT from AlphaFold DB (Fig. S12).

Species-differentiable evolutionary profiles

In PrimateAI (*17*), the evolutionary profile of a target residue is the frequency of each amino acid in the multiz100 alignment. This implies that all species have the same contribution to the amino acid frequency profile, regardless of the genetic distance of a species to human or to other species, thus it is an unrealistic scenario.  Therefore, in PrimateAI-3D, we assigned a different weight to each species of the 592-way whole proteome alignments. We initialized each weight to be 1/592 at the beginning of training, but let each weight be differentiable. This means PrimateAI-3D learns by itself how important each of the 592 species in the MSA is in terms of contribution to human pathogenicity. In Supplementary Text section 3, we implemented this procedure in a convolutional layer $Conv_f$ that, for any target residue, takes a fixed-species multiple sequence alignment as input and outputs an evolutionary profile with 210 features. In order to merge this evolutionary profile with voxels, we obtained a mapping from each voxel to the sequence position of the residue that is closest to the voxel center across all protein atoms in the structure (the nearest neighbor; function defined in Supplementary Text; Fig. S12).

Other protein-specific features

We included the reference amino acid in every voxel as a 1-hot encoded vector with 21 features (Fig. S12). The reference amino acid is the amino acid at the target site in the human reference proteome. Furthermore, we added two binary indicators to every voxel that signal whether the atoms of the target residue had been removed before voxelization or not (section “Model training”; Supplementary Text for details).

##### *Model Architecture*

The first layer of the network performs unpadded 3D convolutions in the voxel feature dimension with a kernel size of 1x1x1 and 128 filters, followed by ReLU activation and batch normalization. This creates an output tensor of shape 7x7x7x128. We then repeatedly applied 3D convolutions with a kernel size of 3x3x3, valid padding, and 64 filters until the output tensor’s shape becomes 1x1x1x64, again each time followed by ReLU activation and batch normalization. In each such layer, the first three dimensions of the output are reduced by 2 (because of the valid padding). Then we flattened the tensor and added a final hidden dense layer with 64 hidden units (ReLU activation, batch normalization). The output layer consists of 20 units (one for each standard amino acid) and uses a sigmoid activation function.

***Model training***

PrimateAI-3D was trained to perform multiple tasks simultaneously (multi-task learning), which are described in detail below. Each task captures an alternative and unique aspect of pathogenicity. We experimented with multiple combination techniques, including transfer learning, but found that simultaneous optimization gives the best generalization performance, which is in line with the current trend in structure prediction models (e.g., AlphaFold2 (*72*) and RoseTTAFold (*134*) to combine various evolutionary and physical aspects of predicted structures in their loss functions).

<u>Human and primate variants</u>

We obtained 4.5 million benign missense variants from human and primate data. We sampled a matched set randomly from the genome, requiring the distribution of mutational probabilities of unknown variants to be identical to that of benign variants. Another difference from PrimateAI (*17*) was that we used a single label vector to represent all the variants at the same amino acid site. This means we predicted the pathogenicity of all alternative amino acids at a site in one forward pass, instead of only a single variant. Note that a single nucleotide substitution cannot generate all 20 amino acids and that the label vector for a residue always contains missing values. We therefore masked missing labels during loss calculations so that they never directly contribut to the gradient. Benign variants have a label value 0 and pathogenic variants have a value 1. We used mean squared error as the loss function for non-missing labels.

<u>Variants from fill-in-the-blank in 3D</u>

The generation of this variant set was inspired by the typical training procedure of language models (*81*). During voxelization, the voxel grid is centered on the Cα atom of a target residue and the distance profile is calculated using all atoms within the scanning radius of a voxel center. For fill-in-the-blank in 3D, we performed the same procedure, except that we removed all atoms of the target residue before calculating the distance profile or nearest neighbor mapping. In effect, all features specific to the target residue were removed from the input tensor to the network. Then we trained the network to pick the amino acid which may be acceptable at the target site. These acceptable / unacceptable amino acids from the multiple sequence alignments formed a second training set. Any amino acid that occurs in the MSA at the target site was considered acceptable (label 0), all others unacceptable (label 1).

<u>Variant ranks from language models</u>

Language models, such as EVE and PrimateAI language model, perform competitively on data sets evaluated on a per-gene basis, such as saturation mutagenesis assays. However, directly using prediction scores from these models as additional features to PrimateAI-3D failed to improve performance. This is due to that the major fraction of the variance in variant pathogenicity across the human proteome can be attributed to the variation of proteins or protein domains. For example, in the human and primate variant set, the ratios between the numbers of common and unknown variants are lower for more pathogenic genes. Similarly, in fill-in-the-blank training, we observed fewer species tend to carry mutations in more pathogenic genes, assuming a fixed set of species across the proteome. Language model scores, however, have not been calibrated well for this inter-protein (or protein domain) variation, thus are ignored by gradients from the previous two datasets with dominant effect on variance. Instead, we hypothesized that using language model scores as an additional target output could be helpful: first, neither of the two datasets capture epistatic patterns (unknown human variants can contain both pathogenic and benign variants; fill-in-the-blank labels are taken only from single protein positions in limited species without considering sequence context). Secondly, voxelized structures may be expressive enough to capture epistatic patterns by themselves. For example, epistatic interactions usually indicate residue-residue interactions in structure (*135*).

We used a rank loss function to incorporate language model scores as a third training dataset, and only computed it on variants from the same protein. More specifically, we first converted the scores from language models to ranks, separately for each protein. This means the ranks for each protein are within range (1,…,N), where N is the length of the protein. These ranks are the truth ranks, i.e. PrimateAI-3D is trained to produce scores that have the same ranks. The pairwise logistic loss from Pasumarthi *et al.*(*117*) measures the distance between two sets of ranks and produces a gradient that ultimately updates the model to predict scores that better match the truth ranks.

*Training procedure*

The dataset of human and primate variants covers 5.6M of 10.8M possible amino acid positions in human proteome. For each of the other two datasets (fill-in-the-blank in 3D and language model ranks), we sampled equally as many amino acid positions. This is primarily due to practical reasons. First, this allows each batch to have the same number of samples from each dataset (~33 with a batch size of 100), leading to more stable training in each dataset individually. Second, it controls the influence of each dataset solely via the sample weight in the loss function, instead of training sample size as an additional free parameter. Third, it keeps training and epoch times at reasonable levels. Last, there was no obvious performance benefit from allowing more samples.

For the language model ranks dataset, we additionally required that all 33 samples in a batch come from the same protein. This allows calculating rank losses not only for the same protein position, but across all samples from the same batch. The number of times that a protein was chosen for a batch was proportional to the length of the protein. In order to make our model robust against protein orientations, we randomly rotated the protein atomic coordinates in 3D before voxelizing a variant.

Model optimization was performed using Adam (*136*) from Keras 2.2.0 with default parameters and a learning rate of 0.001. From each of the three datasets, we sampled 20,000 hold-out protein positions for model validation. For each alternative model and epoch, we predicted these hold-out validation datasets, together with variants from *BRCA1* and *TP53* assays. This produced two area under ROC metrics (human and primate variants and fill-in-the-blank in 3D variants) and three Spearman rank correlation values (language model ranks and *BRCA1* and *TP53* assay ranks) for each epoch in each alternative model. We calculated the rank of each metric across all alternative models, e.g., we converted human and primate AUC values from 9 alternative models in a range (0.6, …, 1.0) to ranks (1, …, 9). Then, for each model, we averaged the ranks of the 5 datasets, and only kept the model with the highest average rank. Furthermore, we used this procedure to optimize the weights of each dataset in the loss function (0.5 for fill-in-the-blank in 3D, 1.0 for language model ranks, 2.0 for human and primate variants).

*Ensemble training and inference*

Once we found the optimal model and model parameters, we repeated this training procedure 40 times, each time with a different initial seed value. This generated 40 different models. In order to predict a variant, we calculated its pathogenicity score using all 40 models 10 times, each time with a different protein orientation. The average of these 400 scores was the final pathogenicity score for the variant.

#### PrimateAI Language Model

PrimateAI language model is a multi-sequence alignment transformer (*83*) for fill-in-the-blank residue classification, which was trained end-to-end on MSAs of UniRef-50 proteins (*118, 119*) to minimize an unsupervised masked language modelling objective (*81*). It outputs classification scores for alternative and reference residues, which serve as inputs to the PrimateAI-3D rank loss.

Traditionally, fill-in-the-blank MSA transformers simultaneously classify multiple masked locations in MSAs during training. Higher numbers of mask locations can add more MLM gradients that inform optimization, thereby enabling a higher learning rate and faster training. However, fill-in-the-blank pathogenicity prediction is fundamentally different from traditional MLM as classification at a mask location depends on predicted values of residues at other mask locations. The classification scores may often be the averages of conditional predictions over all possible combinations of residues at other mask locations. PrimateAI LM avoids this averaging by revealing tokens at other mask locations before making predictions. Our model achieves state-of-the-art clinical performance and denoising accuracy whilst requiring 50x less computation for training than previous MSA transformers.

##### *Preparation of MSA datasets*

We created an MSA for each sequence in a UniRef-50 database (March 2018 version) (*115, 118*) by searching a UniClust30 (*137*) database (October 2017 version). Then an MSA dataset containing 26 million MSAs was created using the protein homology detection software HHblits version 3.1.0 (*138*). Default settings were used for HHblits except that we set the number of search iterations (-n) to 3. This replicates the approach to generating MSAs to train MSA transformer (*139*). In addition, we generated a set of MSAs for 19,071 human proteins using HHblits following the procedure above.

Next, we excluded UniRef-50 MSAs whose query sequences carry rare amino acids, retaining those containing the 20 most abundant residues only. To further simplify our data pipeline, we filtered non-query sequences in the MSA to those that only contain the 20 most common residues and gaps, which represent deletions relative to the query sequence. As input MSAs to PrimateAI LM have a fixed size of 1024 sequences, we randomly sampled up to 1023 non-query sequences from the filtered sequences if MSA depth is larger than 1024. If MSA depth is below 1024, we padded the MSA with zeros to fill the input.

We then applied a periodic mask pattern with a stride of 16, which covers an amino acid position of interest in the query sequence, to MSAs. Using a fixed mask pattern ensures consistent computational requirements for mask revelation discussed below. The position of interest was randomly sampled from all positions in the query sequence during training or chosen by a user during inference. To maximize information about the position of interest, we tried to select a cropping window with a size of 256 residues where the position of interest is at the center. However, the cropping window may be shifted if the position of interest is near the edge of an MSA to avoid padding zeros and increase information about the position of interest. If the query sequence is shorter than the PrimateAI LM cropping window, zeros were padded to fill the window size. Illustration of cropping, masking, and padding of MSAs input to PrimateAI LM is shown in Fig. S23.

We followed AlphaFold 2 (*72*) to assign a smaller probability, $p_{\mathrm{sample}}$, to an MSA being sampled during training if the protein length, $L$, is shorter,

$$p_{\mathrm{sample}} \propto \frac{\max(\min(L, 512), 64)}{512},$$

to rebalance the distribution of lengths for UniRef-50 proteins used for training and human proteins, and to avoid computation being wasted on padding. We also adjusted the probability of sampling non-query sequences to be included in the first 32 sequences of an MSA, where the fixed mask pattern is applied, to penalize the occurrences of gaps in those sequences. The probability, $p_{\mathrm{mask}}$, of a non-query sequence being masked decreases with increasing number of gap tokens, $N_{\mathrm{gap}}$,

$$p_{\mathrm{mask}} \propto \frac{\left(L - N_{\mathrm{gap}}\right)^2}{L^2}.$$

Down sampling of sequences with lots of gaps reduces the fraction of missing data in MSAs.

***Model Architecture***

*MSA Embedding*

We first embedded MSA input for PrimateAI LM, shown in Fig. S24, and applied a fixed mask pattern to the first 32 sequences of MSAs. The MSA tokens were encoded by learned 96-channel embeddings, which were summed with learned 96-channel position embeddings for residue columns before layer normalization (*140*). To reduce computational requirements, embeddings for the 1024 sequences in MSAs were split into 32 chunks, each containing 32 sequences, at periodic intervals along the sequence axis. These chunks were then concatenated in the channel dimension and mixed by linear projection.

*MSA Transformer*

Embedded MSAs were propagated through 12 axial attention blocks shown in Fig. S13. Each axial attention block consists of residuals that add tied row-wise gated self-attention, column-wise gated self-attention, and a transition layer, shown in Fig. S13. The self-attention layer has 12 heads, each with 64 channels, totaling 768 channels, and transition layers project up to 3,072 channels for GELU activation (*141*). The adoption of axial gated self-attention was inspired by AlphaFold 2's Evoformer (*72*). The main change is that we used tied attention (*139*) in PrimateAI LM axial attention layer (Fig. S25), instead of triangle attention in AlphaFold 2. Tied attention is the sum of dot-product affinities, between keys and values, across non-padding rows, followed by division by the square root of the number of non-padding rows, which reduces computational burden substantially.

*Mask Revelation*

Mask revelation, shown in Fig. S26, reveals unknown values at other mask locations after the first 12 axial attention blocks. It combines the updated 768-channel MSA representation with 96-channel target token embeddings at locations indicated by a Boolean mask which labels positions of mask tokens. The Boolean mask, which is a fixed mask pattern with stride 16, is applied row-wise to gather features from the MSA representation and target token embedding at mask token locations. Feature gathering reduces row length from 256 to 16, which drastically decreases the computational cost of attention blocks that follow mask revelation in Fig. S26. For each location in each row of the gathered MSA representation, we concatenated the row with a corresponding row from the gathered target token embedding where that location is also masked in the target token embedding. The MSA representation and partially revealed target embedding are concatenated in the channel dimension and mixed by linear projection.

After mask revelation, the now-informed MSA representation is propagated though residual row-wise gated self-attention and transition layers shown in Fig. S13. The attention is only applied to features at mask locations as residues are known for other positions from the MSA input to PrimateAI LM. Thus, attention only needs to be applied at mask locations where there is new information from mask revelation. After interpretation of the mask revelations by self-attention, a masked gather operation collects features from the resulting MSA representation at positions where target token embeddings remained masked. The gathered MSA representation is translated to predictions for 21 candidates in the amino acid and gap token vocabulary by an output head shown in Fig. S27.

##### *Model Training*

*Loss Function*

PrimateAI LM was trained end-to-end on MSAs for UniRef-50 proteins to minimize a weighted masked language modelling loss,

$$L_{MLM} = -w_{\text{length}} w_{\text{mask}} \sum_{i,j \in M} \log(\mathrm{p}_{ij})$$

where $M$ is the set of positions where MSA input tokens are masked, and probabilities, $p_{ij}$, are computed from PrimateAI LM outputs by softmax normalization, $p_{ij} = s\text{oftmax}(l_{ij})$, of logits, $l_{ij}$, output by PrimateAI LM. Softmax normalization over the amino acid vocabulary is applied independently per position, $j$, in each sequence, $i$. Since query sequences do not contain gap tokens, query sequence gap token logit values are changed to $-10^5$. Loss weights are higher for longer proteins, thus we designed this weight,

$$w_{\text{length}} = \min\left(L^{\frac{1}{2}}, 64\right),$$

to adjust for the effect of a small portion of longer proteins when taking a single fixed-sized crop from their MSAs. Weights are also higher for MSAs with a lower number of masked positions, $N_{\text{mask}}$,

$$w_{\text{mask}} = N_{\text{mask}}^{-\frac{1}{2}},$$

to rebalance contributions from MSAs with various depths and padding.

*Optimizer*

PrimateAI LM was trained for four days on four A100 graphical processing units (GPUs). Optimizer steps are for a batch size of 80 MSAs, which is split over four gradient aggregations to fit batches into 40 GB of A100 memory. PrimateAI LM was trained with the LAMB optimizer (*82*) using the following parameters: $\beta_1 = 0.9$, $\beta_2 = 0.999$, $\epsilon = 10^{-6}$, and weight decay of 0.01.

Gradients are pre-normalized by division by their global L2 norm before applying the LAMB optimizer. Training was regularized by dropout (*142*) with probability 0.1, which was applied after activation and before residual connections, as shown in Fig. S25. Axial dropout (*72*) was applied in self-attention before residual connections: post-softmax spatial gating in column-wise attention is followed by column-wise dropout, and post-softmax spatial gating in row-wise attention is followed by row-wise dropout.

PrimateAI LM was trained for 100,000 parameter updates. The learning rate is linearly increased over the first 5,000 steps from $\eta = 5 \times 10^{-6}$ to a peak value of $\eta = 5 \times 10^{-4}$, and then linearly decayed to $\eta = 10^{-4}$. We applied automatic mixed precision (AMP) to cast suitable operations from 32-bit to 16-bit precision during training and inference (*143*), which increases throughput and reduces memory consumption without affecting performance. In addition, we used a Zero Redundancy Optimizer to reduce memory usage by distributing optimizer states across multiple GPUs (*144*).

*Ensemble*

An ensemble of six PrimateAI LM networks was trained with different random seeds for training data sampling and model parameter initialization. Their top-1 accuracies during training are shown in Fig. S28 for mask locations in the query sequence and all sequences in UniRef-50 MSAs. Top-1 accuracy for the query sequence is much lower than for all sequences as the query sequence does not contain gap tokens, which are easier to predict than residues as they often form long and contiguous segments in MSAs created with HHblits. PrimateAI LM accuracy on query sequences was steadily improving at the end of training. However, PrimateAI LM was trained for no more than $10^5$ iterations due to computational cost.

***Inference and pathogenicity score***

PrimateAI LM fill-in-the-blank predictions are provided for locations of interest at every site in 19,071 human proteins, totaling predictions for 2,057,437,040 variants at 108,286,160 positions. Each prediction is made by our ensemble of six models, with each model contributing at least four inferences with different random seeds for sampling and ordering of sequences in human MSAs. Inferences logits were averaged by taking means of predictions grouped by random seed, and then taking the mean of the means. Each inference for 19,071 human proteins takes nearly 7 days on an A100 and, in total, inference by the ensemble takes nearly 200 A100 days.

Pathogenicity prediction of a variant is traditionally evaluated according to the relative logit of the variant residue compared to the one of the reference amino acid, i.e., $\log(p_{alt}) - \log(p_{ref})$, where $p_{ref}$ and $p_{alt}$ are the reference (ref) and alternative (alt) probabilities obtained from the ensembled logits. The probabilities are normalized over all possible residues disregarding the gap token, such that $\sum_r p_r = 1$ with probability $p_r$ of the r$^{th}$ residue obtained from the ensembled logits. The log difference captures how unlikely the variant amino acid is compared to the reference amino acid. However, the score does not consider the prediction of the other 18 possible amino acids, which contain information about the language models internal estimate of protein site conservation as well as convergence of the language model. We used the entropy evaluated over amino acid predictions $S = -\sum_r p_r \log(p_r)$ with probability $p_r$ of the r$^{th}$ residue to capture a variant agnostic site-dependent contribution to the pathogenicity score. Specifically, a score, $s_{alt}$, for residue alt at a given site is given by the usual log difference of the alt and reference ref logit at that site minus the entropy over amino acids at the given site, i.e.

$$s_{alt} = \log(p_{alt}) - \log(p_{ref}) - S.$$

The entropy term is small whenever the probability over all amino acids is dominated by a single term and large whenever the model is uncertain about the residues and assigns multiple residues high values. Physically, in this case the site is associated with little conservation and likely to mutate. This should lead to less pathogenic signal.  Adjusting the scores by entropy incorporates a model internal estimate of amino acid conservation. A given log difference between residue and reference will be considered as more pathogenic whenever it is associated with a highly conserved site. The score adjustment additionally incorporates the lack of convergence associated with a heavily undertrained model.

#### Model Evaluation

##### *Evaluation datasets*

*Saturation mutagenesis assays*

We compared model performance using deep mutational scanning assays for the following 9 genes: amyloid-beta (*102*), *YAP1 (96), MSH2 (120), SYUA (101), VKOR1 (121), PTEN (99, 100), BRCA1 (122), TP53 (123),* and *ADRB2* (*124*). We excluded from the evaluation analysis a few assays of the genes for which the predication scores of some classifiers are unavailable, including *TPMT (99), RASH (145), CALM1 (146), UBE2I (146), SUMO1 (146), TPK1 (146),* and *MAPK1* (*147*). We also excluded assays of *KRAS* (*148*) (due to different transcript sequence), *SLCO1B1* (*149*) (only 137 variants), and amyloid-beta (*150*) (duplicate of (*102*)). We evaluated model performance by computing the absolute Spearman rank correlation between model prediction scores and assay scores individually for each assay and then taking the mean across all assays. See Table S6 for per-assay rank correlations for each method.

*UK Biobank*

The UK Biobank dataset (*79, 80*) contains 61 phenotypes across 100 genes. Evaluating on common variants of all methods reduces the number to 41 phenotypes across 42 genes. We calculated the absolute Spearman rank correlation between the predicted pathogenicity scores and the quantitative phenotype scores for each pair of gene/phenotype. Only gene/phenotype pairs with at least 10 variants were included in the evaluation (14 phenotypes across 16 genes). We also confirmed that our evaluation is robust to this choice of threshold.

*ClinVar*

We benchmarked model performance in classifying clinical labels of ClinVar (downloaded from https://ftp.ncbi.nlm.nih.gov/pub/clinvar/tab_delimited/variant_summary.txt.gz on September 19, 2021) missense variants (*4*) as benign or pathogenic. Both "benign" and "likely benign" labelled variants were considered benign, the same for "pathogenic" and "likely pathogenic" labelled variants (both considered pathogenic). To ensure high-quality labels, we followed (*67*) and only included ClinVar variants with 1-star review status or above (including "criteria provided, single submitter", "criteria provided, multiple submitters, no conflicts", "reviewed by expert panel", "practice guideline"). This reduced the number of variants from 36,705 to 22,165 for the pathogenic and from 41,986 to 39,560 for the benign class. Following EVE (*67*), we calculated the area under the receiver operating characteristic curve for each gene and then report the mean AUC across all genes.

*DDD / ASD / CHD de novo missense variants*

To evaluate the performance of the deep learning network in clinical settings, we obtained de novo mutations from published studies for intellectual disorders, including autism spectrum disorder (*88-94*) and developmental disorders (*85-87*). ASD contained 2,127 patients with at least one de novo missense mutation. Taken together, there were a total of 3,135 DNM mutations. This reduced to 517 patients with at least one DNM variant and a total of 808 DNM variants after requiring all methods had predictions for those variants. In DDD, 17,952 patients had at least one de novo missense variant (26,880 variants in total), reducing to 6,648 variants after requiring availability of predictions of all methods. We also obtained a set of DNM variants from patients with congenital heart disorders (*95*), consisting of 1,839 de novo missense variants from 1,342 patients (reducing to 564 variants after requiring availability of predictions of all methods). For all the three datasets of de novo variants from affected patients, we used a shared set of DNM variants from healthy controls, which contains 1,823 DNM variants from 1,215 healthy controls with at least one DNM variant and collected from multiple studies (*88-93*). It was reduced to 250 variants (235 patients) after requiring availability of variant prediction scores of all methods. For each disease set of DNMs, we applied Mann-Whitney U test to evaluate how well each classifier can distinguish the DNM set of patients from that of controls.

***Methods for comparison***

Predictions from other methods were evaluated using rank scores downloaded from the database for functional prediction dbNSFP4.2a (*84*). To avoid dramatic reductions in the number of common variants, we removed methods with incomplete sets of scores (methods with less than 67 out of 71 million possible missense variants in hg38), except Polyphen2 (*151*) due to its widespread adoption. We included the following methods (method abbreviation) for comparison: BayesDel_noAF (BayesDel) (*152*), CADD_raw (CADD) (*153*), DANN (*154*), DEOGEN2 (*155*), LIST-S2 (*156*), M-CAP (*157*), MutationTaster_converted (MutationTaster) (*158*), PROVEAN_converted (PROVEAN) (*159*), Polyphen2_HVAR (Polyphen2; due to better performance then Polyphen2 HDIV) (*151*), PrimateAI (*17*), Revel (REVEL) (*160*), SIFT_converted (SIFT) (*161*), VEST4 (*162*), fathmm-MKL_coding (fathmm-MKL; highest performance among the fathmm models for given benchmarks) (*163*).

ESM1v model (*164*) was not released as part of dbNSFP4.2a (*84*). Due to unavailability of full mutation effect predictions of the human proteome for this model, we used the pre-trained ESM1v weights downloaded from GitHub (https://github.com/facebookresearch/esm) and evaluated on all human protein sequences using the published code without any modifications.

***Applying EVE to more proteins***

In the original publication, EVE (*67*) is only applied to a small set of disease-associated genes in ClinVar. To generate our language model-based training data set, it is essential to expand the predictions of EVE to as many proteins as possible. Due to unavailability of EVE source code, we therefore applied a similar method DeepSequence (*165*) and converted DeepSequence scores into EVE scores by fitting Gaussian mixture models. We used an up-to-date version of UniRef100 (*115*), but otherwise followed the alignment depth and sequence coverage filtering steps described in (*67*). We achieved at least 1 prediction in 18,920 proteins and a total of 50.2M predicted variants out of 71.2M possible missense variants. To validate our replication, we evaluated the replicated EVE models using published variants from (*67*). We found that scores from the replicated EVE model results in comparable performance to the published EVE software on all benchmarking datasets, e.g., both methods achieve 0.41 mean absolute correlation on Assays and 0.22 mean absolute correlation for UKBB.

***Benchmarking PrimateAI LM against other sequence-only models for pathogenicity predictions***

PrimateAI LM falls into a class of methods only trained to model proteins sequences but performing surprisingly well as pathogenicity predictors. Despite not achieving the overall best performance by themselves, they make crucial features or components in classifiers incorporating more diverse data. Fig. S14 summarizes the evaluation performance of the PrimateAI LM against other such sequence-only methods for pathogenicity prediction: ESM1v (*164*), EVE (*67*), LIST-S2 (*42*), and SIFT (*48*). Our language model outperforms another language model ESM1v on all the testing datasets except assays using only 1/50$^{th}$ of the training time. This is particularly striking as PrimateAI LM does not rely on any fine-tuning on assays.

***Combining PrimateAI LM with EVE***

Language models are trained to model the entire universe of proteins. EVE (*67*) trains a separate model for each human protein and all similar sequences. This and the differences in model architecture and training algorithms suggest that the models extract distinct features from their input. Therefore, we expected that the scores from EVE and our language model to be complementary and that combining scores may result in improved performance. We found that simply taking the mean of their pathogenicity scores already performs better than any of the two methods alone. More elaborate combinations, e.g., using ridge regression, did not lead to any further improvements. The resulting performance is shown in Fig. S14, where the combined score leads to a performance gain of 6.6% (or 6.8%) in mean correlation on assays compared to the PrimateAI LM (or compared to replicated EVE), 1.4% (or 1.7%) improvement mean AUC on ClinVar and increases in P-value by 11% (29%) for DDD, 3% (26%) for ASD and 17% (23%) for CHD.

***Evaluations of PrimateAI-3D***

PrimateAI-3D performance is benchmarked against both supervised and unsupervised variant pathogenicity classifiers. We found that PrimateAI-3D consistently outperforms other classifiers on all evaluation datasets (Fig. 3D). Summary statistics of model performance across all the six evaluation datasets are provided in Fig. S15 and Table S3. When the results are averaged across benchmarks, PrimateAI-3D is far ahead of any competing classifier (Fig. S15). It appears the reason why other classifiers come close to PrimateAI-3D in individual benchmarks is because we evaluated nearly 30 other algorithms; akin to multiple hypothesis testing, by statistical chance the performance of an algorithm may be an outlier on one particular benchmark, but their lack of consistency in the other five benchmarks indicates regression back to the mean. As we show in Fig. 3D, the second-place algorithm is different in each of the six benchmarks. A detailed breakdown of the performance of PrimateAI-3D for each of the 42 genes in UKBB is provided in Table S4. A detailed summary of performance of PrimateAI-3D and other benchmark classifiers on the 9 assays considered is given in Table S6.

***PrimateAI-3D sensitivity and specificity on ClinVar***

The ClinVar mean per-gene AUC metric (Fig. 3; also used in EVE (*67*)) implicitly corrects for different biases in ClinVar and enables a fairer comparison to methods directly trained on clinical annotations. A complication of directly measuring sensitivity and specificity in the ClinVar database is that human expert annotations are concentrated in a handful of disease genes that have been most heavily studied, with 50% of the total pathogenic missense variants in ClinVar coming from only 1.8% of the protein-coding genes in the genome (Fig. S16). This bias in annotation results in some genes having severe imbalance in their number of benign and pathogenic mutations, with >90% of labeled variants in the gene being either pathogenic or benign. Other machine learning classifiers that have been trained on human annotation databases have inadvertently learned the bias in annotation and take advantage of this property to assign higher scores to variants in genes with a high fraction of pathogenic variants in ClinVar (Fig. S17). To measure each classifier's sensitivity and specificity without the influence of annotation bias, we rank-normalized classifier scores for variants within each gene, and excluded genes with >90% of variants being pathogenic or benign in ClinVar. PrimateAI-3D achieved a sensitivity of 84.7% and specificity of 84.1% (percentile threshold: 57.7), and its AUC of 0.919

was the best performance among all classifiers (Fig. S18), which was notable given that the other classifiers (with the exception of EVE, PROVEAN, SIFT, LIST-S2) had been trained either on ClinVar or highly overlapping annotation databases such as HGMD (*166, 167*).

***Using PrimateAI-3D to improve ClinVar***

To assess the utility of PrimateAI-3D for revising ClinVar annotations, we compared a snapshot of the ClinVar database from September 2017 with the current database, and asked whether PrimateAI-3D scores were predictive of variants whose annotations had been revised in the interim. We defined the most confident *X*% PrimateAI-3D predictions as the *X*/2% variants with the highest predicted pathogenicity merged together with the *X*/2% variants with the lowest predicted pathogenicity. This implicitly converts the continuous PrimateAI-3D score into a binary benign/pathogenic classification. We downloaded ClinVar September 2017, reduced it to benign or pathogenic missense variants and only kept variants from genes with at least 20 variant annotations (30,295 variants). Then we looked up the annotations of those variants in ClinVar 2021. We found that 4,850 of the 30,295 variants (16%) had changed or lost their clinical annotation by 2021. Filtering the 30,295 variants to those that are also in the top 10% most confident PrimateAI-3D predictions, 2,905 variants remain. For 55/2,905 (2%) variants, ClinVar and PrimateAI-3D disagree. 29/55 (53%) variants changed ClinVar labels between 2017 and 2021. This is a 5 times higher fraction than for variants that agree between PrimateAI-3D and ClinVar (273/2755, 10%; Fisher's test P-value=$10^{-24}$) and indicates that at least half of all annotations that disagree between high-confidence PrimateAI-3D and ClinVar will change annotation in ClinVar over time. We repeated this analysis with different confidence thresholds for PrimateAI-3D (Fig. S19). Among the variants that were annotated as pathogenic in 2017, the top 10% with the lowest (most benign) PrimateAI-3D scores were 4-fold more likely to have had their ClinVar annotation consequence changed in the interim ($P < 10^{-3}$). Conversely, among the variants annotated as benign in 2017, the top 10% with the highest (most pathogenic) PrimateAI-3D scores were 6-fold more likely to have had their annotation changed ($P < 10^{-18}$) (Fig. S19). Even using the top 75% of all PrimateAI-3D predictions, there remains a 2-fold increase of ClinVar label changes among variants that disagree between ClinVar and PrimateAI-3D. In summary, discordance between PrimateAI-3D and ClinVar indicates a significantly elevated chance of annotation error in ClinVar and can be used to increase confidence in ClinVar annotations.

**Candidate gene discovery**

We tested enrichment of de novo mutations in genes by comparing the observed number of DNMs to the number expected under a null mutation model (*47*). We tested the de novo enrichment using twenty different missense pathogenicity predictors (see Methods for comparison section). We report genes that are identified as enriched when only counting missense DNMs with a PrimateAI-3D score ≥0.821. Some missense classifiers predicted scores in limited sets of genes, and we compensated for this by scaling the null mutation rates by the fraction of all missense sites with a pathogenicity score. For each missense predictor we estimated the excess missense DNMs without any missense classifier, and identified the pathogenicity threshold at which we captured that number of missense DNMs above that threshold. For two classifiers (DANN, LIST-S2) we found missense DNMs from healthy controls had skewed score distributions compared to the distribution of all scores genome-wide. We calculated per-threshold inflation factors via the ratio of the quantile from control DNMs to the quantile from all sites, at the same threshold. We identified an adjusted threshold as the highest threshold at which the excess corrected for the corresponding inflation factor exceeded the original excess. For example, for PrimateAI-3D the threshold was 0.821. We adjusted the genome-wide expectation for damaging missense DNMs by the fraction of missense variants that meet the threshold (roughly one-sixth of all possible missense mutations genome-wide). Each gene required four tests, one testing protein truncating enrichment and one testing enrichment of protein-altering DNMs, and both tested for just the DDD cohort and one where we excluded individuals with a protein-altering DNM in a gene previously identified with monoallelic or hemizygous inheritance for intellectual disability (ID). The genes with known links to ID were obtained from Genomics England PanelApp ID gene panels with confidence level of 3 (*108*), or found in Gene2Phenotype DDG2P subset where confidence was "definitive" (*109*). The enrichment of protein-altering DNMs was combined by Fisher's method with a test of the clustering of missense DNMs within the coding sequence. The *P* value for each gene was taken from the minimum of the four tests, and genome-wide significance was determined as $P < 6.41 \times 10^{-7}$ ($\alpha = 0.05$; 19,500 genes with four tests). We also excluded two genes, *BMPR2* and *RYR1* as borderline significant genes that already had well-annotated non-neurological phenotypes.

Fig. S22 shows the number of candidate genes discovered for each classifier, and while PrimateAI-3D is among the best performing algorithms. Because the number of candidate disease genes discovered in DDD (< 300) is a very small number compared to the number of variants evaluated in the six clinical benchmarks (which typically contain on the order of tens of thousands of variants), it is too noisy to be used as a meaningful metric for evaluating classifier performance, as the differences between the algorithms are not statistically significant.

### Supplementary Text

#### 1 Protein voxelization

Protein voxelization is the process of converting sets of protein atomic coordinates into tensors that have the same shape for all sets of coordinates. We explain our voxelization method first using two-dimensional atomic coordinates, followed by a straightforward extension to three dimensions.

To voxelize a variant, we first obtained the protein structure in which the variant occurs and determine the atomic coordinates of the Cα atom of the variant (Fig. S11A: blue point). Note that the protein structure either has the reference amino acid or no amino acid at all at the site of mutation (fill-in-the-blank in 3D; section "Model training"), but never an alternative amino acid. Next, we reduced the atoms of the protein to Cα atoms of alanine residues (Fig. S11A: red points). Then we defined a square of a certain size (Fig. S11B; imagine e.g. 6x6 Angstrom (Å)). We subdivided this square into smaller regular-sized squares called “voxels”, in this case 3x3 voxels of size 2Åx2Å. We identified each voxel by its row and column index. For example, the voxel in the top right is referred to as $V_{(1,1)}$. Then we moved the center of the central voxel in the voxel grid (Fig. S11: $V_{(2,2)}$, in the middle of the grid) onto the Cα coordinate of the residue with the mutation so that both points have the same coordinates (Fig. S11C: blue point from Fig. S11A has the same coordinates as the center of voxel $V_{(2,2)}$ in Fig. S11B). Now we looped through all voxel centers in the grid, starting with $V_{(1,1)}$. We determined the Cα atom that is closest to the center of $V_{(1,1)}$ (i.e. the “nearest neighbour”; Fig. S11C: point *NN*) and still within the scan radius $r$ (Fig. S11C: red circle; typically 5Å). We calculated the Euclidean distance $d$ between the two points (Fig. S11C). The closeness $c$ between $V_{(1,1)}$ and *NN* is then defined as

**Equation 1**

$$c\left(V_{(1,1)}, A, C\alpha\right) = 1 - d/r$$

Finally, we repeated this procedure for all 21 amino acids (20 standard amino acids and one additional amino acid representing all non-standard amino acids), starting with the reduction to the Cα atoms of all cysteine residues, instead of alanine. This is followed by a repeat of the previous 21 iterations, but with Cβ atoms, instead of Cα (however, the Cα is kept for the variant with the mutation, i.e. the coordinates of the voxel grid center stay the same; Fig S11D). In the end, each voxel is associated with 42 closeness values, one for each amino acid and atom type combination (Figs. S11E-F). We refer to this set of values as the atomic distance profile of a variant. We also experimented with including more atom types, but only found diminishing returns.

The atomic distance profile is the first of two outputs of the voxelization procedure. The second output is calculated with the following differences: we did not reduce to certain amino acids or atom types. The *NN* therefore becomes simply the closest atom to a voxel center, across all amino acid atoms in the protein. Then we looked up the position of the residue from which atom *NN* comes in the sequence of the target protein. Instead of calculating closeness $c$, we only defined a function $s$ that maps each voxel to the sequence position of this closest residue. For example, given a target variant at sequence position 11 in protein *BRCA1*, the atom closest to $V_{(1,1)}$ may come from a glycine residue at sequence position 32. Hence, $s(V_{(1,1)}) = 32$.

Moving the algorithm above from two to three dimensions, we only needed to increase the dimensionality of the voxel grid by one dimension. For example, the first voxel $V_{(1,1)}$ becomes $V_{(1,1,1)}$. Given a 3x3x3 voxel grid, the central voxel $V_{(2,2)}$ becomes $V_{(2,2,2)}$.

Evaluating different voxel grid sizes (from 3x3x3 to 19x19x19) and voxel sizes (from 1Åx1ÅxÅ to 3Åx3Åx3Å), we found that grid sizes above 7x7x7 and voxel sizes below 2Åx2Åx2Å failed improve performance. Therefore, we chose a grid size 7x7x7 with 2Åx2Åx2Å voxels. Note that an atom does not actually need to be inside the voxel grid to contribute to both voxelization outputs $c$ and $s$. It suffices if the atom is within the scan radius $r$ of a voxel. Conversely, even if an atom is within a scan radius, it does not automatically imply that the atom will contribute to $c$ and $s$. With our chosen parameters, on average the atoms of 41 (minimum 1, maximum 205) different residues are within the scan radius of a voxel in the grid. Of those, an average of 21 different residues (minimum 1, maximum 67) are mapped to by $s$.

#### 2 Details of structure quality features

We included several features indicating the confidence in the structure surrounding each voxel. First, we defined four binary indicators: whether the protein structure is present in AlphaFold DB as-is (i.e., with a perfect sequence match to one of our protein sequences), whether we used Modeller with AlphaFold DB structures as templates (in case the match was not perfect), whether we used HHpred and AlphaFold DB structures as templates in Modeller (when large parts of a protein were missing in AlphaFold DB), or whether we used HHpred exclusively (in case there was no similar match in the AlphaFold DB). For the structures that involved HHpred, we added three features corresponding to the minimum, mean and maximum predicted TMscore (*168*) of the templates found by HHpred. We also included B-factors from both AlphaFold DB and Modeller (where available) using the mapping function, which maps each voxel center to the sequence position of the residue with the closest atom to that center. For example, if $s\left(V_{(1,1,1)}\right) = 31$, then we looked up the AlphaFold DB B-factor of residue 31 and added it to the feature vector of voxel $V_{(1,1,1)}$. HHpred output contains a sequence alignment of up to 6 template proteins with known structure. For each residue in the target protein, we counted how often the other sequences in the alignment have non-gap amino acids at that position. This value indicates how many templates could be used at each position when a structure needs to be predicted with HHpred. We added this value to the feature vector of each voxel in the same way as for B-factors. In total, there are 10 features describing the quality of the protein structure around a voxel (Fig. S12B).

**3 Species-differentiable evolutionary profiles**

Let $\Sigma = \{\mathrm{A}, \dots, \mathrm{W}, -\}$ be the set of all 20 amino acids A, … , W plus a gap token denoted as "$-$". $M$ is a $|S|$-way multiple sequence alignment $S$ ($|S| = 592$) and $M_i^s$ is the amino acid at protein position $i$ in alignment $s \in S$. The evolutionary profile of amino acid $A$ at protein position $i$ is defined as $f_i^A = \sum_s w[M_i^s = A]$, where [] is the Iverson bracket and $w = \frac{1}{|S|}$ is a normalization constant. $f$ calculated only from the multiz100 alignments ($|S| = 100$) is part of the input features for PrimateAI (*17*). We interpreted $f$ as a signal for human pathogenicity. A constant $w$ implies that variants from all species and alignments are equally important contributors. For example, a variant in chimp (closely related to human) increases the frequency of an amino acid by the same amount as the same variant in zebrafish (distant from human). We argue that this should not be the case. For PrimateAI-3D, we therefore allowed each of the 592 whole proteome alignments to have a different weight. More precisely, we defined a new evolutionary profile $f'^{A}_{i} = \sum_s w_s[M_i^s = A]$, where $w_s$ is the weight of each alignment.

For example, assume that $i = 31$, $S = \{S_1, S_2, S_3\}$ and $\Sigma = \{L, P\}$ and that the two amino acids in the alignments at position 31 are $L$ for alignment $S_1$ and $P$ for alignments $S_2$ and $S_3$ respectively. Assume further that $w_{S_1} = 0.1$, $w_{S_2} = 0.05$ and $w_{S_3} = 0.2$. The one-hot encodings of the amino acids at position 31 are $< 1{,}0 >$ (for $S_1$), $< 0{,}1 >$ (for $S_2$) and $< 0{,}1 >$ (for $S_3$). It follows that $f'^{L}_{31} = w_{S_1} = 0.1$ and $f'^{P}_{31} = w_{S_2} + w_{S_3} = 0.05 + 0.2 = 0.25$.

Instead of calculating each $w_s$ from the alignments themselves (e.g. via similarity to other sequences, as often performed for Potts models of MSAs (*169*)), we initialized each $w_s$ to $\frac{1}{|S|}$ before training, and let the parameters be differentiable during training. This means PrimateAI-3D learns itself how important each of the 592 species is as contributors to the pathogenicity signal of $f'^{A}_{i}$.

**4 Evolutionary profile features**

The evolutionary profile function $f'_i$ is equivalent to a 1D-convolutional layer without bias if we interpret the 21 amino acids (20 standard amino acids plus a gap token) as input samples and the number of alignments ($|S|$) as features (considering only one amino acid, the 1-hot encoding of an amino acid is only a single value that is either 0 or 1). Denote this convolutional layer $Conv_{f'}$. To make use of the typical parameters associated with a convolutional layer, we introduced a bias term, increased the number of filters to 10 and activated each output via the ReLU activation function (the latter enables modelling non-linear relationships between species). In the end, the input to $Conv_{f'}$ are the $|S|$ 1-hot encoded amino acids at a sequence position $i$ and the output is an evolutionary profile with 10*21=210 elements. In order to merge this evolutionary profile with voxels (above), we again make use of function $s$. For example, given $s\left(V_{(1,1,1)}\right) = 31$, we extracted the $|S|$ amino acids at sequence position 31 from the alignments of the target protein and 1-hot encode and convolved them using $Conv_{f'}$. The output is concatenated with the feature vector of $V_{(1,1,1)}$, extending it by 210 elements (Fig. S12C). Note that unlike for the other features, the gradient does not stop at these 210 elements, but is backpropagated and used to update the weight $w_s$ of each alignment.

#### Supplemental Figures

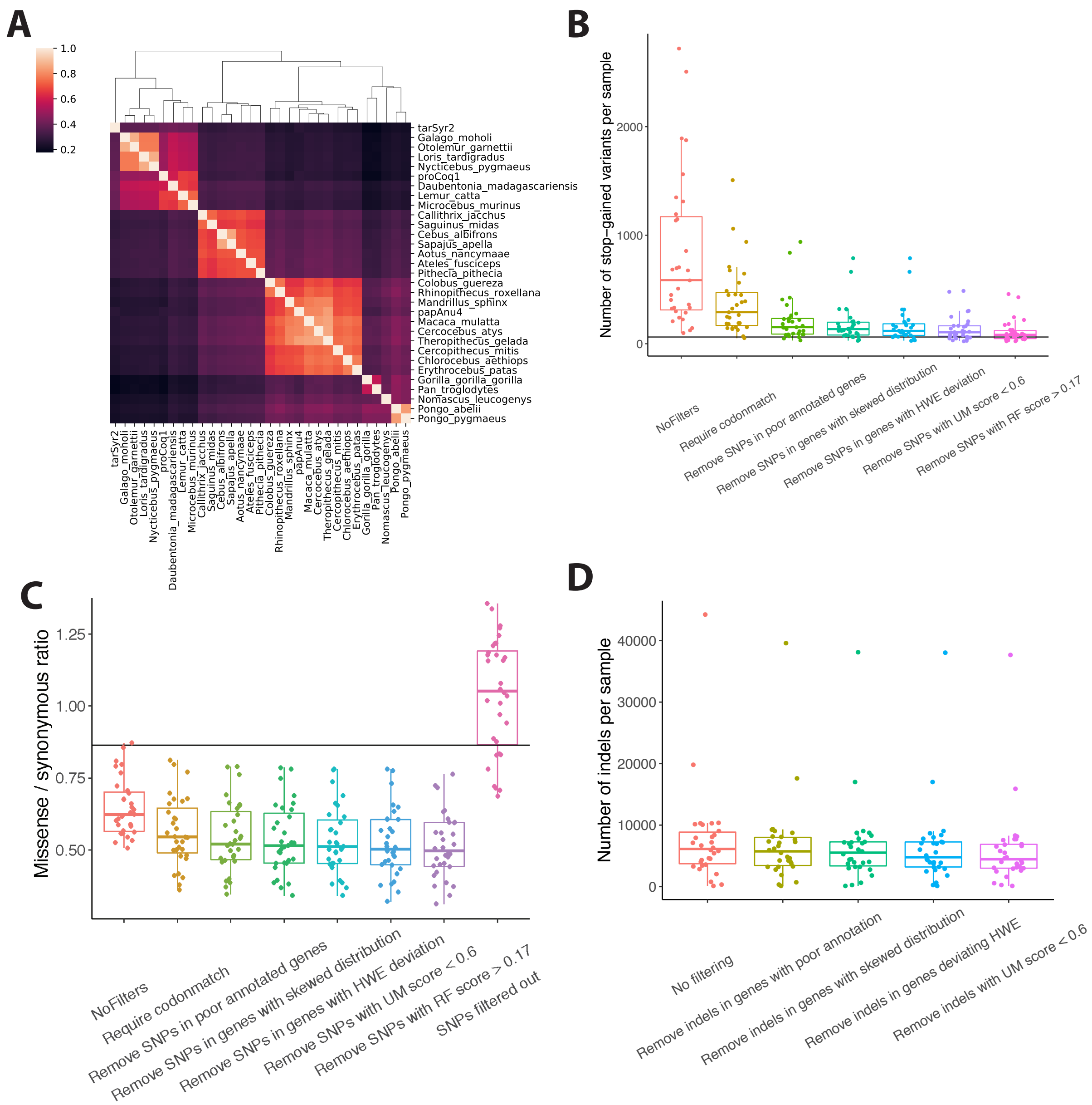


**Fig. S1. Variant filtering steps improve variant quality.** (**A**) Heatmap of Spearman correlation of codon-match metrics among 31 primate reference species naturally clustering primates into four major groups, including great apes, Old World monkeys, New World monkeys, and lemurs / tarsiers. Codon-match indicates a specific codon of primate reference species matches human. Lighter colors represent higher correlation between two species. (**B**) Boxplots showing that the average number of stop-gained variants per sample of each primate reference species was gradually reduced to close to human level after a series of variant filtering steps, including requiring codon-match, removing SNPs in poorly-annotated genes or in genes with skewed random forest score distribution or deviating from Hardy Weinberg equilibrium (HWE), and removing SNPs with unique-mapper (UM) score <0.6 or RF score >0.17. Each dot represents the average number of stop-gained variants of each primate reference species. The black line shows the average number of stop-gained variants of human samples from Platinum genome project. (**C**) Boxplots showing that missense / synonymous ratios (MSR) decreased after variant filtering steps. The pink box shows the MSR of variants that were filtered out. Each dot represents the MSR of each primate reference species. The black line represents MSR of human samples. (**D**) The average number of indels per sample of each primate reference species diminished after filtering steps. For all the boxplots, box lengths represent the interquartile of data points and the whiskers extend to 1.5 times the interquartile range from the box.

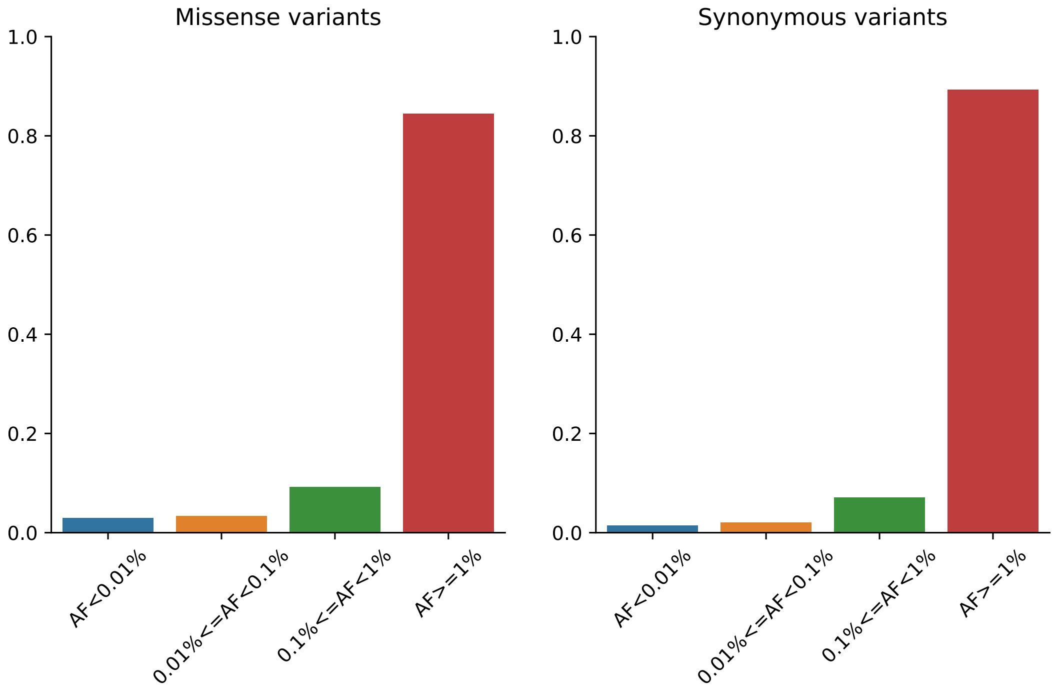


**Fig. S2. Allele frequency spectra for simulated primate variants.** Barplots show the fraction of primate missense (left panel) and synonymous (right panel) variants falling in each of the four allele frequency bins. The simulated primate variants were sampled according to the gnomAD allele frequencies, mimicking the sample sizes of primate species.

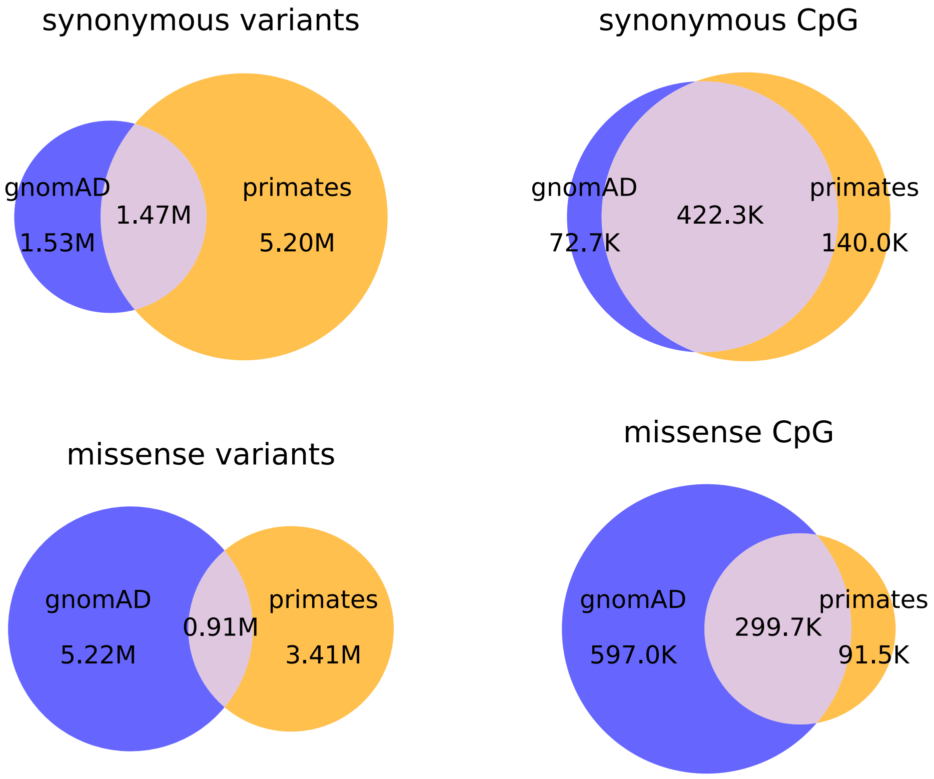


**Fig. S3. Venn diagrams show small overlap between gnomAD (blue) and primates (orange) for both synonymous and missense variants.** Large fractions of the transition variants occurring at CpG sites are shared between gnomAD and primates, particularly the synonymous variants.

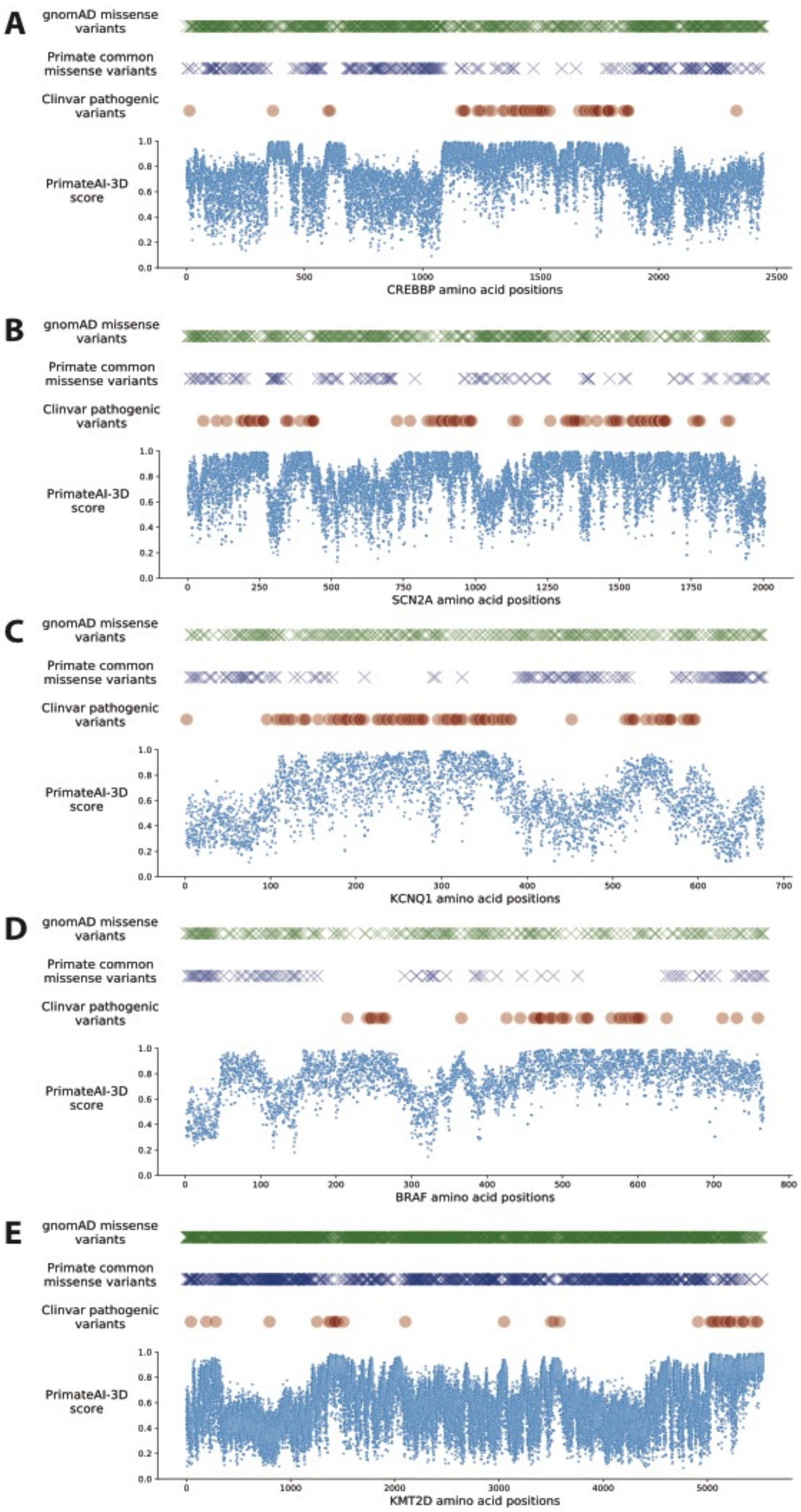


**Fig. S4. Observed gnomAD (green) or primate (purple) missense variants at each amino acid position in genes. (A-D)** The distribution of gnomAD missense variants (green crosses) along the genes *CREBBP* **(A)**, *SCN2A* **(B)**, *KCNQ1* **(C)**, *BRAF* **(D)**, and *KMT2D* **(E).** Blue crosses represent observed primate missense variants along the genes. Dark red circles represent observed ClinVar pathogenic missense variants along the genes. Blue dots of the bottom scatterplots show the predicted pathogenicity scores of all possible missense substitutions, which are the PrimateAI-3D scores at each amino acid position.

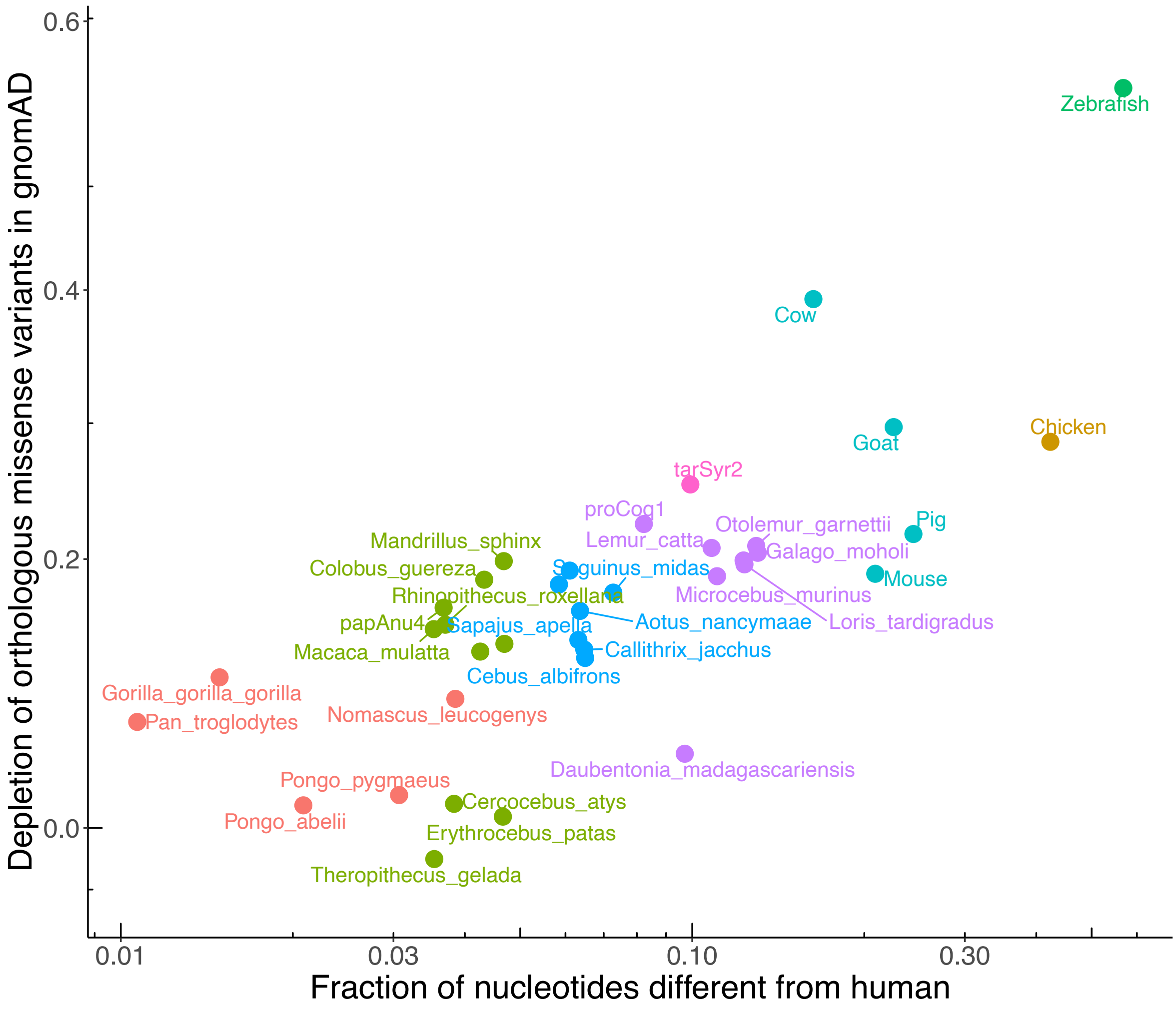


**Fig. S5. Scatter plot showing that natural selection purifies potentially deleterious missense variants across species.** The x-axis shows the depletion of orthologous missense variants observed in primates, mammals, chicken and zebrafish at common human allele frequencies (>0.1%) from gnomAD. The y-axis shows the species' genetic distance from human, measured by the fraction of nucleotides different from human.

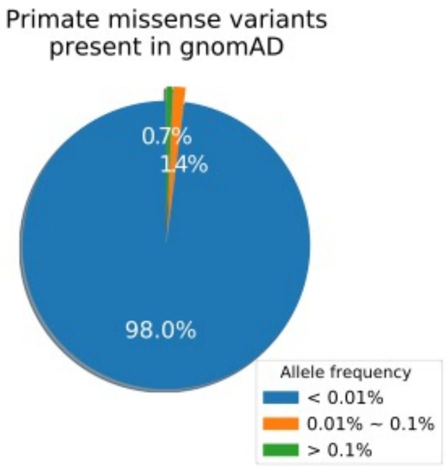


**Fig. S6. Pie chart showing fractions of primate missense variants observed at different allele frequencies (AF) in gnomAD databases.** Blue represents variants either not observed in gnomAD or with rare allele frequencies (< 0.01%). Orange and green show the fraction of primate variants with 0.01% <AF < 0.1% or common allele frequencies (>0.1%), respectively.

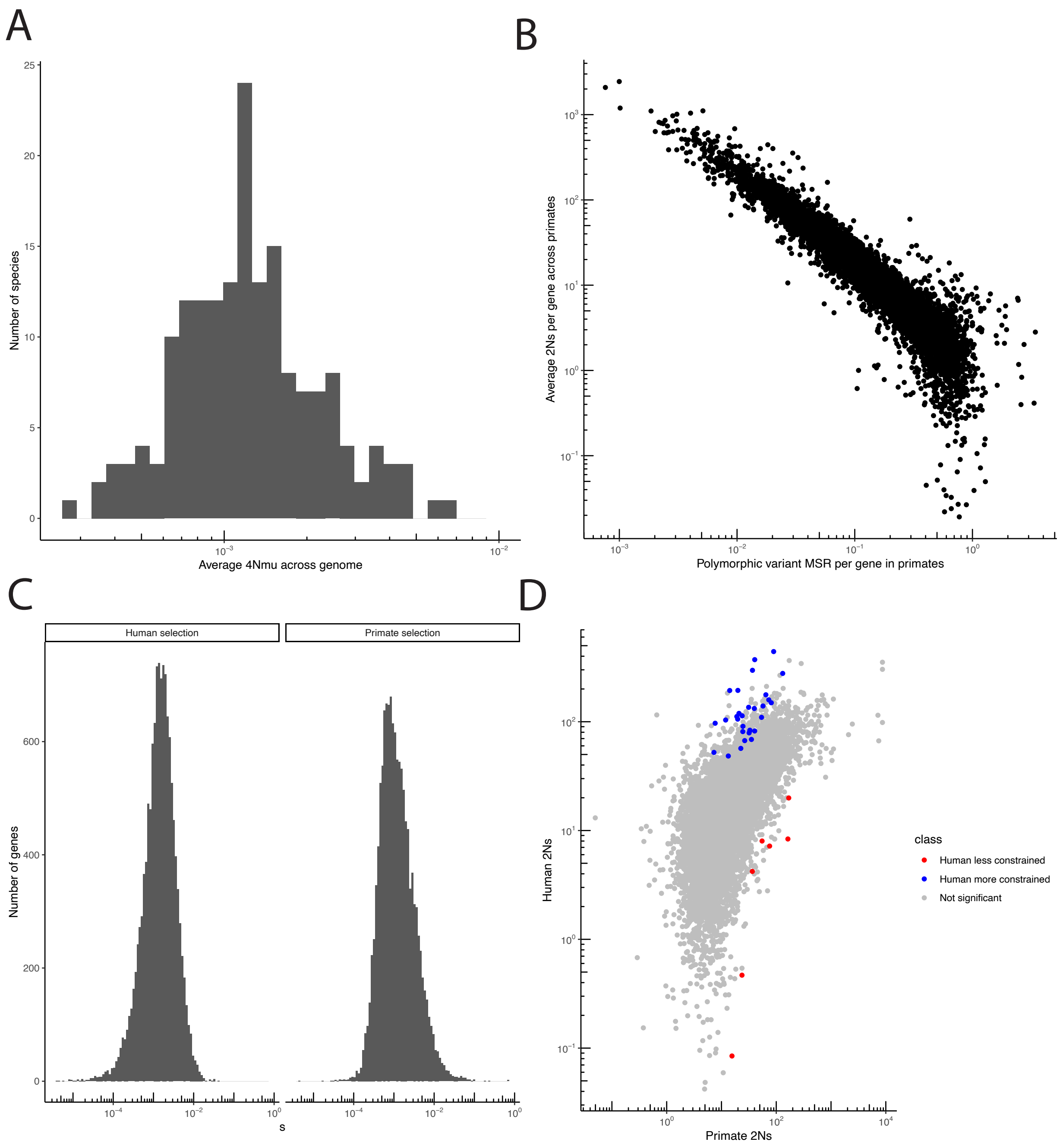


**Fig. S7. Population genetic model fitting to human and primate data. (A)** The distribution of population-scaled mutation rates across all species. **(B)** The correlation between pooled primate missense: synonymous ratio and inferred selection. The x-axis shows the missense : synonymous ratio for each gene when pooled across all non-human primates. The y-axis shows the inferred 2Ns for each gene. **(C)** The distribution of fitness effects across genes in humans and non-human primates. Histogram is over all genes that pass the filters to be used in the analysis. **(D)** The correlation between human selection and primate selection. The x-axis shows the strength of selection in humans, and y-axis shows the strength of selection in primates. Highlighted points are significant according to point the population genetic model and the MSR regression.

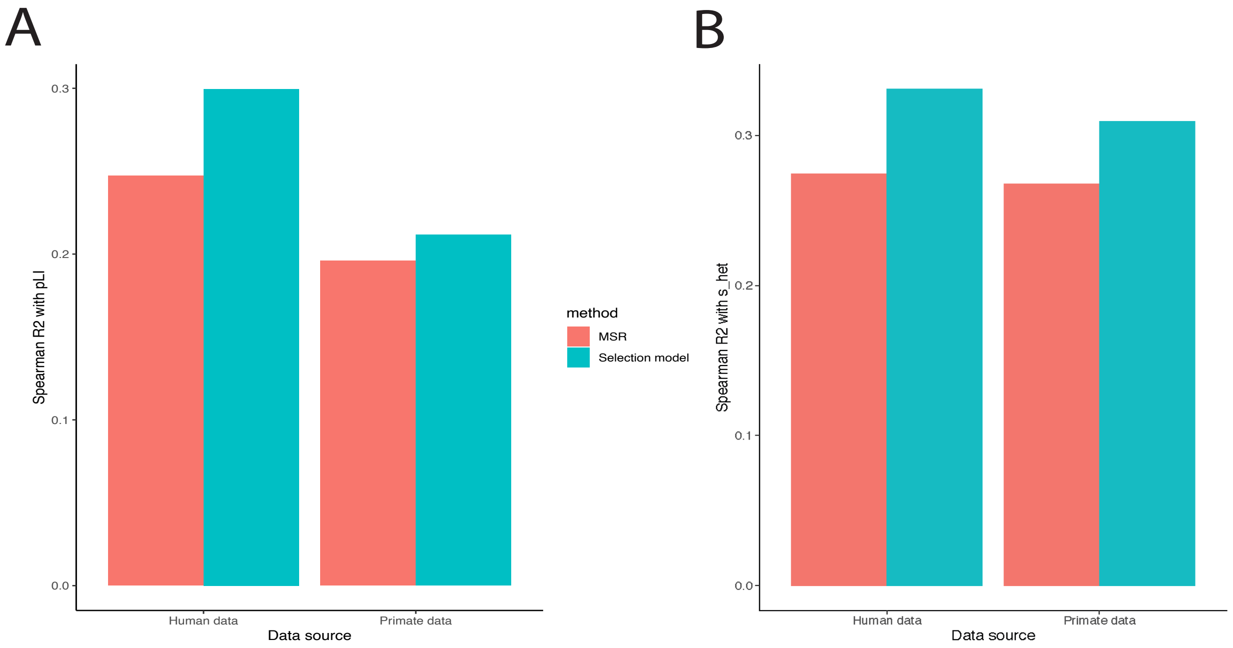


**Fig. S8. Correlation of missense : synonymous ratios and selection coefficient estimates in humans and primates. (A-B)** Barplots show the Spearman correlation of those two metrics with pLI (**A**) and s_het (**B**). Red bars represent correlation of missense : synonymous ratio of polymorphic variants with pLI or s_het. Blue bars show correlation of estimated selection coefficients. Bars are grouped by whether they are based on human data or pooled primate data.

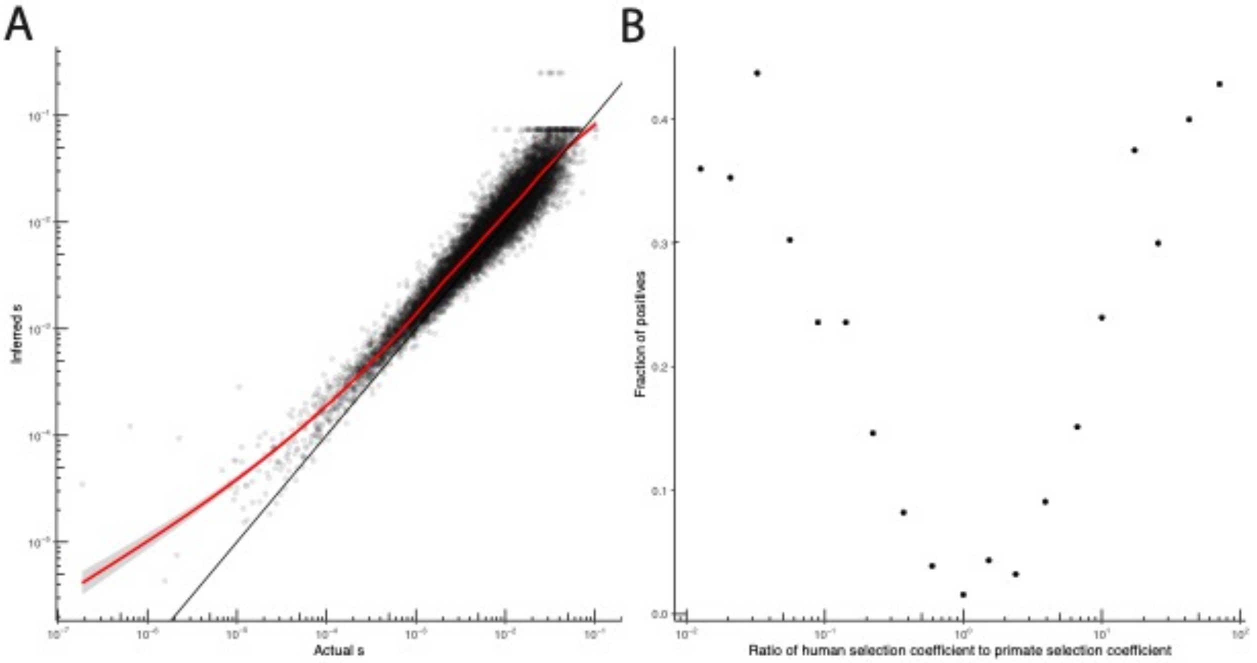


**Fig S9. Performance of Poisson random field model in simulations. (A)** Inference of simulated selection coefficients is highly accurate. The x-axis shows the selection coefficient simulated, as described in the text, while the y-axis shows the inferred selection coefficient using the method described in the text. The solid black line indicates the line y = x, while the red line is a moving average. **(B)** The likelihood ratio test is well powered and has a low false positive rate. The x-axis shows the ratio between the simulated human selection coefficient and the simulated primate selection coefficient. The y-axis shows the number of positives at an FDR of 5% following the Benjamini-Hochberg procedure. Note that the middle point, $10^0 = 1$ indicates no difference between human and primate selection.

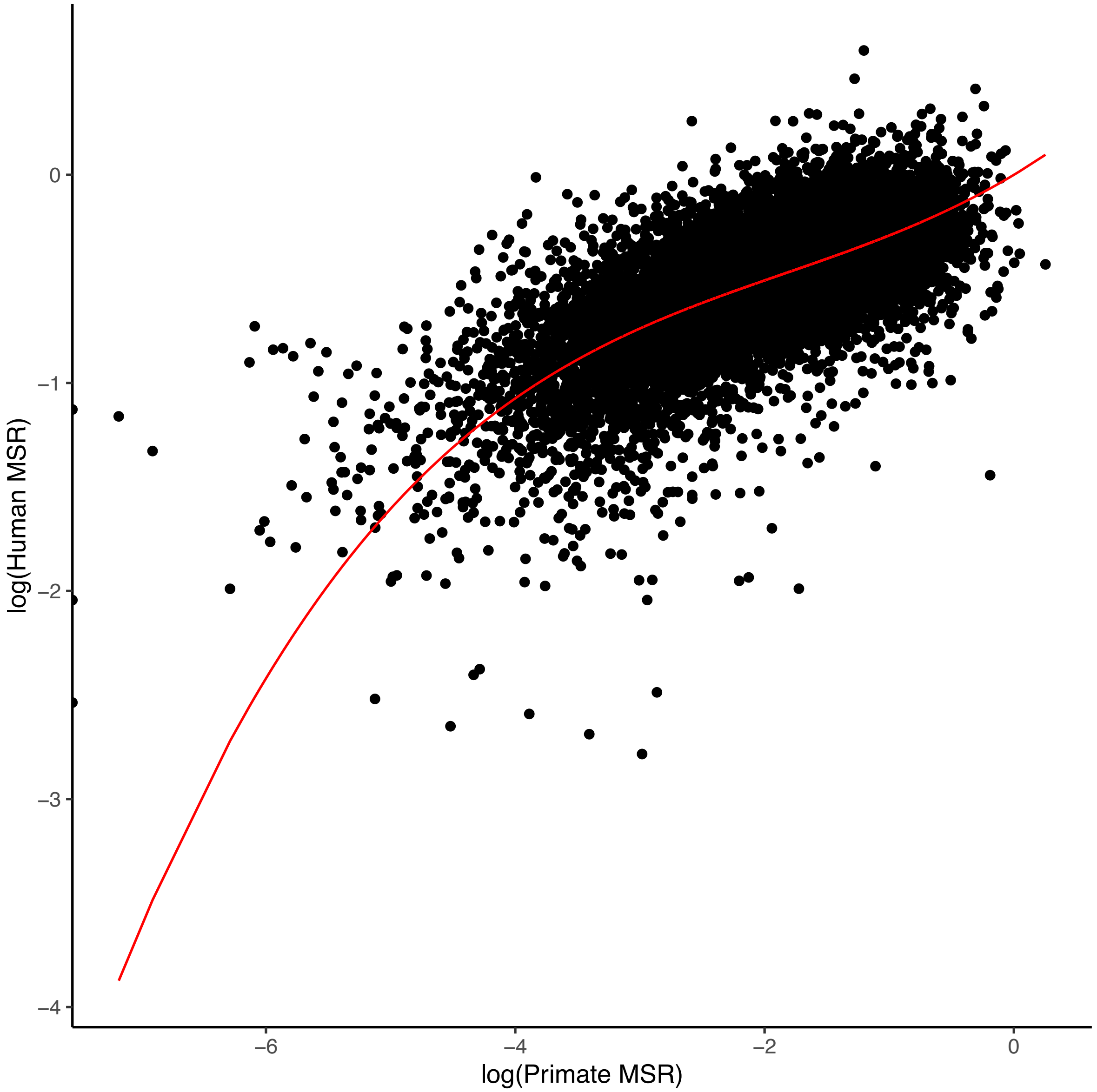


**Fig. S10: Relationship of human and primate missense : synonymous ratio based on Poisson generalized linear mixed modeling.** The x-axis shows the log-scaled missense : synonymous ratio among polymorphic variants in primates compared. The y-axis shows the log-scaled missense : synonymous ratio among polymorphic variants in humans. The red line represents the best-fit relationship as inferred by the model.

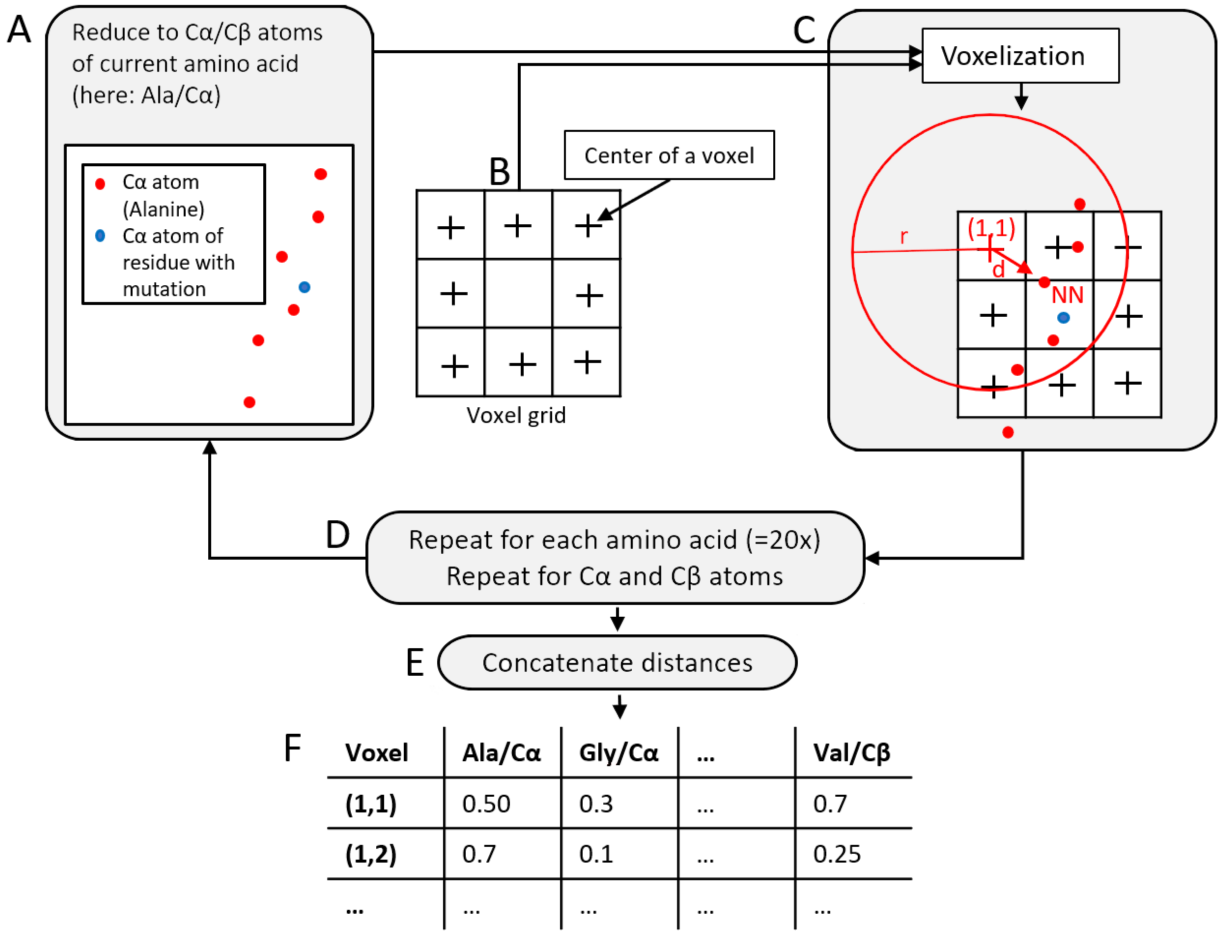


**Fig. S11. Illustration of voxelization procedure in 2D projection. (A)** Given a target site, the corresponding structure is reduced to **Cα** atoms of a particular amino acid type (here: alanine; red), plus the target **Cα** atom (blue). A voxel grid (**B**) is centred on the target **Cα** (**C**). The **Cα** atom (NN) closest to the first voxel (1,1) is determined. Their Euclidean distance $d$ is divided by maximum scan radius $r$ to obtain a relative distance. Subtracting that ratio from 1 turns the relative distance into a relative closeness. This procedure is repeated on various levels: for each voxel in the grid; for each amino acid type; for **Cβ** atoms instead of **Cα** atoms.

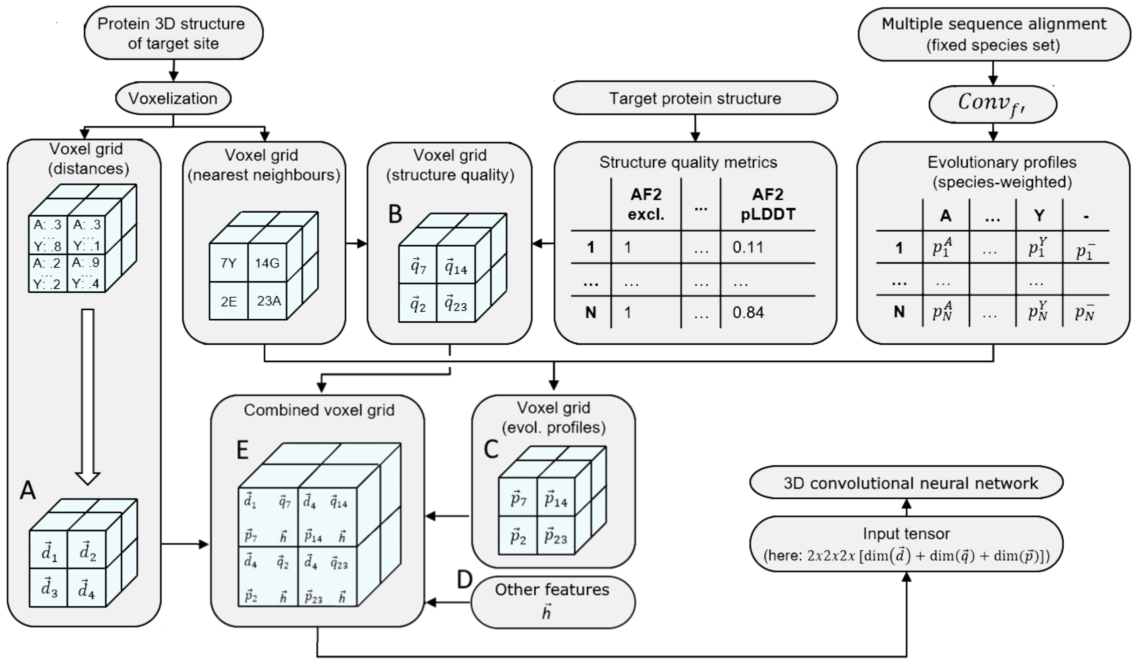


**Fig. S12. Associating voxels with features.** (**A**) Each voxel is associated with its amino acid distance profile *d*. (**B**) The mapping of each voxel center to the closest residue is used to look up structure quality metrics *q*. (**C**) Similarly, the mapping is used to associate each voxel with its evolutionary profile *p* of the closest residue. (**D**) Other features that are associated with each voxel include the 1-hot encoding of the target amino acid before mutation and indicators whether the central residue has been removed (fill-in-the-blank in 3D; section "Model training") or not. (**E**) Feature vectors *d, q, p* and *h* are concatenated and used as input to a 3D convolutional neural network.

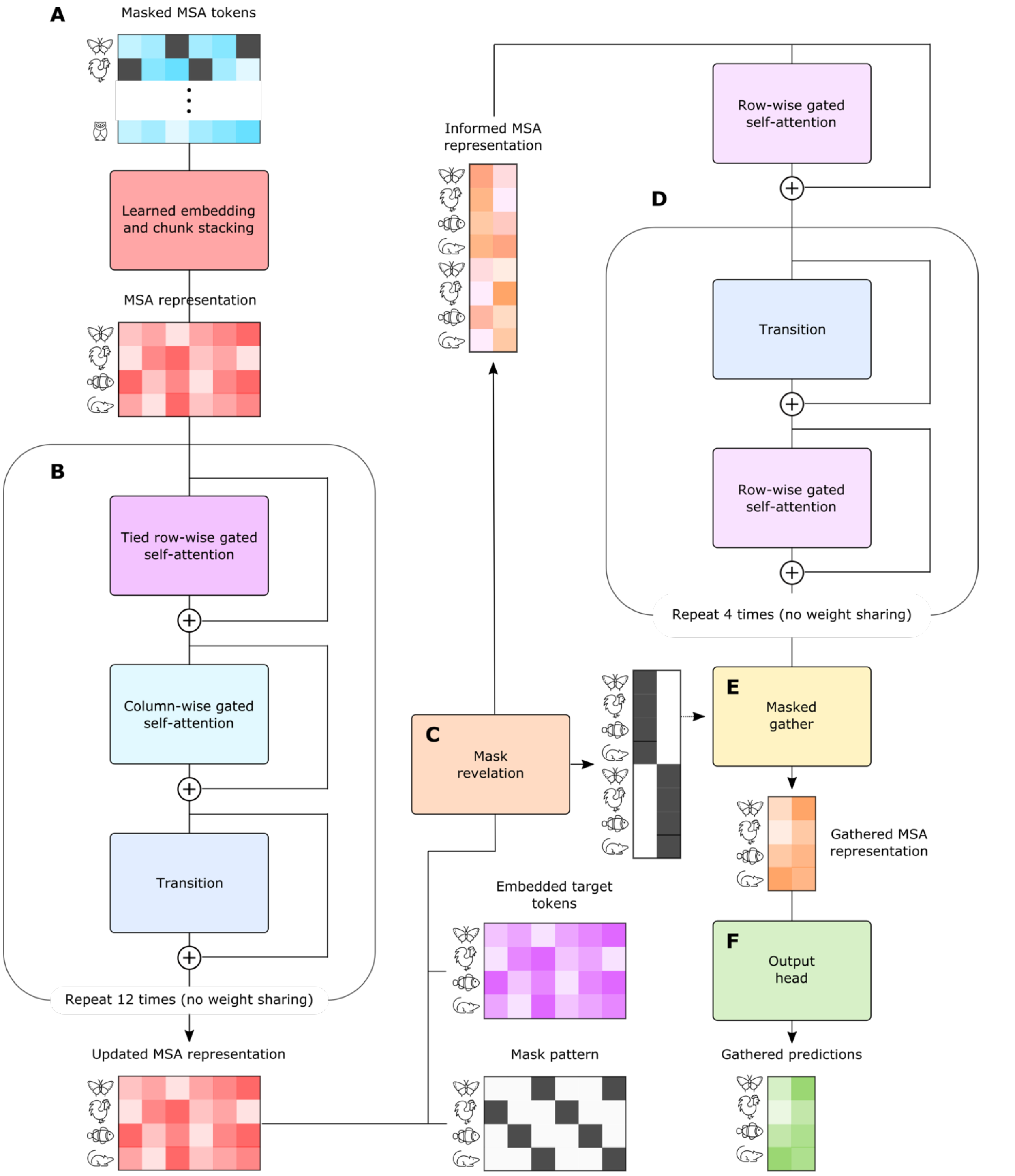


**Fig. S13: PrimateAI language model architecture.** (**A**) An initial MSA representation is created by learned embedding and stacking of MSA sequences. (**B**) Axial attention blocks develop the MSA representation. (**C**) Mask revelation gathers features aligned with mask sites. For each masked residue in a row, it reveals embedded target tokens at other masked locations in that row. (**D**) Attention is applied to gathered rows to interpret mask revelations. (**E**) MSA features are gathered from locations where target embeddings remained masked. (**F**) An output head, consisting of a transition and perceptron, maps the gathered MSA representation to predictions.

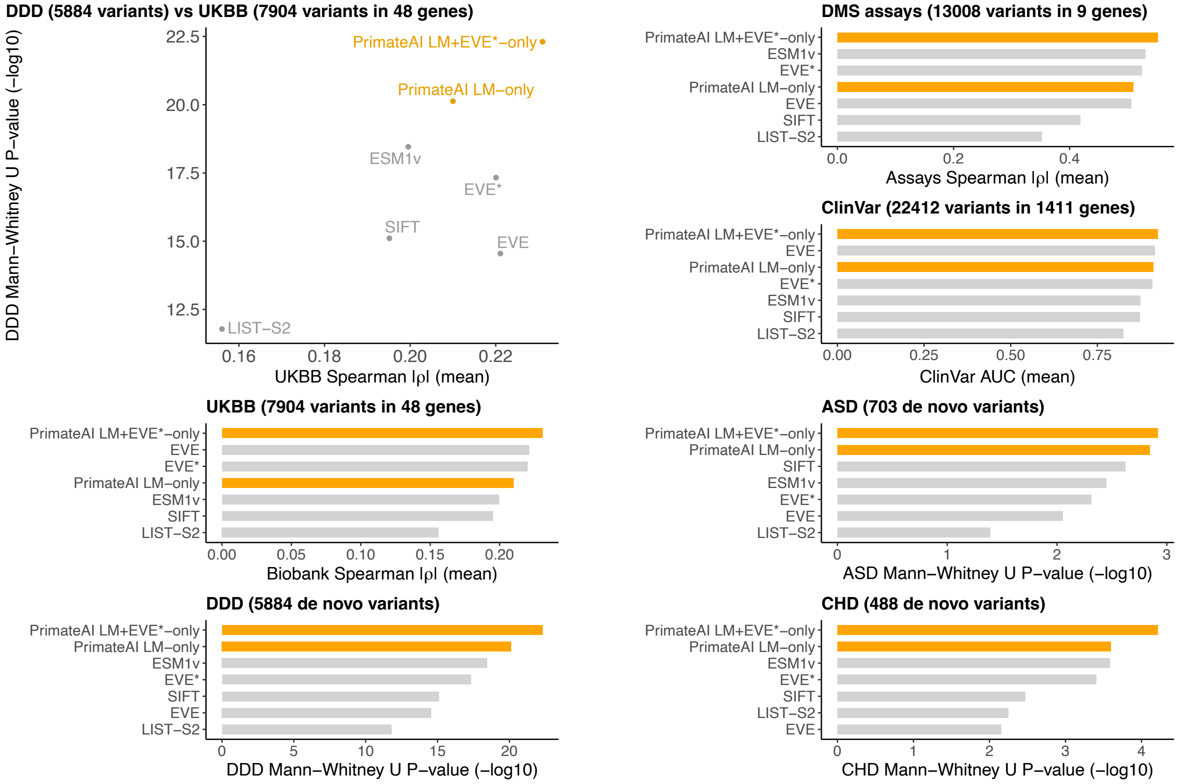


**Fig. S14. Evaluation performance of the language modelling part of PrimateAI-3D (PrimateAI LM-only)**. The performance of PrimateAI LM-only is compared to the replicated VAE part of EVE (labelled "EVE*") model and their combined score (labelled "PrimateAI LM+EVE*-only"). It is further compared to a selection of competitive unsupervised methods (ESM1v, SIFT, LIST-S2). In clockwise direction starting from the top left, the individual panels correspond to evaluation on DDD vs UKBB, DMS assays, ClinVar, ASD, CHD, DDD and UKBB. For DMS assays and UKBB, the summary statistics are given in terms of absolute value (|corr|) of correlation between score and an experimental measure of pathogenicity, i.e., mean phenotype (UKBB) or assays score (DMS assays). For DDD/ASD/CHD, we calculated the P-value of Mann-Whitney U test for control and case distributions over all datasets. For ClinVar, we measured the AUC averaged over all genes.

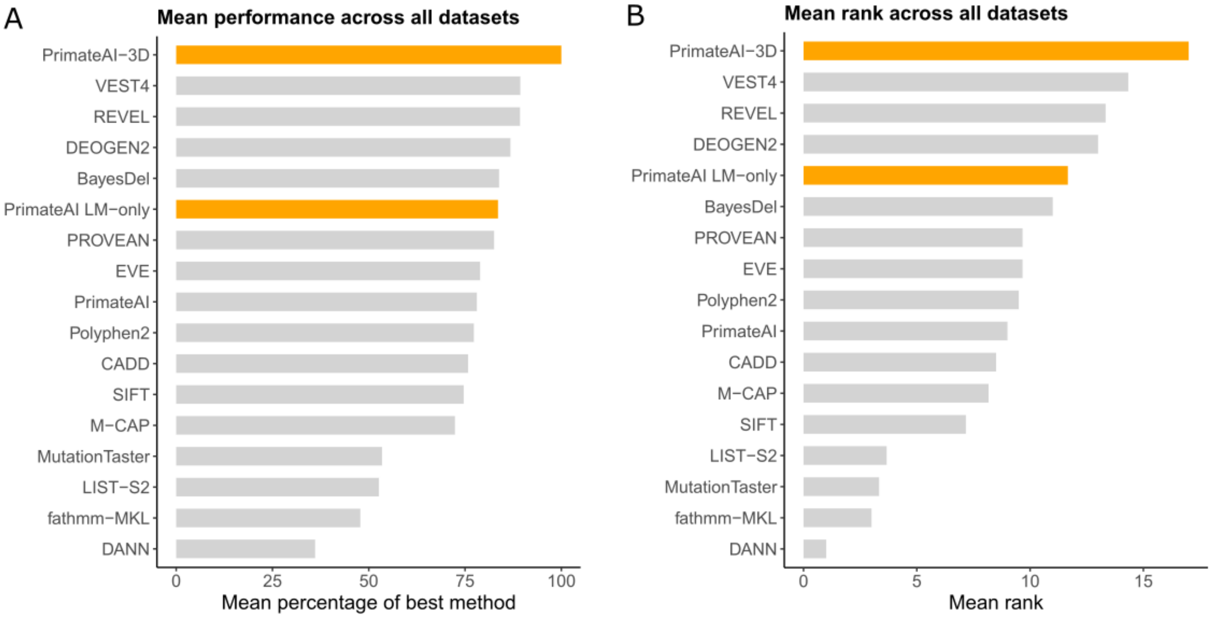


**Fig. S15. Combining performance metrics from all evaluation datasets. (A)** For each of the six evaluation datasets, the performance metric of each method was divided by the maximum performance achieved across all methods. The average of that percentage across datasets is the "Mean percentage of best method" for each method. **(B)** The rank of the performance metric of each method is determined separately for each dataset and then averaged to create one mean rank value for each method.

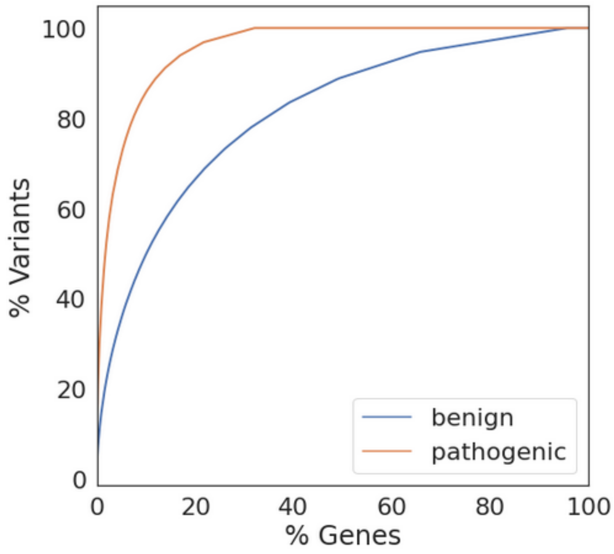


**Fig. S16. Fractions of ClinVar variants covered by genes in ClinVar.** The x-axis is the percentage of genes in ClinVar with at least one 1-star variant annotation. The y-axis is the percentage of ClinVar 1-star variants covered by genes in ClinVar. Each point in a line indicates the minimum percentage of genes required (x-axis) to cover a certain percentage of ClinVar variants (y-axis). For example, 50% of all pathogenic variants in ClinVar come from 1.8% of ClinVar genes.

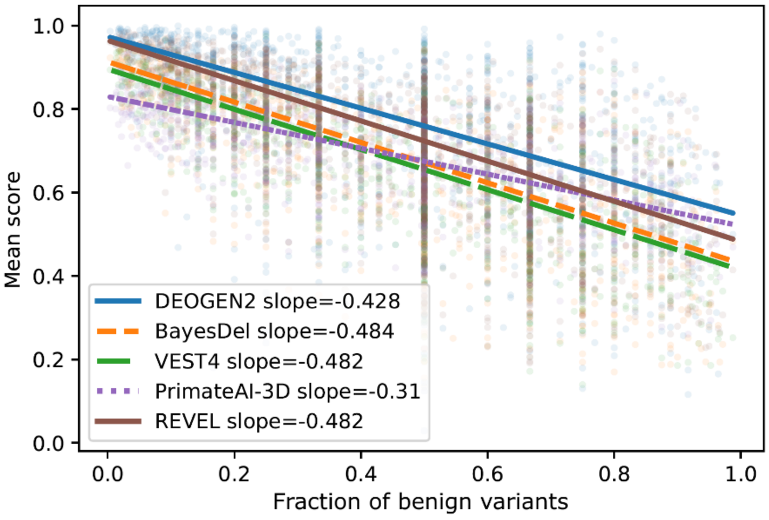


**Fig. S17. Dependency of pathogenicity scores on the per-gene ratio of benign to pathogenic variants in ClinVar.** We calculated the ratios "#benign/(#benign+#pathogenic)" for ClinVar genes with at least one benign and pathogenic variant (x-axis). For each prediction method, we then calculated the mean score for each gene (y-axis). Fitting a separate regression line for each method, the slope of PrimateAI-3D (-0.31) is below the slopes of other methods (-0.428 to -0.484), indicating that other classifiers' performance on ClinVar are affected by fitting to the per-gene ratio of benign to pathogenic variants in ClinVar, likely because they were trained directly on ClinVar or highly overlapping databases such as HGMD.

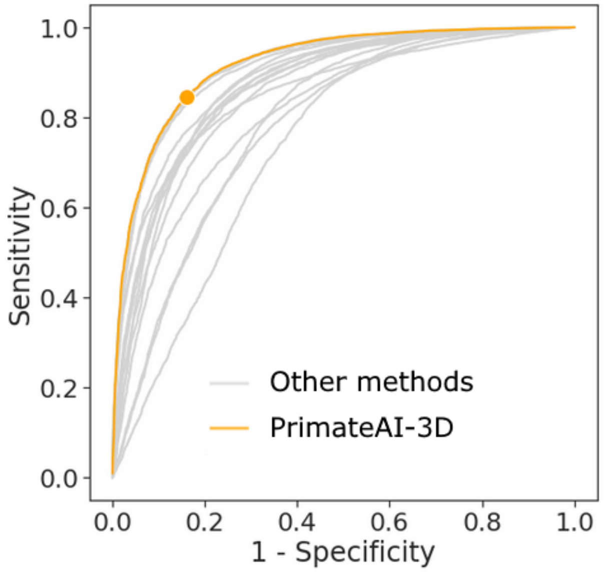


.

**Fig. S18. Global ClinVar ROC curves.** Other methods represent those classifiers used in Fig. 3. For PrimateAI-3D, the point that maximizes sensitivity+specificity is highlighted (sensitivity: 84.7%; specificity: 84.1%; percentile threshold: 57.7). This sensitivity+specificity is also the maximum among all other methods.

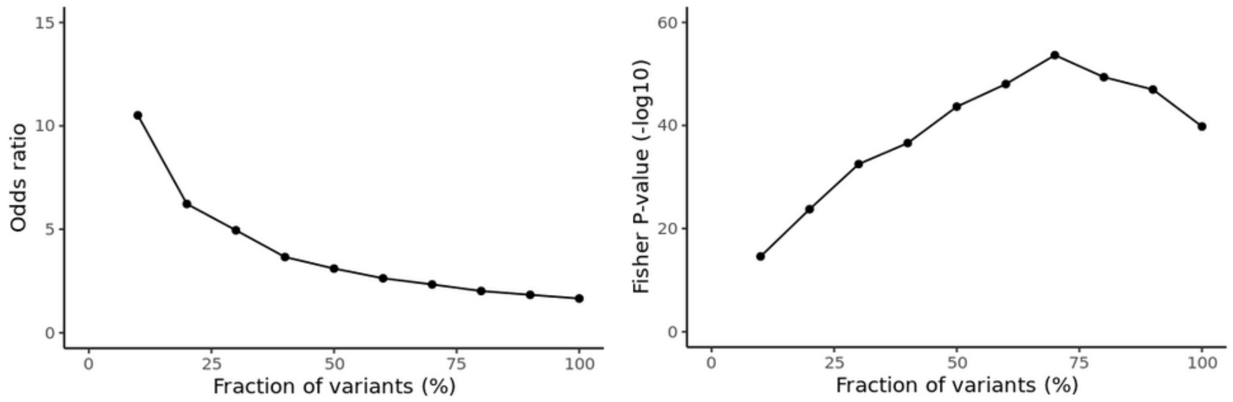


**Fig. S19. PrimateAI-3D detects low-confidence ClinVar annotations.** We took different percentages of the most confident PrimateAI-3D predictions and measured their agreement with benign and pathogenic annotated missense variants found in the September 2017 version of ClinVar. We also distinguished between ClinVar variants whose annotations were changed or unchanged between September 2017 and September 2021. We determine the odds of annotation change in variants that agree between PrimateAI-3D and ClinVar (left panel). We do the same for variants that disagree. The y-axis is the conditional maximum likelihood estimate of the ratio of the two odds. The x-axis indicates the fraction of top PrimateAI-3D variants. For example, a 20% fraction of variants means that we used the 10% most pathogenic and 10% most benign predicted variants. The x-axis of the right panel is the same as in the left panel and the y-axis indicates the Fisher's exact test P-value of the corresponding odds ratio shown in the left panel.

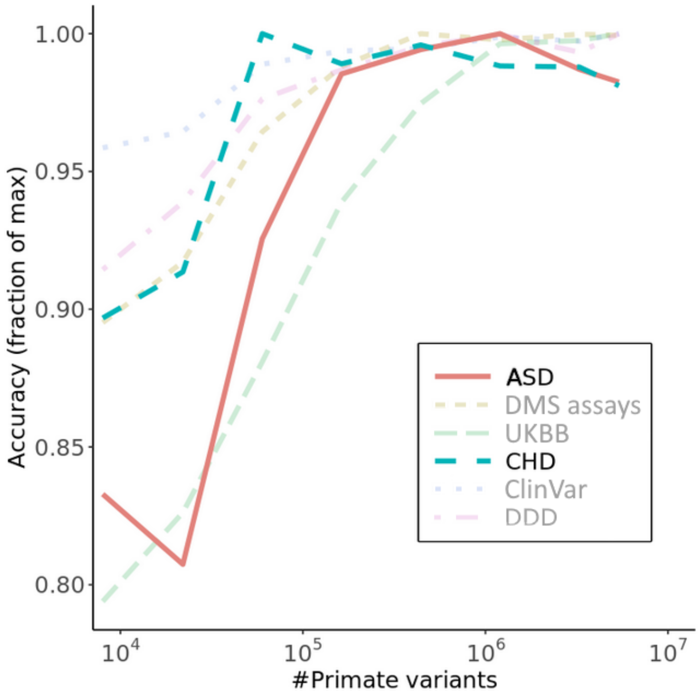


**Fig. S20. Impact of training dataset size on classification accuracy (extended).** Performance of PrimateAI-3D increases with the number of common human and primate variants in the training dataset (x-axis). Performance of each dataset (y-axis) was divided by the maximum performance observed across all training dataset sizes. ASD and CHD are highlighted because they were excluded in Fig. 5A.

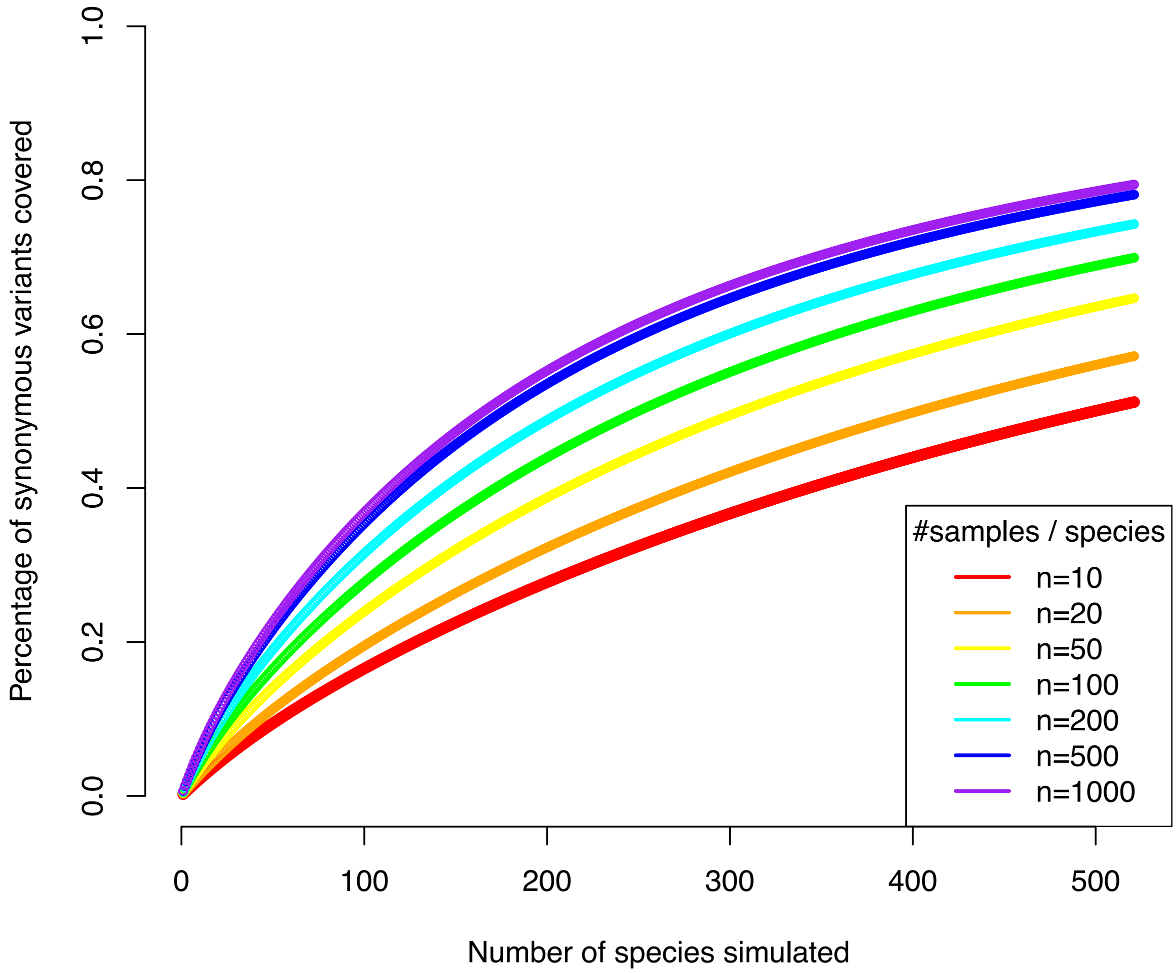


**Fig. S21. Saturation of human synonymous variants by sampling common variants present in the 521 extant primate species.** The line colors represent various sample sizes for the simulated primate species, including 10, 20, 50, 100, 200, 500 and 1000.

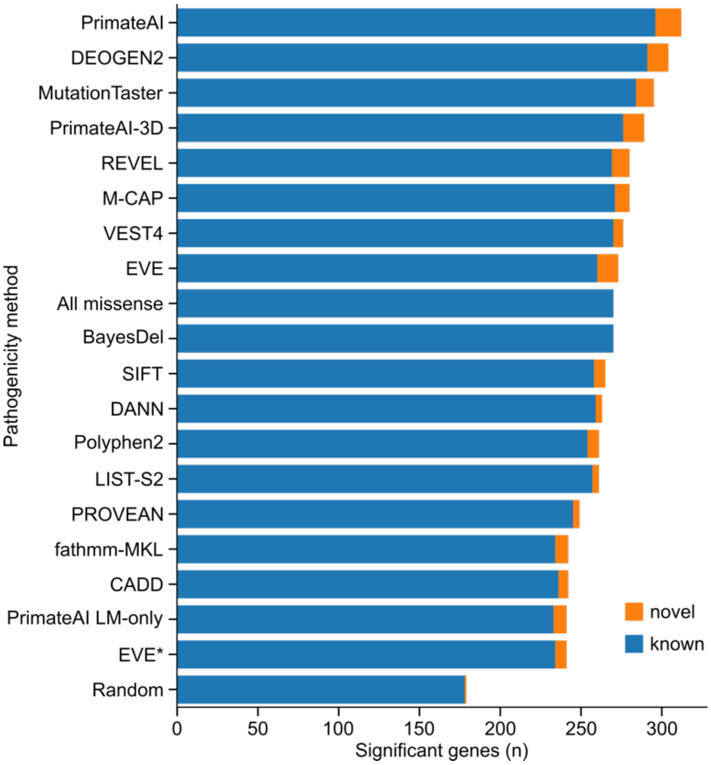


**Fig. S22. Number of genome-wide significant genes by missense pathogenicity prediction methods, identified through enrichment of de novo mutations over expectation.** Note that due to the small number of genes discovered (< 300) the differences between the algorithms are not significant.

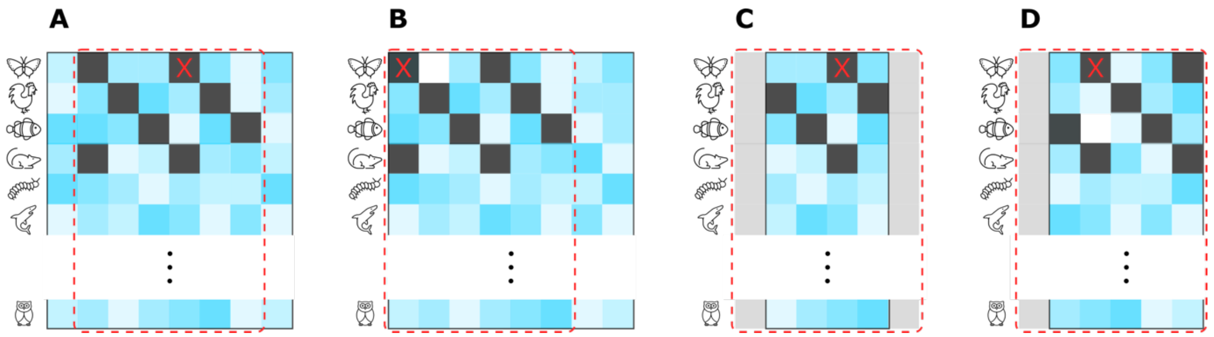


**Fig. S23: Cropping, padding, and masking of MSAs for PrimateAI LM.** A location of interest in the query sequence is indicated by an X, mask locations are black, padding is gray, and crop regions are indicated by a dashed line. In these examples, mask stride is 3 and cropping window width is 6 residues. (**A**) Away from MSA edges, a position of interest is at the right side of the center of the crop region. (**B**) A crop region is shifted to the right of the location of interest to avoid going over an MSA edge. (**C**) An MSA for a short protein is padded to fill a crop region. (**D**) A crop region is shifted to the right of a location of interest to minimize padding and the MSA is padded to fill the crop region.

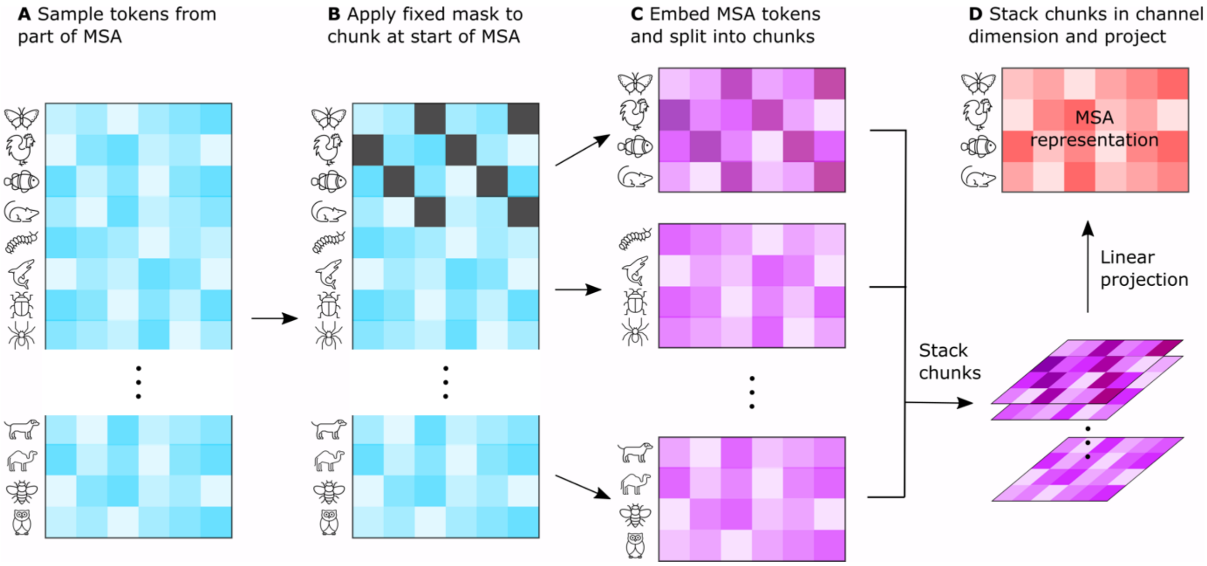


**Fig. S24. MSA masking, learned embedding, and chunk stacking.** (**A**) Part of an MSA with a contiguous set of residues and a random set of non-query protein sequences is sampled. (**B**) A fixed mask pattern is applied to a chunk of sequences at the start of the MSA. In this example, the mask pattern is applied to the first 4 sequences and has a stride of 3. (**C**) Tokens are replaced with learned embeddings, which are summed with learned position embeddings for residue columns before layer normalization. The embedded tokens are divided into chunks, which (**D**) are concatenated in the channel dimension and then linearly projected to form an initial MSA representation.

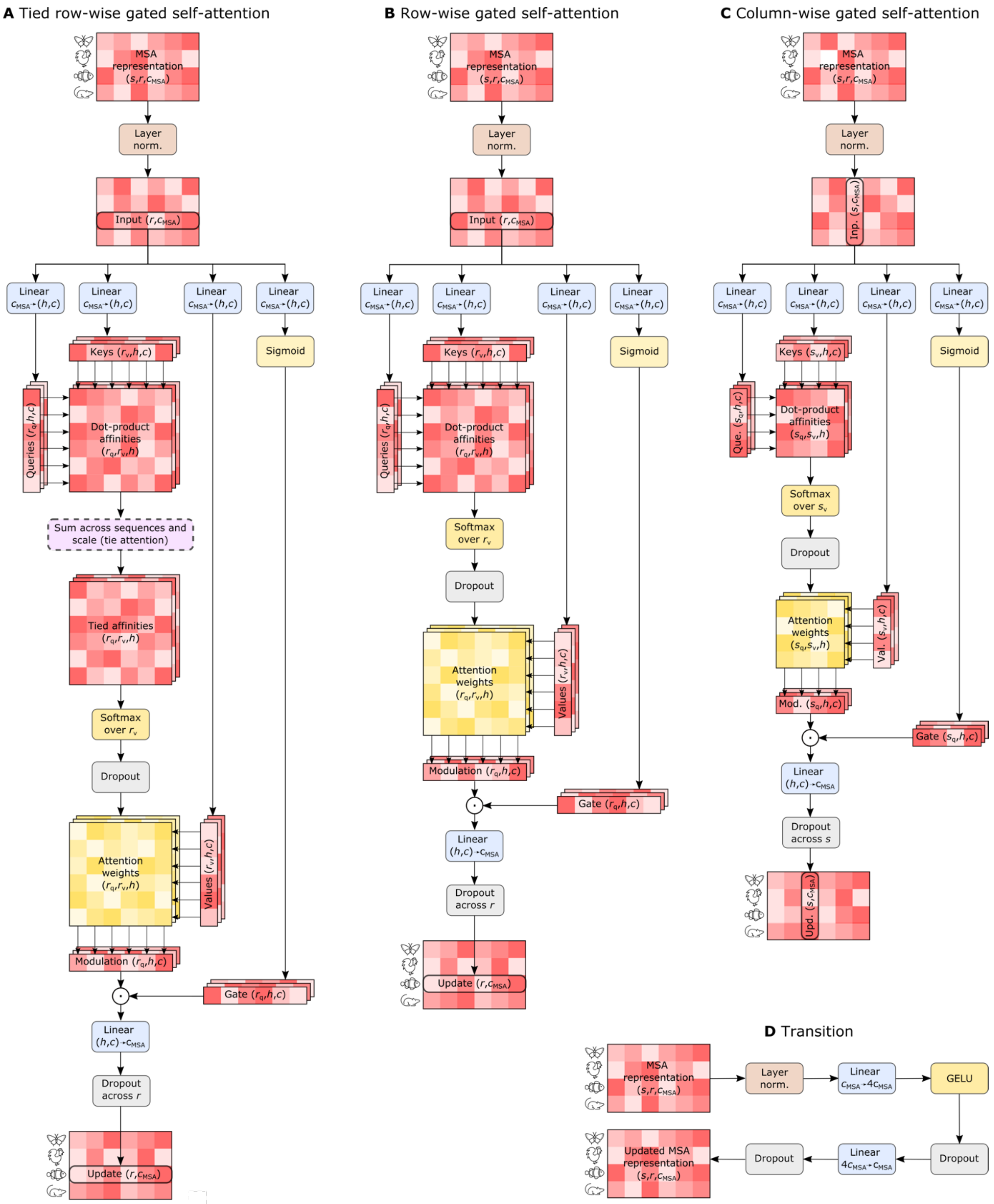


**Fig. S25. PrimateAI language model components.** (**A**) Tied row-wise gated self-attention. (**B**) Row-wise gated self-attention. (**C**) Column-wise gated self-attention. (**D**) Transition. Dimensions are shown for sequences, $s = 32$, residues, $r = 256$, attention heads, $h = 12$, and channels, $c = 64$ and $c_{\mathrm{MSA}} = 768$.

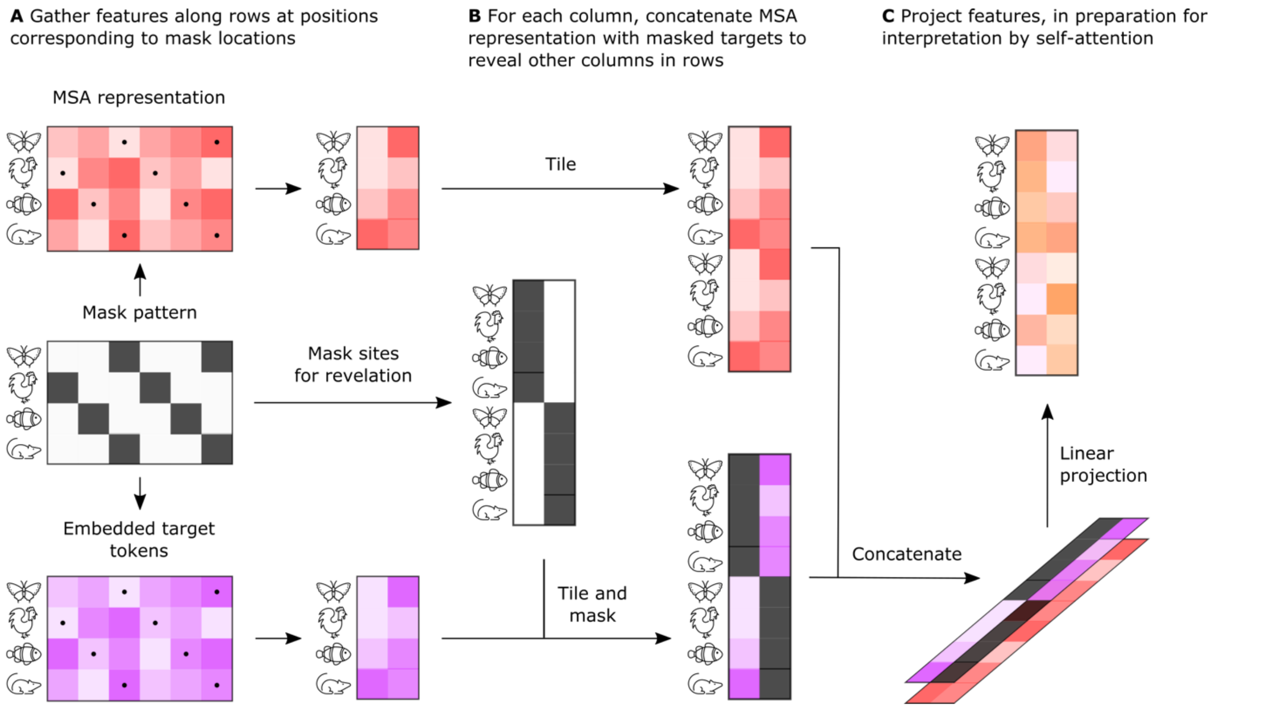


**Fig. S26. Mask revelation.** (**A**) A mask pattern is used to gather features, indicated with dots, from the updated MSA representation and embedded target tokens. (**B**) For each protein, for each masked residue in that protein, reveal embeddings for residues at other masked locations within that protein. The partially revealed target embeddings are concatenated with the MSA representation and (**C**) linearly projected, in preparation for interpretation by self-attention.

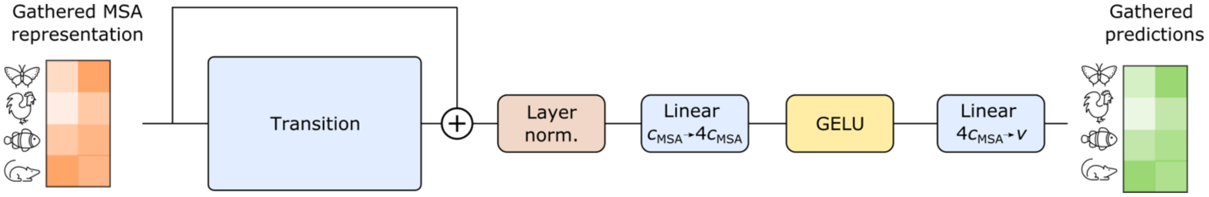


**Fig. S27. Revelation output head.** Dimensions are shown for channels, $c_{\mathrm{MSA}} = 768$, and vocabulary size, $v = 21$.

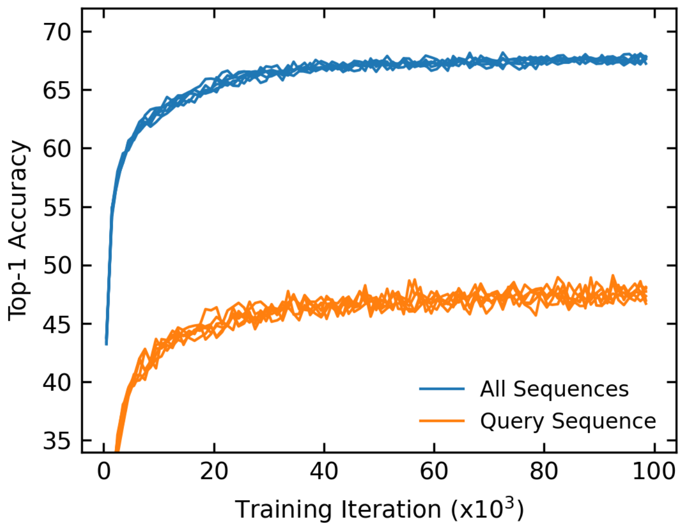


**Fig. S28. Top-1 training accuracy.** Training accuracies are averaged over $10^3$ iteration segments for six PrimateAI LM models used in our ensemble. The accuracies are lower for query sequences, which do not contain gap tokens.

#### Supplemental tables

**Table S1**. **36 pathogenic ClinVar variants present in primates curated by our clinical laboratory experts.** A variant may occur in multiple species, shown in multiple rows. Curation evidence is provided along with extra information on each variant, including in silico tool predictions, penetrance, hypomorphic or not, recessive/dominant, etc.

**Table S2**. **Expected and observed counts, inferred selection coefficients, and missense : synonymous ratio deviation for 18,883 human genes.** Expected and observed synonymous counts in humans are in columns exp_syn_h and obs_syn_h, respectively, and expected and observed missense counts in humans are exp_mis_h and obs_mis_h, respectively. Expected and observed synonymous counts in the primate cohort are exp_syn_p and obs_syn_p, respectively, and expected and observed missense counts in the primate cohort are obs_mis_p and exp_mis_p, respectively. Selection coefficients in human and primate are indicated by s_human and s_primate, respectively. Benjamini-Hochberg corrected p-values testing the null hypothesis that s_human = s_primate are provided under p.adj.popgen. MSR regression was only performed on 12,738 genes passing cut offs. Genes that did not pass the cutoff have a low_quality flag set to TRUE. For genes in the MSR regression, the log-scaled deviation from expected MSR is given under MSR_deviation. Benjamini-Hochberg corrected p-values testing the null hypothesis MSR_deviation = 0 are provided under p.adj.MSR.

**Table S3**. **Performance of PrimateAI-3D and 16 other pathogenicity classifiers on six benchmarking datasets.** Information about the six evaluation datasets (UKBB, DDD, ASD, CHD, ClinVar and DMS assays) is provided, including the number of patients and the number of variants, as well as evaluation methods. Performance metrics of PrimateAI-3D and 16 pathogenicity classifiers are provided in the remaining columns. The average rank of performance across datasets of each method is provided in the last row, with 1 corresponding to the best overall performing across all datasets and a rank of 17 corresponding to the worst.

**Table S4. PrimateAI-3D performance on 42 genes with phenotype associations from UKBB.** Column “Phenotype” provides 41 phenotypes and column “EnsembleID” provides 42 genes. For each of 78 gene-phenotype pairs, the number of common variants and evaluation metric of PrimateAI-3D (absolute value of Spearman correlation) are provided.

**Table S5**. **Enrichment of de novo mutations for all genes with >0 nonsynonymous de novo mutations.** P-values are provided for two enrichment tests, one with missense mutations restricted to PrimateAI-3D scores ≥ 0.821, and the other including all missense mutations.

**Table S6. Performance of PrimateAI-3D and 16 other pathogenicity classifiers on 9 saturation mutagenesis assays.** Absolute value of Spearman correlation of prediction scores of 17 classifiers (rows) with assay measurement across the 9 saturation mutagenesis assays (columns labelled by gene symbol) are provided.
